# A Compound that Inhibits Glycolysis in Prostate Cancer Controls Growth of Advanced Prostate Cancer

**DOI:** 10.1101/2023.07.01.547355

**Authors:** Takuma Uo, Kayode K. Ojo, Cynthia C. Sprenger, Kathryn Soriano Epilepsia, B. Gayani K. Perera, Mamatha Damodarasamy, Shihua Sun, Soojin Kim, Hannah H. Hogan, Matthew A. Hulverson, Ryan Choi, Grant R. Whitman, Lynn K. Barrett, Samantha A. Michaels, Linda H. Xu, Vicky L. Sun, Samuel L.M. Arnold, Haley J. Pang, Matthew M. Nguyen, Anna-Lena B.G. Vigil, Varun Kamat, Lucas B. Sullivan, Ian R. Sweet, Ram Vidadala, Dustin J. Maly, Wesley C. Van Voorhis, Stephen R. Plymate

## Abstract

**Purpose:** Metastatic castration-resistant prostate cancer remains incurable regardless of recent therapeutic advances. Prostate cancer tumors display highly glycolytic phenotypes as the cancer progresses. Non-specific inhibitors of glycolysis have not been utilized successfully for chemotherapy, because of their penchant to cause systemic toxicity. This study reports the preclinical activity, safety, and pharmacokinetics of a novel small molecule preclinical candidate, BKIDC-1553, with antiglycolytic activity.

**Experimental design:** We tested a large battery of prostate cancer cell lines for inhibition of cell proliferation, in vitro. Cell cycle, metabolic and enzymatic assays were used to demonstrate their mechanism of action. A human PDX model implanted in mice and a human organoid were studied for sensitivity to our BKIDC preclinical candidate. A battery of pharmacokinetic experiments, absorption, distribution, metabolism, and excretion experiments, and in vitro and in vivo toxicology experiments were carried out to assess readiness for clinical trials.

**Results:** We demonstrate a new class of small molecule inhibitors where antiglycolytic activity in prostate cancer cell lines is mediated through inhibition of hexokinase 2. These compounds display selective growth inhibition across multiple prostate cancer models. We describe a lead BKIDC-1553 that demonstrates promising activity in a preclinical xenograft model of advanced prostate cancer, equivalent to that of enzalutamide. BKIDC-1553 demonstrates safety and pharmacologic properties consistent with a compound that can be taken into human studies with expectations of a good safety margin and predicted dosing for efficacy.

**Conclusion:** This work supports testing BKIDC-1553 and its derivatives in clinical trials for patients with advanced prostate cancer.

## INTRODUCTION

Prostate cancer (PCa) is responsible for the deaths of more than 375,000 men worldwide and the second most common solid tumor in men(1). Although primary therapies such as surgery or radiotherapy appear to cure many men diagnosed with PCa, the disease becomes incurable once becomes metastatic(2, 3).

Treatment modalities for PCa have expanded significantly over the past decade with development of potent androgen receptor signaling inhibitors (ARSi) (e.g., abiraterone, enzalutamide)(2, 4). Additionally, tumors with loss of function mutations in homologous recombination repair pathway genes (e.g., BRCA1/2) define a subgroup of patients who can benefit from treatment with poly ADP ribose polymerase (PARP) inhibitors and possibly other DNA damaging therapies such as carboplatin(5). More recently, radionuclide therapies such as Radium 223 and PSMA targeting with Lutetium 177 have provided new avenues of therapy(6, 7). Although these therapies have been demonstrated to improve overall survival, the improvement is incremental, and resistance develops in the majority of cases(2, 8). One avenue of therapy for castration-resistant prostate cancer (CRPC) that has yet to realize its full potential involves the development of agents designed to selectively inhibit tumor metabolism(9, 10). To date, targeting tumor metabolism, particularly glycolysis or mitochondrial metabolism, has been stymied due to excessive toxicity and/or lack of efficacy(11).

Changes to cell metabolism is a hallmark of cancer that drives and supports oncogenesis(12–14). Both tumor genetics and tissue context define cancer dependence on specific metabolic pathways, with the tumor ultimately undergoing progression and being driven by combinations of oxidative phosphorylation, glycolysis, and potentially fatty acid metabolism(15). One hundred years have passed since Otto Warburg’s pioneering work to report the dominance of aerobic glycolysis in tumors(16). Nevertheless, non-specific antiglycolytic agents such as 2-deoxyglucose (2-DG), have never been clinically translated as they not only affect tumor cells but also inhibit normal proliferating cells(15, 17).

While glycolysis is not a significant driver in primary PCa, in more aggressive lethal forms of the disease glycolysis is a major driver of tumor progression especially since the tumor exists in a hypoxic environment(18–21). Challenging our ability to develop drugs against this pathway are multiple enzymatic isoforms in the glycolytic pathway that perform overlapping functions, complicating specific targeting(15). Accordingly, flexibility and redundancy among metabolic genes could shape the landscape of positive and negative impact on targeting tumor metabolism(9, 15). As far as glycolysis is concerned, isoform specific contribution to tumor progression is exemplified by pyruvate kinase M2 (PKM2) and hexokinase 2 (HK2), where activation and inhibition have been shown to reduce tumor progression, respectively, despite continued expression of corresponding isoforms such as PKM1 and HK1(22, 23). HK2 has been reported to be essential for PTEN/p53 deficient CRPC growth(24).

Previous protein kinase profiling of bumped kinase inhibitor (BKI) derived compounds (BKIDCs) (BKIDC-1369, 1553, 1561, and 1649) revealed potential human kinase targets, including Dual specificity mitogen-activated protein kinase kinase 2 (MEK2), human receptor-interacting serine/threonine-protein kinase 2 (RIPK2) and protein kinase D (PKD1-3)(25). Given that other studies have shown adenocarcinoma prostate cancer (adeno-PCa) cell lines are inhibited by pan-PKD inhibitors – irrespective of AR status – we set out to screen for cell types that might be affected by BKIDC compounds (26, 27). However, as we will show in this study, BKIDCs do not appear to act against PCa growth by inhibiting protein kinases.

In this study, we report on the discovery of a novel series of BKIDCs, that display selective inhibition of PCa growth and that are associated with a cessation of glycolysis immediately on exposure to the compounds. Furthermore, we describe a lead compound (referred to as BKIDC-1553) that demonstrates promising pharmacological and safety properties and activity in a mouse preclinical model of advanced PCa.

## MATERIALS AND METHODS

### Data availability

Derived data supporting the findings of this study are available upon request. Information and requests for resources and reagents should be directed to and will be fulfilled by lead contacts Stephen Plymate, Wesley Van Voorhis, and Takuma Uo. Synthesis protocols of all unique, non-FDA chemical compounds, synthesized for this publication are given below in the section **Synthesis of PP and PrP based BKIDCs.**

### Cell culture

LNCaP clone FGC (obtained in 2018. CRL-1740, RRID:CVCL_1379), C4-2 (obtained in November 2019. CRL-3314, RRID:CVCL_4782), C4-2B (obtained in November 2019. CRL-3315, RRID:CVCL_4784), VCaP (obtained in 2018. CRL-2876, RRID:CVCL_2235), DU145 (obtained in 2018. HTB-81, RRID:CVCL_0105), PC3 (obtained in 2018. CRL-1435, RRID:CVCL_0035), HepG2 (obtained in May 2013, HB-8065, RRID:CVCL_0027), HFF-1 (obtained in August 2020. SCRC-1041, RRID:CVCL_3285), and lymphoblastoid WIL2NS (obtained in April 2011, CRL-8155, RRID:CVCL_2761) were obtained from the American Type Culture Collection (ATCC). 293FT cell line was obtained in June 2016 from ThermoFisher Scientific (Invitrogen™, Cat# R70007, RRID:CVCL_6911). LAPC4 (RRID:CVCL_4744) was obtained from Dr. Charles Sawyers (Memorial Sloan Kettering Cancer Center, New York NY) through Dr. Peter Nelson (Fred Hutchinson Cancer Center, FHCC) in October 2019. U2OS cells (RRID:CVCL_0042) from Dr. Raymond Monnat (University of Washington) in October 2018, NCI-H660 in January 2020 (RRID:CVCL_1576) and LNCaP^APIPC^ from Dr. Peter Nelson in November 2011(28), LNCaP95 (obtained in 2012. RRID:CVCL_ZC87) from Dr. Alan Meeker (Johns Hopkins University), MSKCC EF1 derived from the organoid line MSKCC-CaP4(29) from John K. Lee (FHCC) in January 2020. LTL331R cells established from LTL331R tumor(29) were obtained from Dr. Yuzhuo Wang at Vancouver Prostate Centre (Vancouver BC, Canada) in March 2023.

Cells were maintained in RPMI 1640 (Gibco™, Cat# 21-870-092) (LNCaP, C4-2, C4-2B, DU145, PC3, MSKCC EF1, HepG2, WIL2NS, 293FT) supplemented with GlutaMax-I (Gibco™, Cat# 35050061), DMEM (Gibco™, Cat# 10569-010) (VCaP, U2OS, HFF-1), and IMDM (Iscove’s Modified Dulbecco’s Medium, Gibco™, Cat# 21056023) (LAPC4) supplemented with 5% fetal bovine serum (FBS) (Gibco™, Cat# 16000-044), 100 units/mL penicillin, and 0.1 mg/mL streptomycin (Gibco™, Cat# 15140122). LNCaP95 cells were grown in phenol-red-free RPMI medium (Gibco™, Cat# 11835030) supplemented with 5% charcoal stripped serum (Gibco™, Cat# 12676029). NCI-H660 was cultured in HITES medium (RPMI 1640 medium plus 5 μg /mL insulin, 10 μg/mL transferrin, 30 nM sodium selenite (ITS (Insulin-Transferrin-Selenium), Corning, Cat# 25-800-CR), 10 nM hydrocortisone (Sigma Aldrich, Cat# H0135), 10 nM β-estradiol (Sigma Aldrich, Cat# E2257)) with 10% FBS. The LTL331R cells were maintained in RPMI1640 (ATCC modification: Gibco™, Cat# A1049101) containing 5% FBS, 1% ITS (Gibco™, Cat# 41400-045), 10 nM hydrocortisone, 10 nM β-estradiol, 10 µM Y27632 (Tocris Bioscience, Cat# 1254), and 1% Cultrex Basement Membrane Extract, PathClear (R&D Systems, Cat# 3432-010).

All cell lines were routinely tested for mycoplasma contamination with Universal Mycoplasma Detection Kit (ATCC, Cat# 30-1012K ™) and used within 20 passages of receipt. The latest tests relevant to this study were done in December 2018 (VCaP, DU145, U2OS), January 2019 (293FT), May 2019 (LNCaP, LNCaP95, LNCaP^APIPC^, PC3), November 2019 (C4-2, C4-2B, LAPC4), February 2020 (NCI-H660, MSKCC EF1), September 2021 (HepG2), October 2021 (CRISPR-engineered LNCaP lines), October 2023 (HFF-1, WIL2NS), December 2023 (LTL331R).

Cell authentication was carried out with ATCC FTA Sample Collection Kit for Human Cell Authentication Service (ATCC 135-XV-20 ™). The latest test date was July 2023 for LNCaP (test specimen from cells collected in July 2021), LNCaP95 (July 2021), U2OS (January 2019), PC3 (January 2019), VCaP (August 2019), DU145 (December 2018), and HepG2 (June 2020) and October 2023 for WIL2-NS (July 2020) and NCI-H660 (February 2020).

Three cell lines were used for pharmacological characterization of BKIDC-1553. MDR1-MDCKII (RRID:CVCL_IZ19) was used sed to assess transport of BKIDC-1553 in both directions (apical to basolateral (A-B) and basolateral to apical (B-A)) across the cell monolayer and monitored for key markers of MDR1 transporter expression.

Mycoplasma was monitored with MycoAlert® PLUS Mycoplasma Detection Kit (Lonza, Cat# LT07-705) in April 2022 and the cell line was used in February 2023. The cell line was used on passage 9 from thaw. Caco-2 cells at passage 19 (RRID:CVCL_0025) were obtained from ATCC. Performance of the Caco-2 line in monolayer permeability experiments was characterized by comparison of results for reference compounds to published data. Cells were confirmed to be mycoplasma-free based on a luminescence-based enzymatic assay (MycoAlert®, Lonza, Cat# LT07-118) conducted within 1 week of the commencement and completion of the experiments. Cells used in the experiments were at passage 37 after thawing stocks at passage 30.

### Metabolomic flux assay

LNCaP95, LNCaP, and DU145 cells were plated at 500 K cells per well on 6-well plates (Thermo Scientific™, Cat# 140675) coated with 10 µg/mL poly-D-lysine (PDL) (Sigma Aldrich, Cat# A-003 M), overnight in media containing unlabeled 10 mM D-glucose and 2 mM L-glutamine. Cells were treated in triplicate with drug or vehicle for 30 min. Cells were then quickly washed two times with RPMI media with L-glutamine and without glucose (Gibco™, Cat# 11879020). Media was then changed to fresh RPMI with 10 mM [U]^13^C-glucose (Cambridge Isotope Laboratories, CLM 13961369) and unlabeled 2 mM glutamine in the presence of different concentrations of BKIDC-1553-N-Me. DMSO (Sigma-Aldrich, Cat# D5879) was used as vehicle control. Cells were then harvested 30 min after treatment with [U]^13^C-glucose and inhibitors. Cells were then quickly washed twice with ice cold blood bank saline and quenched with 500 µL HPLC-grade 80% methanol (Fisher Chemical, Cat# A4561) containing 2.5 µM valine D8 extraction standard. Samples were then centrifuged at 4°C at 17,000 g for 5 min. 400 µL of supernatant was transferred to a new microfuge tube and centrifuged at 4°C at 17,000 g for 5 min. 350 µL of supernatant was then transferred to a new microfuge tube and dried down with a Labonco Centrivap Centrifugal Vacuum Concentrator at 4°C and placed at −80°C until time of analysis. Samples were then re-constituted with 50 µL of 80% HPLC-grade methanol, vortexed for 10 min at 4°C, and centrifuged at 17,000g for 5 min at 4°C. 40 µL of supernatant was transferred to a liquid chromatography-mass spectrometry (LC-MS) vial for measurement by LC-MS. Metabolite quantitation was performed using a Q Exactive HF-X Hybrid Quadrupole-Orbitrap Mass Spectrometer equipped with an Ion Max API source and H-ESI II probe, coupled to a Vanquish Flex Binary UHPLC system (Thermo Scientific). Mass calibrations were completed at a minimum of every 5 days in both the positive and negative polarity modes using LTQ Velos ESI Calibration Solution (Pierce). Samples were chromatographically separated by injecting a sample volume of 1 μL into a SeQuant ZIC-pHILIC Polymeric column (150 x 2.1 mm 5 µM, EMD Millipore). The flow rate was set to 150 µL/min, autosampler temperature set to 10°C, and column temperature set to 30°C. Mobile Phase A consisted of 20 mM ammonium carbonate and 0.1% (v/v) ammonium hydroxide, while Mobile Phase B consisted of 100% acetonitrile (ACN). The sample was gradient eluted (%B) from the column as follows: 0-20 min: linear gradient from 85% to 20% B; 20-24 min: hold at 20% B; 24-24.5 min: linear gradient from 20% to 85% B; 24.5 min-end: hold at 85% B until equilibrated with ten column volumes. Mobile Phase was directed into the ion source with the following parameters: sheath gas = 45, auxiliary gas = 15, sweep gas = 2, spray voltage = 2.9 kV in the negative mode or 3.5 kV in the positive mode, capillary temperature = 300°C, RF level = 40%, auxiliary gas heater temperature = 325°C. Mass detection was conducted with a resolution of 240,000 in full scan mode, with an AGC target of 3,000,000 and maximum injection time of 250 msec. Metabolites were detected over mass range of 70-1050 m/z in full scan mode with polarity switching. Quantitation of all metabolites was performed using Tracefinder 4.1 (Thermo Fisher Scientific RRID:SCR_023045) referencing an in-house glycolytic metabolite standards library using ≤ 5 ppm mass error.

### Glucose uptake assay

#### Luciferase-based assay

LNCaP cells were plated at 240 K cells/well in PDL-coated 24-well plates (Corning, Cat# 353226). 24 h later, medium was changed to serum and glucose-free medium containing BKIDC at 20 µM or cytochalasin B (CytB, C6762 Sigma-Aldrich, Cat# C6762) at 50 µM for 30 min prior to the experiment. Cells were then rinsed and incubated media containing 2-DG and BKIDC or CytB for 10 min. Glucose uptake was measured using the Glucose Uptake-Glo™ Assay (Promega, Cat # J1341) according to the manufacturer’s protocol. This luciferase-based assay measures 2-DG-6-phosphate (2-DG-6-P) which accumulates as a dead-end product after hexokinase-mediated conversion of 2-DG. Luminescence signals were measured on 96-well plate (Corning, Cat# 3915) on Promega GloMax® plate reader (RRID:SCR_015575). Either BKIDC-1553-N-Me or CytB was added after cellular harvest, to demonstrate these compounds did not interfere with the assay to evaluate 2-DG accumulation. The data were presented as mean relative luminescence units (RLU) ± standard deviation (SD) (n=3).

#### Radiometric assay

Uptake of ^3^H-3-O-methylglucose (American Radiolabeled Chemicals, St. Louis, MO) and ^3^H-2-deoxyglucose (Perkin Elmer, Waltham, MA) was measured as carried out previously(30, 31). Cells were incubated in 12 x 75 mm polypropylene test tubes containing Krebs-Ringer bicarbonate solution (KRB) (with 5 mM bicarbonate and 0.1% BSA) at 37°C for 30 min in a shaking water bath. Cell densities were 2 x10^6^ cells per 200 μL for all cell types. Radiolabeled molecules (0.175 μCi) were added to the tubes using an Eppendorf Repeater pipet and the cells were further incubated in the presence of a radiotracer for 30 min. (Experiments demonstrated that between 10 and 30 min, uptake of tracer was linear). The specific activities and the concentrations in the incubation media of the radiotracers were as follows: [^3^H]3-O-MG, 90 Ci/mmol, 24 nM; [^3^H]2-deoxyglucose, 20 Ci/mmol, 119 nM. Accumulation of radiolabel was determined by separating the cell-associated radioactivity (CAR) from the free radioactivity by transferring the cell suspension to a 0.4 mL centrifuge tube (USA Scientific, Ocala, FL) containing a layer of oil consisting of 1:37.5 n-dodecane:bromo-dodecane (Sigma–Aldrich, St. Louis, MO) and spinning at 12,535 g for 8 s (Beckman E centrifuge, Beckman Coulter, Fullerton, CA). The tube was placed briefly in liquid nitrogen, cut through the radioactive-free oil layer with a razor blade, and the bottom portion of the tube containing the cell pellet was placed into a 7 mL glass scintillation vial. The ^3^H labels were counted using a Beckman liquid scintillation counter (Model LS6500) after adding 5 mL of Ecolume liquid scintillation cocktail (MP Biomedicals, Cat# 0188247001) and vortexing the vials. Values of non-specific association of radioactivity was determined by performing the assay in the presence of 5 mM glucose at 4°C. The CAR data for each sample was normalized by subtracting off the non-specific values, and then dividing by the cell number and the dose. Quench correction for samples was not needed since counts were normalized to the dose and quenching did not vary from sample to sample.

### In vitro AMPK activity measurement with total cell lysates

LNCaP95 cells were grown on 24-well plate for 24 h and treated with BKIDC at 20 µM for 1 h. AMPK activity in total cell lysates was measured using CycLex AMPK Kinase Assay kit (MBL International Corporation, Cat# CY-1182) according to the manufacturer’s instructions. Briefly, total cell lysates were prepared in lysis buffer (20 mM Tris-HCl, pH 7.5, 250 mM NaCl, 10% glycerol, 0.5% NP-40, 1 mM DTT) supplemented with Halt Protease Inhibitor Cocktail and Halt Phosphatase Cocktail (Thermo Scientific™, Cat# 78440) and incubated for 30 min at 30°C in a pre-coated plate with an AMPK substrate peptide corresponding to mouse insulin receptor substrate-1 (IRS-1)(supplied as part of kit). AMPK activity was measured by monitoring the phosphorylation of Ser-789 in IRS-1 using an anti-mouse phospho-Ser-789 IRS-1 monoclonal antibody (AS-4C4: supplied as part of kit) and peroxidase-coupled anti-mouse IgG antibody. Conversion of the chromogenic substrate tetramethylbenzidine was quantified by absorbance measurement at 450 nm in BioRad iMark™ Microplate Absorbance Reader (RRID:SCR_023799).

### Real-time cellular bioenergetic assay

We used Agilent XFe24 analyzer (RRID:SCR_019539) to measure extracellular acidification rate (ECAR) and oxygen consumption rate (OCR) which reflect the rates of glycolysis and oxidative phosphorylation, respectively. Cells were plated at 40 K cells/well in Seahorse XF24 V7 PS Cell Culture Microplates (Agilent, Cat# 10777-004) coated with 10 µg/mL PDL and incubated overnight. The culture medium was then replaced by Seahorse XF RPMI medium (Agilent, Cat# 103576-100) and cells were pre-incubated in a CO_2_-free incubator at 37°C for 1 h prior to metabolic analysis. The effect of BKIDC was evaluated by pre-treating cells with 20 µM BKIDC during 1h-pre-incubation period with and without glucose prior to mitochondrial and glycolytic stress tests, respectively. Mitochondrial stress test was carried out by sequentially injecting oligomycin (Sigma, Cat# O4876), carbonyl cyanide 4-(trifluoromethoxy)phenylhydrazone (FCCP: Sigma, Cat# C2920), and rotenone (Sigma, Cat# R8875)/antimycin A (Sigma, Cat# A8674). Glycolytic stress test was done by sequential injection of glucose (Gibco™, Cat# A2494001), oligomycin, and 2-deoxy-D-glucose (2-DG: Sigma, Cat# D8375). The acute effect of BKIDC was assessed by directly adding BKIDC through the injection port. The bioenergetic profiles were analyzed from at least 3 biological replicates. Results were automatically generated and analyzed by Seahorse Wave Desktop Software, which were exported as files formatted for GraphPad Prism (RRID:SCR_002798).

### Measurement of intracellular ATP levels

50 K cells were plated on 48-plate (Corning, Cat# 353078) and grown overnight. Cells were treated with the indicated compounds for 2 h unless specified. Metabolic inhibitors used include: oligomycin, antimycin A, FCCP, rotenone, 2-DG, and sodium oxamate (Cayman Chemical, Cat# 565-73-1). After removing culture media, 80 μL of the CellTiter-Glo 2.0 Cell Viability Assay kit (Promega, Cat# G9241) was added to the cells. Intracellular ATP levels under each condition were assessed on 96-well plate (Corning, Cat# 3915) on GloMax® plate reader.

### CRISPR/Cas9-mediated genome editing

We used a CRISPR/Cas9 approach to establish a stable knockout LNCaP cell lines deficient of either HK1 or HK2(32). sgRNA protospacer of HK1 (HK1-1: GGTGAGGTGGAAATGAGCCA; HK1-3: GGAGGGCAGCATCTTAACCA) and HK2 (HK2-1: GATGCGCCACATCGACATGG; HK2-2: TAAGCGGTTCCGCAAGGAGA) was cloned into lentiCRISPR v2 construct (RRID:Addgene_52961) using a protocol available online (Zhang_lab_LentiCRISPR_library_protocol.pdf). gRNA sequence (control: GTATTACTGATATTGGTGGG) was used to prepare the lentiviral construct for a non-targeting control(33). The oligonucleotides were commercially synthesized at Eurofins Scientific. 293FT cells were seeded on 6-well plates coated with poly-D-lysine. The following day, 293T cells were co-transfected with lentiCRISPR v2 construct and the second-generation lentiviral packaging plasmids pMD2.G for VSV-G (RRID:Addgene_12259) and pCMVR8.74 for GAG, POL, TAT, and REV (RRID:Addgene_22036) using Lipofectamine 3000 reagent (Invitrogen™, Cat# L3000015). Lentivirus-containing supernatant was collected at 12 and 24 h after transfection, filtered through a 0.45-μm filter (Cytiva, Cat# WHA-9916-1304), and used to infect LNCaP as a recipient cell line, overnight in the presence of 5 µg/mL polybrene (Sigma, Cat# 107689). The infected cells were selected with 5 µg/mL of puromycin (Gibco™, A1113803) for 3 days. Selected cells were immediately used as pool cell populations to evaluate glycolytic capacities in relation to BKIDC. On the other hand, to isolate morphologically single colonies, they were also plated at the density of 0.5 cell/well in 96-well plates in the conditioned media from native LNCaP line. Resulting colonies were screened by Western blot analysis for loss of either HK1 or HK2 expression. After a second confirmatory Western blot analysis, the genomic junctions flanking corresponding sgRNA binding sites were amplified with sequence-specific primers by PCR using Platinum Taq DNA Polymerase and the amplicons were cloned into pCR™2.1-TOPO™ plasmid with TOPO™ TA Cloning™ Kit (Invitrogen™, Cat# 450641) and Sanger-sequenced using the T7 promoter primer (TAATACGACTCACTATAGGG) and M13 Reverse primer (CAGGAAACAGCTATGAC) at GENEWIZ (South Plainfield, NJ). Particular isolates (HK11-9, HK13-5, HK21-5, HK22-1) were chosen for further experiments. Pool cell population which was infected with non-targeting control virus served as control cell lines for the experiments with both clonal cell lines and pool cell populations.

### Cell proliferation assay

Cell proliferation was determined by MTS [3-(4,5-dimethylthiazol-2-yl)-5-(3-carboxymethoxyphenyl)-2-(4-sulfophenyl)-2H-tetrazolium, inner salt] assay using CellTiter 96® AQueous Non-Radioactive Cell Proliferation Assay (MTS) kit (Promega, Cat# G3582) at 72 h after BKIDCs were added for all cell lines, except for VCaP at 120 h. Each cell line was tested in 96 well plates at 5 K cells/well in the culture media. The data shown are representative of at least two independent biological experiments, normalized to the values obtained from cells grown in medium containing 0.1% DMSO vehicle and presented as mean ± standard deviation (SD) (n=3-5).

### CellTiter-Glo® 3D Cell Viability Assay

LTL331R cells were plated on 96-well plate (Thermo Scientific, Cat# 167425) at 15 K cells/well. At 2 days after plating, cells were treated with either BKIDC-1553 (1.25-20 µM) or enzalutamide (1.25-20 µM) using DMSO as vehicle control. At 72 h after treatment, cell proliferation was determined using CellTiter-Glo™ 3D Cell Viability Assay (Promega, Cat# G9683.) on GloMax® plate reader. The data shown are representative of at least two independent biological experiments, normalized to the values obtained from cells grown in medium containing 0.1% DMSO vehicle and presented as mean ± standard deviation (SD) (n=3-4).

### Crystal Violet Assay

Cell proliferation of LNCaP95 and PC3 was assessed by crystal violet assay at 72 h after BKIDCs were added. Each cell line was tested in 96 well plates (Thermo Scientific™, Cat# 167425) at 5 K cells/well in the culture media. Cells were fixed with 5% glutaraldehyde (Sigma-Aldrich, Cat# G6257) for 5 min and stained with 0.5% crystal violet (Sigma-Aldrich, Cat# C3886) solution in 20% methanol (VWR Chemicals BDH®, Cat# BDH1135) for 30 min followed by rinsing with distilled water 3 times. After completely air-drying the plate overnight, 50 μL of 10% acetic acid was added to each well to dissolve the cell-bound crystal violet. 10 min later, absorption was determined at 590 nm in BioRad iMark™ Microplate Absorbance Reader (RRID:SCR_023799). The data shown are representative of at least three independent biological experiments, normalized to the values obtained from cells grown in medium containing 0.1% DMSO vehicle and presented as mean ± standard deviation (SD) (n=3-5).

### Cell Cycle Analysis

Analysis was performed as previously described(34). Cells treated with BKIDCs for 30, 48, or 72 h were collected and resuspended in a solution of 10 µg/mL 4,6-diamidino-2-phenylindole (DAPI) and 0.1% Nonidet P-40 detergent in a Tris buffered saline. The suspension was analyzed on the LSR II flow cytometer (RRID:SCR_002159, BD biosciences), with ultraviolet or violet excitation and DAPI emission collected at >450 nm. DNA content and cell cycle were analyzed as previously described using the software program MultiCycle(34)(Phoenix Flow Systems, San Diego, CA).

### Western blot analysis

Cells were lysed in M-PER Mammalian Protein Extraction Reagent (Thermo Scientific™, Cat# 78501) supplemented with Protease Inhibitor Cocktail and Halt Phosphatase Cocktail (Thermo Scientific™, Cat# 78440). Protein measurement was determined using Pierce™ BCA Protein Assay Kits BCA assay (Thermo Scientific™, Cat# 23225) using Pierce™ Bovine Serum Albumin Standard Pre-Diluted Set (Thermo Scientific™, Cat# 23208). Samples were run on 4–15% Mini-PROTEAN® TGX™ Precast Protein Gels (Bio-Rad, Cat# 4561084) and proteins transferred to Immun-Blot® PVDF Membrane (Bio-Rad, Cat# 1620177) using Bio-Rad TransBlot Turbo transfer system (RRID:SCR_023156). Tris-buffered saline (TBS: 20 mM Tris-HCl, pH 7.5, containing 0.15 M NaCl) with 0.3% Tween 20 (Fisher Scientific, Cat# BP337-100)(TBS-T) was used. Membranes were blocked with the blocking buffer (TBS-T with 5% nonfat milk (LabScientific, Cat# M-0842). Blots were incubated with appropriate antibodies in the blocking buffer. TBS-T was used for all washes. Chemiluminescence with Immobilon Forte Western HRP substrate (Millipore Sigma, Cat# WBLUF0500) and near-Infrared fluorescence imaging were performed with FluorChem R (ProteinSimple) and Odyssey^®^ Imaging System (Li-COR Biosciences), respectively. The following antibodies were used for Western blots with the indicated dilution factors: Hexokinase I (C35C4) Rabbit mAb (Cell Signaling Technology, Cat# 2024, RRID:AB_2116996) at 1:1,000, HK2 Monoclonal Antibody (3D3) (Invitrogen™, Cat# MA5-15679, RRID:AB_10986812) at 1:1,000, PARP Antibody (Cell Signaling Technology, Cat# 9542, RRID:AB_2160739) at 1:2,000, Acetyl-CoA Carboxylase (C83B10) Rabbit mAb (Cell Signaling Technology, Cat# 3676, RRID:AB_2219397) at 1:1,000, Phospho-Acetyl-CoA Carboxylase (Ser79) (D7D11) Rabbit mAb (Cell Signaling Technology, Cat# 11818, RRID:AB_2687505) at 1:500, Rabbit Anti-GAPDH Monoclonal Antibody Unconjugated, Clone 14C10 (Cell Signaling Technology, Cat# 2118, RRID:AB_561053) at 1:5000, S6 Ribosomal Protein (54D2) Mouse mAb (Cell Signaling Technology, Cat# 2317, RRID:AB_2238583) at 1:2000, Phospho-S6 Ribosomal Protein (Ser235/236) (D57.2.2E) XP Rabbit mAb (Cell Signaling Technology, Cat# 4858, RRID:AB_916156) at 1:2000, Anti-β-Actin antibody-Mouse monoclonal (Sigma-Aldrich, Cat# A1978, RRID:AB_476692) at 1:5,000. Secondary antibodies used for chemiluminescence detection were HRP conjugated anti-mouse antibody (Cytiva, Cat# NA931) and anti-rabbit antibody (Vector Laboratories, Cat# PI10001). Near-infrared detection was carried out with IRDye® 680RD Donkey anti-Rabbit IgG (H + L) (LI-COR Biosciences, Cat# P/N 925-68073) and RDye® 800CW Donkey anti-Mouse IgG (H + L) (LI-COR Biosciences, Cat# P/N 925-32212). Western blot Images were processed with GIMP (GNU Image Manipulation Program, Version 2.10).

### In vivo BKIDC-1553 studies

Male severe combined immunodeficient (SCID) mice were used for all in vivo experiments (Charles River Laboratories, strain code# 236). PC3 (100,000 cells) were mixed 1:1 with Matrigel (Corning, Cat# 356234) and injected subcutaneously into rear flanks of intact SCID mice (10 mice were enrolled per treatment group for PC3 study). When tumors reached ∼100 mm^3^, either vehicle (7% Tween-80, 3% EtOH in pharmaceutical grade PBS) or 20 mg/kg of BKIDC-1553 was given by oral gavage three times weekly (Mon, Wed, Fri), for up to 4 weeks. A patient-derived PCa xenograft, LuCaP 35 (RRID:CVCL_4853), was implanted subcutaneously into rear flanks of intact SCID mice (The number of enrolled mice was: 8 for control, 9 for 1553, 7 for enzalutamide). When tumors reached ∼100 mm^3^, mice were given vehicle (7% Tween-80, 3% EtOH in pharmaceutical grade PBS), 20 mg/kg of BKIDC-1553, or 10 mg/kg enzalutamide (Medivation, log# 083112-02, lot: BREC-0035-123T) by oral gavage three times weekly (Mon, Wed, Fri) for up to 5 weeks. All mice were assigned to treatment or control groups using a random number generator. ARRIVE guidelines were followed for all animal experiments. All animal studies were approved by the University of Washington Institutional Animal Care and Use Committee (protocol 2797-04).

### Determining BKIDC-1553 concentrations in tumors

Each tumor was flushed with Dulbecco’s phosphate-buffered saline (DPBS), weighed and placed in storage at −80°C. On the day of homogenization, 0.9% NaCl was added to each sample to achieve 100 mg/mL of tissue concentration. Samples were then homogenized using a handheld motorized homogenizer (Thermo Fisher Scientific) on ice, and 100 μL of each homogenate sample was placed in a 96-well plate. In addition, tissues from vehicle control-treated mice were homogenized and placed in a 96-well plate to be used as blanks or standards. Calibration curves were generated individually for each type of tumor. An internal standard was added to each sample to a final concentration of 20 nM. All homogenates were stored at −80°C. On the day of tissue extraction, 10 μL of each sample were added to a 96-well plate that contained 90 μL of 80% ACN:20% water. The samples were mixed and centrifuged at 4,200 g for 20 min. 25 μL of the supernatants were placed in a 96-well plate that contained 475 μL of 80:20 ACN:water. Then, the samples were mixed and centrifuged at 4,200 g for 10 min. Supernatant from each sample was placed in a new 96-well plate, sealed and analyzed with an Acquity UPLC in tandem with a Xevo TQ-S micro mass spectrometer (Waters Corp). Sample concentrations were determined using the internal standard normalized calibration curves. To report tissue concentrations of BKIDC-1553, diluted homogenates were adjusted post-integration on a weight:weight basis. Individual tumor concentrations, means, and standard deviations are reported in **Supplementary Table 5**.

### Pharmacology (PK), Absorption, Distribution, Metabolism, and Excretion (ADME), and Toxicology Methods

PK, ADME and Toxicology methods with BKIDC-1553 were carried out as described previously where the compound was referred to as Compound **32** (35) and reanalysis of PK data was carried out with Phoenix WinNonlin software (Certara, NJ). A human dose prediction for BKIDC-1553 was generated using estimated human pharmacokinetic parameters and BKIDC-1553 concentrations associated with efficacy in a preclinical prostate cancer mouse model. BKIDC-1553 clearance (Cl) and volume of distribution (V) in a 74 kg human were estimated by allometric scaling using pharmacokinetic parameters observed with IV dosing in rats, dogs, and non-human primates (Table S7)(36–38). Since the exponent for Cl using body weight was >1, the product of Cl x brain weight was used(39). Based on the rapid and nearly complete absorption observed in preclinical models, BKIDC-1553 was assumed to have an oral bioavailability of 1.0 in humans. The BKIDC-1553 concentration associated with efficacy in vivo was determined using a population pharmacokinetic modeling approach. Briefly, observed BKIDC-1553 plasma concentrations in mice were fit with a 1-compartment structural model and a proportional error model. BKIDC-1553 concentrations were simulated based on post-hoc estimates of pharmacokinetic parameters (absorption rate constant (ka), Cl, V) for each mouse (n = 3). The simulated concentrations were used to estimate the maximum BKIDC-1553 plasma concentration (C_max_) and area under the curve (AUC) over the last day of dosing (AUC_264-288h_) and the entire dosing period (AUC_0-288h_).

PK experiments were performed as previously described(35) with the following modifications. Male SCID mice were given an oral dose of 20 mg/kg BKIDC-1553 (in 7% Tween-80, 3% EtOH in pharmaceutical grade PBS vehicle) on Monday. Wednesday, and Friday for two weeks for a total of 6 doses. Blood was sampled after the 1^st^, 4^th^, and 6^th^ doses, with a terminal bleed taken on the third Monday, 72 h after the final dose. Blood samples at each dose were taken pre-dose and at 1 h, 2 h, 4 h, 6.5 h, and 24 h post dose. Plasma protein determination was performed using the dialysis membrane technique as previously described(40). Caco-2 permeability testing was performed in Hanks balanced salt solution containing 20 mM HEPES using 0.3 cm^2^ polycarbonate filter transwells seeded with 6 x 10^4^ cells. On day 24 post-seeding, donor solutions containing test compounds in DMSO (0.1% final) were added to confluent cell monolayers and compound flux was assessed from the acceptor chamber at 5-6 time points over 120 min. Acceptor chamber volume was replaced with blank buffer and dilution corrected with each sample collection. Samples were stored frozen at −80°C until analysis by LC-MS. Apparent permeability coefficient (P_app_) calculations were determined using the equation:

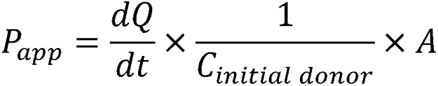

Where dQ/dt = apparent steady-state transport rate (µmol/s); A = monolayer surface area; and C_initial donor_ = concentration in donor at experiment initiation. Apparent efflux ratios were calculated based on the P_app_ values in the A-B and B-A directions.

Additional permeability testing in MDR1-MDCKII cells (RRID:CVCL_IZ19) was conducted by Eurofins CRO services (Eurofins Scientific, Luxembourg) using 10 μM BKIDC-1553. Hepatic microsome metabolism of BKIDC-1553 was performed with cryopreserved liver microsomes from dogs, humans, and rats. Microsome incubations were prepared by pre-incubating 1.0 mg/mL cryopreserved liver microsomes in phosphate buffer (pH 7.4, 100 mM) for 5 min at 37_C. The reaction was started with the addition of UDPGA (5 mM), NADPH (2 mM) and incubation was performed for 1 h. The final concentration of test article was 10 µM with <0.5% organic. The sample was quenched with an equal volume of acetonitrile, vortexed, centrifuged at 13,000 rpm for 5 min, the supernatant (170 µL) was transferred to 96 well plate and partially dried under nitrogen to ∼ 120 µL for ultra-high-performance liquid chromatography-high resolution mass spectrometry (UPLC-HRMS, with UPLC instrument: Waters ACQUITY I-Class system, and Mass Spectrometer instrument: Waters Xevo G2 XS) analysis. The positive control was a separate incubation of verapamil using the same microsomes to confirm the viability of the incubation. The negative controls were separate incubation of the test article using the incubation media and same microsome incubation without NADPH and UDPGA. The LC/MS full scan data was processed manually, assisted by Metabolynx, to identify the metabolites. Product ion spectra were acquired by a separate UPLC-HRMS/MS run to assign the structures of the metabolites by MS/MS data interpretation. In vitro inhibition of Cytochrome P450 (CYP) enzymes was performed in five major isoforms (CYPs 1A2, 2C9, 2C19, 2D6, and 3A4) in human liver microsomes using up to 20 μM BKIDC-1553 and included standard CYP specific inhibitors (furafylline, sulfaphenazole, ticlopidine, quinidine, and ketoconazole) and substrates (phenacetin, tolbutamide, *(S)*-mephenytoin, dextromethorphan, and midazolam/testosterone). Various concentration of BKIDC-1553 and positive control inhibitors were co-incubated at 37°C with each substrate in human liver microsomes (XenoTech LLC, Lenexa KS) for 10-40 min, with reactions initiated by the addition of an NADPH-regenerating system and quenched by addition of ice-cold acetonitrile containing 0.15 µg/mL of diazepam internal standard. Quenched samples were analyzed by LC-MS (Waters/Micromass Xevo TQD triple-quadrupole) and concentrations determined relative to standards in quenched microsomal matrix. Inhibitory activities were based on the reduction in formation of the specific CYP-mediated metabolite relative to no-inhibitor controls and reported as 50% inhibition values (IC_50_). Additional CYP inhibition studies were conducted by Eurofins CRO services on CYPs 2B6 and 2C8 with 10 μM BKIDC-1553 using Clopidogrel and Montelukast as inhibitors and bupropion and amodiaquine as substrates for 2B6 and 2C8, respectively. The mutagenic toxicity of BKIDC-1553 was assessed by measuring its ability to induce reverse mutations in Salmonella TA98 (frame shift) and TA100 (base-pair) strains in the presence and absence of a metabolizing Aroclor 1254-induced rat S9 mix with the top concentration of 180 µg/well, in the modified 24-well assay, which is equivalent to 5000 µg/plate in the standard plate assay. A positive response is defined with a 2-fold increase in revertant colonies(41). The genotoxicity of BKIDC-1553 was assessed in an in vitro micronucleus test carried out with Chinese hamster V79 cells and determined by measuring the extent of micronucleus formation with and without metabolizing Aroclor 1254-induced rat S9 mixture. BKIDC-1553 was tested at a top concentration of 500 µg/mL. The safety 44 screen assay was performed by Eurofins as described (https://cdnmedia.eurofins.com/corporate-eurofins/media/1069358/safetyscreen44_epdsfl420june16.pdf)(42).

Safety testing of BKIDC-1553 in mice, using oral gavage of BKIDC-1553 suspended in a lipid-based vehicle (60% Phosal 53 MCT: 30% PEG-400: 10% ethanol), was performed with 35 mg/kg, 70 mg/kg, and 100 mg/kg BKIDC-1553 once daily oral doses for 7 days. Mice were monitored daily for signs of toxicity and glucose levels were sampled from tail snip using an Accu-Chek Guide Me glucometer (Roche) at pre-dose, 2, and 4 h post dose time points. Blood samples were collected after the 3^rd^ dose and processed for LC-MS/MS (Waters Acquity Xevo TQ-S micro) analysis of BKIDC-1553 levels in plasma.

Non-GLP toxicology studies were carried out in dogs by StillMeadow Inc., (Sugarland, Texas). In the first experiment, four groups of one male and one female >9-month-old beagle dogs each were dosed orally by gavage tubes with either the vehicle or BKIDC-1553 daily for 14 days. Group I was a Vehicle Control (80% Phosal 53 MCT, 15% PEG 400 and 5% Ethanol) at 1 mL/kg and Groups II, III and IV received 1.6 mg/kg, 5 mg/kg and 15 mg/kg of BKIDC-1553 in the same vehicle at 1 mL/kg, respectively. Dogs were monitored during the entire time of the experiment for changes in body weight and food consumption and for acute signs of toxicity and general health. Heparinized plasma for BKIDC-1553 quantitation was collected pre-dose, and 2, 4, 6, 24, and 48 h after the last dose, and measured by LC-MS/MS. Dogs were monitored for blood glucose levels (AlphaTRAK glucometer, Zoetis) prior to and 2 h after the first dose of the day, each day of dosing. Blood and urine samples were also collected for serum chemistry, hematology, and coagulation and urinalysis pre-dose and 24 h after the last dose. These samples were analyzed by Antech Diagnostics. In the second non-GLP dog toxicology experiment, a male and female >9-month-old beagle dog was first treated for 5 days with oral gavage twice a day (6 h apart) of the 4 mL/kg lipidic vehicle (80% Phosal 53 MCT, 15% PEG 400 and 5% Ethanol), blood and urine were collected at the 24 hours after dosing, tested as above. After a 4-week washout period, the same dogs were treated by oral gavage with BKIDC-1553 at 7.5 mg/mL in 4 mL/kg lipidic vehicle at 6 h intervals (30 mg/kg twice a day) for two days. Vomiting, decrease activity, and salivation was noted in both animals on day two of BKIDC-1553 administrations, and the next day, day 3, the BKIDC-1553 (7.5 mg/mL)-vehicle volume was reduced to 2 mL/kg administered twice daily at 6 h intervals (15 mg/kg twice a day) for an additional 3 days. Blood glucose was monitored by glucometer measurements before and 2 h after administration of each dose. Plasma for PK measurements were collected as noted above, after the last dose on the fifth day and the plasma BKIDC-1553 levels determined via LC-MS/MS.

Rat cardiovascular (CV) safety studies were carried out at AbbVie Inc as previously described(43).

### Synthesis of PP and PrP based BKIDCs

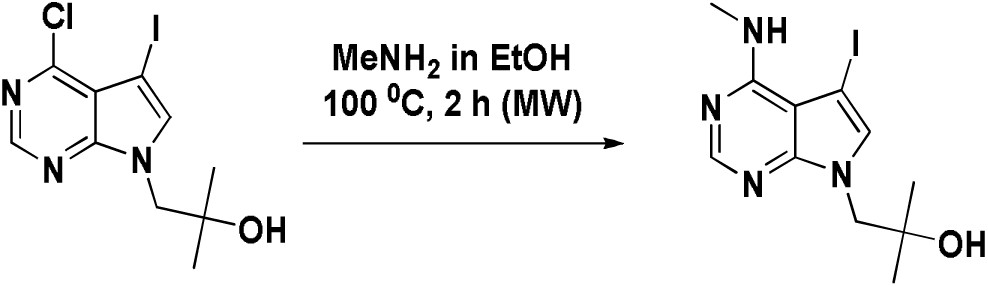

1-(4-chloro-5-iodo-7H-pyrrolo[2,3-d]pyrimidin-7-yl)-2-methylpropan-2-ol (0.6 g, 1 eq) was heated to 100 °C for 2 h in a microwave with a solution of 33% methylamine in ethanol (5.7 mL, keeping the concentration of the pyrrolopyrimidine scaffold around 0.3 M in solution). The crude mixture was concentrated under reduced pressure and purified by flash chromatography using 5% MeOH/DCM to obtain 1-(5-iodo-4-(methylamino)-7H-pyrrolo[2,3-d]pyrimidin-7-yl)-2-methylpropan-2-ol with a 64% yield. ^1^H NMR (500 MHz, CDCl_3_) δ 8.18 (s, 1H), 7.30 (s, 1H), 4.12 (s, 2H), 3.11 (s, 3H), 1.14 (s, 6H). Calcd for C_11_H_15_IN_4_O 346.03, found (M+H)^+^ 347.1.

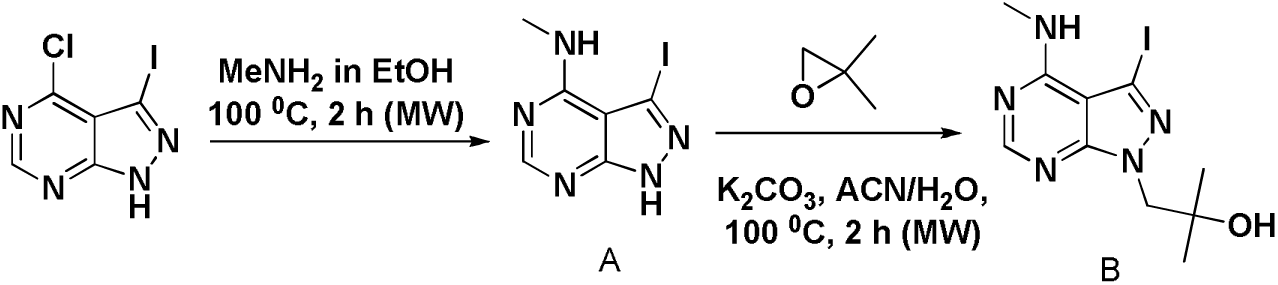

A mixture of 4-chloro-3-iodo-1H-pyrazolo[3,4-d]pyrimidine (125 mg, 0.354 mmol) and 33% methyl amine in ethanol (1.2 mL) was heated to 100 °C for 2 h in a microwave reactor. The crude mixture was concentrated under reduced pressure. Crude A was obtained in 62% yield and was used in the next reaction without further purification. ^1^H NMR (500 MHz, MeOD) δ 8.16 (s, 1H), 5.39 (s, 1H), 3.04 (d, *J* = 9.8 Hz, 3H).

Compound A (60 mg, 0.22 mmol) was heated to 150 °C for 3 h with 2,2-dimethyloxirane (40 uL, 2.0 eq) and K_2_CO_3_ (61 mg, 2.0 eq) in a mixture of 8.5:1.5 ACN/water [A=0.64M]. Crude mixture was concentrated under reduced pressure and purified by flash chromatography using 5% MeOH/DCM to obtain compound B with a 57% yield. ^1^H NMR (500 MHz, MeOD) δ 8.28 (s, 1H), 4.32 (s, 2H), 4.30 (s, 1H), 3.16 (s, 3H), 1.23 (s, 6H). Calcd for C_10_H_14_IN_5_O 347.02, found (M+H)^+^ 348.3.

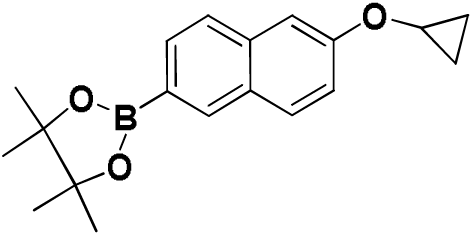

2-(6-cyclopropoxynaphthalen-2-yl)-4,4,5,5-tetramethyl-1,3,2-dioxaborolane was synthesized according to previously published protocols(35).

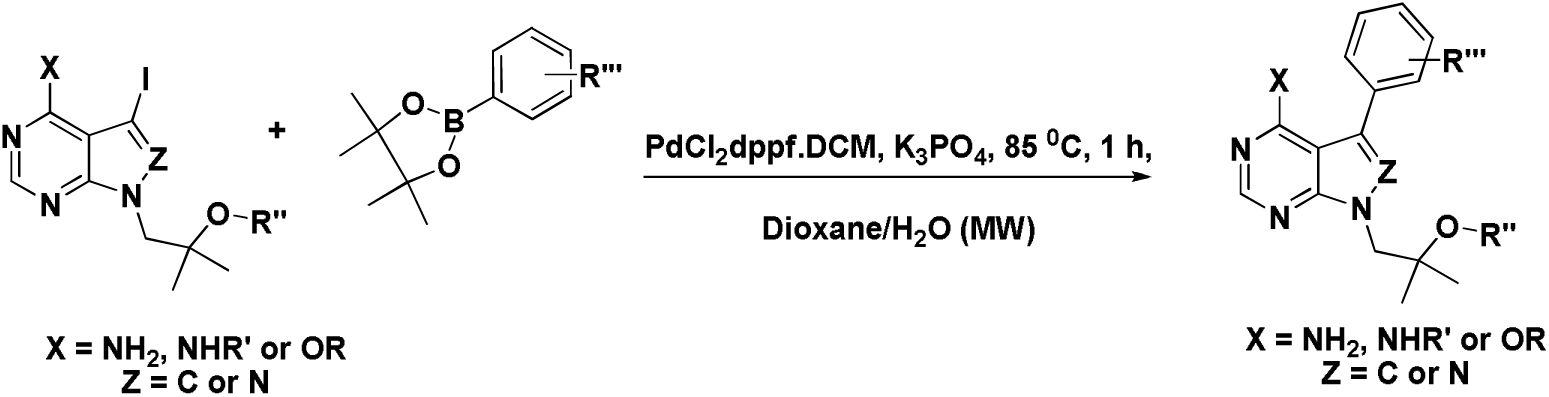

#### General Suzuki coupling condition to obtain the final compounds

The appropriate 3-iodo pyrazolopyrimidine or 5-iodo pyrrolopyrimidine scaffold (1.0 eq), the desired boronic acid or the boronate pinacole ester (1.5 eq), PdCl_2_.dppf.DCM (0.05 eq), K_3_PO_4_ (2.2 eq) were heated to 85 °C for 1 h in a microwave using a 3:1 mixture of dioxane/water as the solvent. The crude mixture was concentrated and purified by HPLC using ACN/water as the mobile phase to obtain the final compounds.

### 1553 (aka BKIDC-1553)

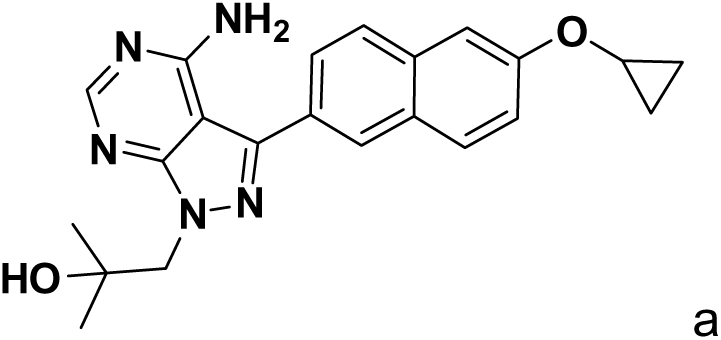

1-(4-amino-3-(6-cyclopropoxynaphthalen-2-yl)-1H-pyrazolo[3,4-d]pyrimidin-1-yl)-2-methylpropan-2-ol was synthesized and characterized as previously described (35) (identified as Compound **32**).

### 1553-N-Me

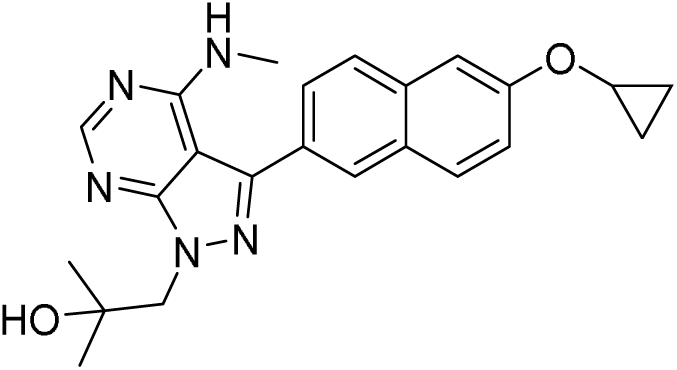

1-(3-(6-cyclopropoxynaphthalen-2-yl)-4-(methylamino)-1H-pyrazolo[3,4-d]pyrimidin-1-yl)-2-methylpropan-2-ol: ^1^H NMR (500 MHz, CDCl_3_) δ 8.47 (s, 1H), 8.05 (s, 1H), 7.94 (d, *J* = 8.4 Hz, 1H), 7.84 (d, *J* = 9.1 Hz, 1H), 7.73 (d, *J* = 8.4 Hz, 1H), 7.53 (s, 1H), 7.25 (m, 1H), 5.52 (q, *J* = 4.8 Hz, 1H), 4.48 (s, 2H), 3.91 (m, 1H), 3.09 (d, *J* = 4.8 Hz, 3H), 1.26 (s, 6H), 0.90-0.88 (m, 4H). Calcd for C_23_H_25_N_5_O_2_ 403.2, found (M+H)^+^ 404.4. HPLC purified product was determined to be ≥95 % pure under 220 nm and 254 nm detection in analytical HPLC under both solvent systems.

### 1817

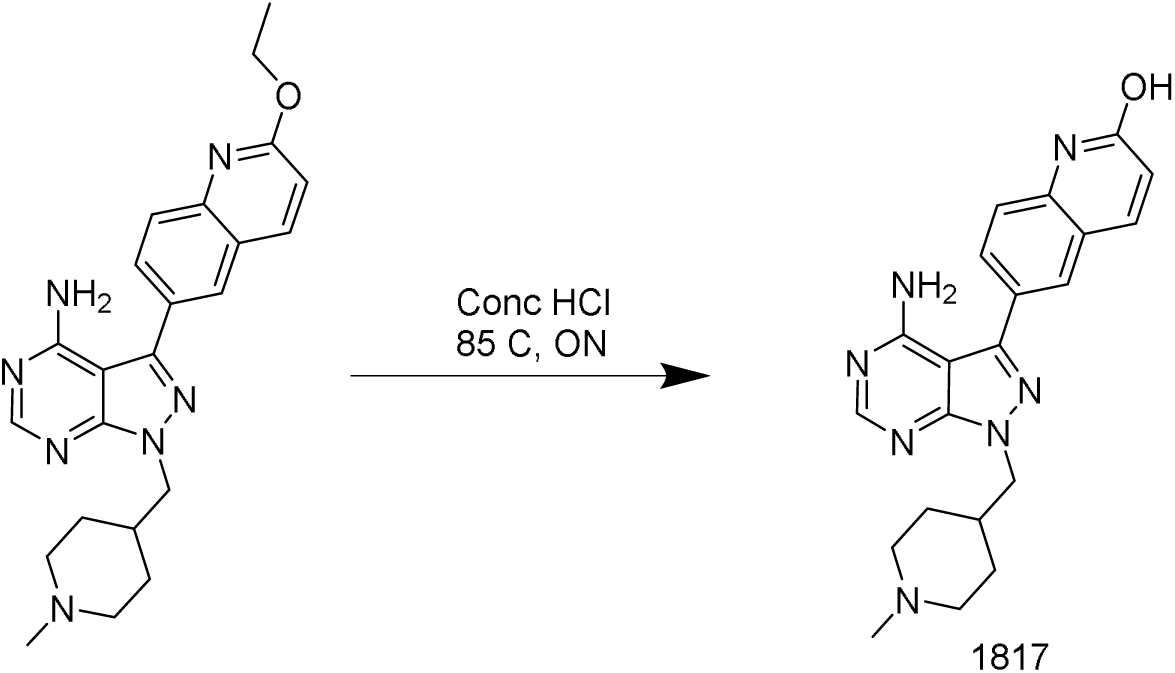

Conc HCl (3 mL) was added to 1-(6-ethoxynaphthalen-2-yl)-3-((1-methylpiperidin-4-yl)methyl)imidazo[1,5-a]pyrazin-8-amine(35)(50 mg) and heated overnight at 85 C. Crude mixture was diluted with aq. NaHCO_3_ and the product was extracted using ethyl acetate. The organic layer was dried over Na_2_SO_4_, concentrated, and purified by HPLC using ACN/water as the mobile phase to obtain 1817. ^1^HNMR(300 MHz, MeOD) δ 8.39 (s, 1H), 8.11 – 7.98 (m, 2H), 7.89 (dd, *J* = 8.5, 1.6 Hz, 1H), 7.55 (d, *J* = 8.5 Hz, 1H), 6.71 (d, *J* = 9.5 Hz, 1H), 4.46 (d, *J* = 6.7 Hz, 2H), 3.56-3.52 (m, 2H), 3.09 – 2.93 (m, 2H), 2.85 (s, 3H), 2.42-2.38 (s, 1H), 1.99-1.95 (m, 2H), 1.70-1.63 (m, 2H). Calcd for C_21_H_23_N_7_O 389.20, found (M+H)^+^ 309.1. HPLC purified product was determined to be ≥99 % pure under 220 nm and 254 nm detection in analytical HPLC under both solvent systems.

### 1826

Compound D (0.064 mmol, 1 eq) and sodium iodide (0.13 mmol, 2 eq) were dissolved in DMF (0.3 mL). The mixture was cooled to 0 °C and NaH (0.13 mmol, 2 eq) was added followed by compound C (0.096 mmol, 1.5 eq). The reaction mixture was allowed to come to 70 °C and allowed to stir over night. The mixture was diluted with water and the crude product was extracted into ethyl acetate. The organic layer was dried over sodium sulfate and concentrated to obtain compound E (Calcd for C_40_H_56_N_6_O_9_ 764.41, found (M+Na)^+^ 787.6). Crude E was then stirred with a 30% TFA/DCM mixture (1.6 mL) for 3 h at room temperature. The crude mixture was concentrated by rotary evaporation and purified by flash chromatography using a 5% methanol/DCM supplemented with 2% triethylamine as the mobile phase to obtain ∼90% pure 1826 with a 32% yield. ^1^H-NMR(300 MHz, MeOD) δ 8.36 (s, 1H), 8.16 (s, 1H), 8.01 (d, J = 8.6 Hz, 1H), 7.94 (d, J = 9.0 Hz, 1H), 7.81 (dd, J = 8.5, 1.6 Hz, 1H), 7.65 (d, J = 2.3 Hz, 1H), 7.27 (dd, J = 8.9,

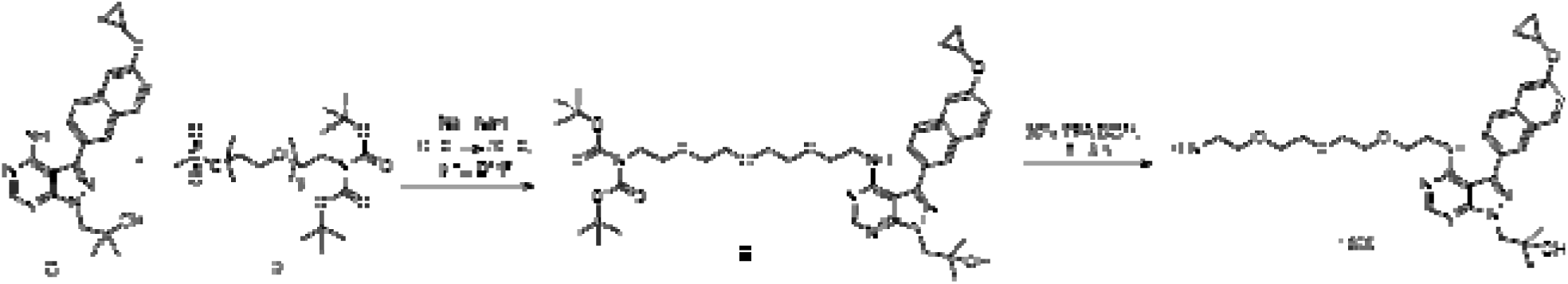

2.4 Hz, 1H), 4.44 (s, 2H), 4.03 – 3.97 (m, 1H), 3.78 – 3.67 (m, 9H), 3.60 – 3.57 (m, 2H), 3.51 – 3.48 (m, 3H), 3.43-3.39 (m, 2H), 3.27-3.24 (m, 1H), 3.15-3.10 (m, 1H), 3.05 – 2.99 (m, 2H), 1.30 (s, 6H), 0.94 - 0.90 (m, 2H), 0.86 – 0.74 (m, 2H). Calcd for C_30_H_40_N_6_O_5_ 564.3, found (M+H)^+^ 565.6.

### Immunohistochemistry (IHC)

Xenograft tumors were fixed with formalin and then paraffin embedded. The resulting FFPE (formalin-fixed, paraffin-embedded) samples were sectioned at 10 µm, mounted on slides, deparaffinized, rehydrated, heated up in Tris-based buffer (Vector Laboratories, Cat# H-3301-250) in the pressure cooker to accomplish antigen unmasking. Endogenous peroxidases were inactivated with 3% (v/v) H_2_O_2_ (Fisher Scientific, Cat# H325) in Tris-buffered saline. The sections were then blocked in 2% (v/v) normal goat serum (Invitrogen™, Cat# 10000C) at room temperature for 1 h before application of primary antibody incubation at 1:500 dilution (Phospho-Acetyl-CoA Carboxylase (Ser79) (D7D11) Rabbit mAb (Cell Signaling Technology, Cat# 11818, RRID:AB_2687505)) overnight at 4°C. The 3,3′-diaminobenzidine (DAB) based-chromogenic detection was performed using biotinylated secondary antibodies in conjunction with VECTASTAIN® ELITE® ABC HRP Detection Kit (Vector Laboratories, Cat# PK-6100) and ImmPACT® DAB Substrate Kit, Peroxidase (HRP) (Vector Laboratories, Cat# SK-4105). DAB-stained slides were counterstained with hematoxylin, dehydrated, and mounted with a coverslip. Appropriate negative controls were run in parallel. For DAB-stained slides, at least three digital images per section were obtained at 20× magnification with EVOS™ M7000 Imaging system (ThermoFisher Scientific) with the same magnification and exposure. The images were opened in ImageJ (RRID:SCR_003070, NIH image analysis freeware, http://imagej.nih.gov/ij/) and converted to RGB stack. The threshold of the green channel was optimized until the DAB-stained areas showed optimal contrast in the red channel. The same optimized threshold was used for all the images and the intensity of DAB staining was expressed as a percentage of stained area per a standard 20x field.

#### Kinobead profiling of 1553-N-Me using a 1:1 mixture of HCT116/HEK293T lysate Competitive kinobead profiling of 1553-N-Me

Kinobeads were synthesized by immobilizing five pan-kinase inhibitors on NHS activated sepharose (compounds 1 (33.3%), 2 (33.3%), 3 (8.3%), 4 (8.3%) and 7 916.7% from Golkowski et al (2017))(25), using a slightly modified protocol(25). The synthesized beads were stored as a 50% slurry in 20% ethanol until further use. A mixture (1:1) of HCT116 and HEK293T lysate was generated with a final concentration of 2 mg/mL in mod. RIPA buffer (pH 7.8, 50 mM Tris, 150 mM NaCl, 10 mM NaF, 1% NP40, 0.25% sodium deoxycholate, 5% glycerol, supplemented with Pierce Protease Inhibitor Tablets, 1 mM PMSF and phosphatase inhibitor cocktail 2 and 3 from Sigma Aldrich). 1553-N-Me (20 µM) or DMSO were added to the lysate mixture, followed by incubation for 30 min at 4 °C. 150 µL of DMSO- or 1553-N-Me-treated lysate was then added to 5 μL of pre-washed kinobeads. Samples were then rotated for 3 h at 4°C. Following kinobead enrichment, beads were washed with 500 μL of cold mod. RIPA lysis buffer (2x) followed by 500 μL of chilled TBS (50 mM Tris, 150 mM NaCl, pH = 7.8) (3x) in order to remove detergent. Each competitive profiling experiment was performed in quadruplicate. Following washing, kinobeads were resuspended in 50 μL of denaturing buffer (6M guanidinium chloride, 50 mM Tris containing 5 mM TCEP and 10 mM CAM, pH = 8.5) and heated to 95°C for 5 min. The bead slurry was the diluted with 50 µL of 100 mM TEAB (triethylammonium bicarbonate buffer, pH 8.5). 0.8 μg of LysC (Wako) was then added to the beads and the overall pH was adjusted to 8-9 with 1 N NaOH. A thermomixer (Eppendorf) was then used to agitate the mixture at 37°C for 2 h at 1400 rpm. Further dilution was performed by adding 100 µL of 100 mM TEAB, followed by the addition of 0.8 μg of sequencing grade trypsin (Pierce). Proteolytic digestion was performed for 14 h at 37°C at 800 rpm in the thermomixer. After overnight digestion, 200 µL of Buffer A (5% ACN, 0.1% TFA) containing 1% formic acid was used to digest the samples and the pH was adjusted to 2-3 with formic acid. Homemade stage-tips were prepared by running 50 μL of Buffer B (80% ACN, 0.1% TFA, H_2_O) through them followed by 50 μL of Buffer A (5% ACN, 0.1% TFA, H_2_O). The supernatants of spun down and kinobeads were added to the stage-tips. Stage-tip loading was performed by letting the proteolytic digest solutions pass through the stage-tips. Loaded stage-tips were then washed with 50 μL of Buffer A and eluted with 50 μL of Buffer B. Samples were dried with a speedvac and the resuspended in Buffer A. Resuspended samples were then injected onto an LC-MS. Samples were separated using a nanoAcquity UPLC instrument and a homemade 10 cm capillary column packed with 5 μm 120 Å reverse-phase C18 beads (ReproSil-Pur®, Maisch). Liquid chromatography was performed over 120 min using an initial 20 min isocratic trapping of 3% Buffer B and a flow rate of 700 nL/min, followed by a 100 min gradient of 35% Buffer B to an 80% Buffer B gradient at 350 nL/min. LC Buffer A solvent was 0.1% Acetic acid and LC Buffer B was 99.9% ACN, 0.1% acetic Acid. Data were collected using a Thermo Scientific Orbitrap Fusion Tribrid mass spectrometer. Raw files were analyzed using MaxQuant, followed by Perseus by filtering intensity values only identified by site, reverse, or potential contaminant. To determine kinases that were significantly competed by treatment with 20 μM of 1553-N-Me, a two-tailed two-sample t-test in Perseus with an FDR of 0.05 was applied. Kinases were considered as being non-competed by an inhibitor if it had a Log2 ratio (Log2 Difference) <1.

### Competitive profiling of 155-N-Me with immobilized 1826

1826 beads were synthesized using a previously published protocol with slight modifications(25). In brief, 26.7 µL of a 20 mM DMSO stock of 1826 and 6 μL of triethyl amine were added to 0.8 mL of 50% slurry of pre-washed NHS-sepharose beads. The mixture was rotated at room temperature for 20 h in the dark. At the end of this time, supernatant was removed, and the beads were washed thrice with 1:1 DMF/ethanol solution. Then the beads were incubated with a mixture of 1:1 DMF/ethanol (528 μL), 1M aminoethanol dissolved in 20 mM acetic acid in 1:1 DMF/ethanol (272 μL) and 1 M EDCI (120 μL) for 12 h in dark at room temperature for capping. The beads were then washed 3x times with 1:1 DMF/ethanol, 2x with 0.5 M NaCl and finally with a 20% ethanol solution. The beads were stored at 4°C until further use as a 50% slurry in 20% ethanol. 1553-N-Me was profiled using 5 µL of the 1826 beads described above and a 1:1 mixture of HCT116/HEK293T lysate using the same protocol as described for kinobead profiling of 1553-N-Me.

### Measurement of in vitro AMPK activity

Measurement with recombinant proteins was assessed using recombinant human AMPK (α1β1γ1 complex). Protein kinase activity of recombinant full-length AMPK (A1/B1/G1) enzyme (SignalChem, Richmond, BC Canada) was determined using a non-radioactive assay which measure changes in initial concentration of ATP via luminescence. Enzyme kinase activity was measure in the presence of 45 µM peptide substrate SAMStide (HMRSAMSGLHLVKRR) (SignalChem) in a buffered solution containing 5 mM MOP (pH 7.5), 5 mM MgCl_2_, 1 mM EGTA, 2.5 mM β-glycerol-phosphate, 0.5 mM EDTA and 0.05 mM DTT. A series of AMPK (A1/B1/G1) concentrations i.e., 40, 25, 15, and 10 ng were used to detect enzyme activity enhancement. The reaction was initiated with addition of 1 µM ATP. Incubation of enzyme reaction was at 37°C and 90 rpm agitation for 90 min. The assay was terminated with the addition of Kinase-Glo^®^ Luminescent Kinase Assays reagent (Promega, Cat# V6711) and read in an EnVision Multilabel Plate Reader (Perkin Elmer, USA). Compound A-769662 (Cayman Chemical, Cat# 11900) is a known positive control that stimulates AMPK activity, Compound C (Cayman Chemical, Cat# 11967), also known as dorsomorphin, is a known inhibitor of AMPK activity (both of these compounds were used at 2 µM final concentrations), and DMSO was used as a no compound control.

### Activity measurement of hexokinase

The full-length human HK2 open reading frame (ORF) flanked by the linkers was amplified by PCR using plasmid FLHKII-pGFPN3 (RRID:Addgene_21920) as a template with primers (*GGGTCCTGGTTCG*<u>ATG</u>ATTGCCTCGCATCTGCTTGCCTACTTC and *CTTGTTCGTGCTGTTTA*<u>CTA</u>TCGCTGTCCAGCCTCACGGATGCGGC: Italic, underlined, and double underlined text correspond to linker, initiation codon, and stop codon, respectively.) The resulting PCR fragment was cloned into bacterial expression plasmid AVA0421 by ligation-independent cloning(44, 45). The sequence integrity of the cloned fragment (GenBank accession No.: NM_000189.5) was confirmed by Sanger sequencing at GENEWIZ (South Plainfield, NJ). Human HK2 protein was expressed in *Escherichia coli* BL21(DE3) using Studier auto-induction method at 20°C and 180 rpm(78), followed by cell lysis and purification using immobilized metal-affinity chromatography (IMAC) in a Ni–NTA (Qiagen, Valencia, CA) column in binding buffer composed of 20 mM HEPES pH 7.25, 500 mM NaCl, 5% glycerol, 30 mM imidazole, 0.5% CHAPS and 1 mM tris(2-carboxyethyl)phosphine (TCEP) as previously described(79). This was followed by elution in the same buffer supplemented with 250 mM imidazole and size-exclusion chromatography in a 26/60 Superdex 75 size exclusion column. SDS-PAGE analysis revealed that the purified HK2 enzyme was near homogeneity.

For **Figure S13A** and **B**, hexokinase activity was measured using Hexokinase Colorimetric Assay Kit (Sigma-Aldrich, Cat# MAK091) on 96-well plates according to the manufacturer’s instructions. 50 μL of sample containing either LNCaP crude extract or recombinant HK2 protein (R&D Systems™, Cat# 8179HK020) was added to a well of 96-well plates. The reaction was initiated by addition of 50 μL reaction mixture containing HK Assay Buffer, HK Enzyme mix, HK Developer, HK Coenzyme, HK Substrate, and test compounds (BKIDC-1553-N-Me or 3-bromopyruvate (Sigma-Aldrich, Cat# 16490) using DMSO as a vehicle control). Hexokinase activity was determined by measuring a colorimetric product with absorbance at 450 nm proportional to the enzymatic activity present. For **Figure S13C**, in-house recombinant human HK2 protein was used to evaluate the effect of BKIDC on glucose phosphorylation activity. The non-radioactive Kinase-Glo^®^ Luminescent Kinase Assays (Promega, Cat # V6711) was used to measure ATP consumption in the presence of its natural substrate glucose. Kinase reactions were performed in 25 mM HEPES-KOH, pH 7.5 containing 150 mM NaCl, 10 mM MgCl_2_, and 10 mM CaCl_2_. The reaction mixture contained 1 mM of D-glucose substrate, 10 µM of BKIDC or 100 µM 3BP. The activity of HK2 was measured in a total reaction volume of 25 µL using a serial dilution of the enzyme in the range between 70 and 840 nM. All reactions were initiated by the addition of 10 µM ATP and terminated after 120 min incubation at 37°C and 90 rpm shaking with addition of Kinase-Glo® reagent. Change in initial ATP concentration was measured as bioluminescence readout in counts per second on the Envision plate reader (PerkinElmer).

### Statistical Analysis

Statistical analyses were conducted using GraphPad Prism (RRID:SCR_002798) v8 and are indicated within all figures. One-way ANOVA followed by Tukey’s or Dunnett’s test for multiple comparison or 2-tailed unpaired t test was performed to determine significance, which was defined as *p* < 0.05.

## RESULTS

### BKIDC effects in Prostate Cancer cell lines

Our lead compound, BKIDC-1553 has oral availability with a favorable safety and pharmacokinetics profile (compound **32** in (35)). BKIDC-1553 was tested in the MTS (3-(4,5-dimethylthiazol-2-yl)-2,5-diphenyltetrazolium bromide) assay for proliferation inhibition in a series of human PCa cell lines (LNCaP, LNCaP95, C4-2, C4-2B, LAPC4, VCaP, PC3, DU145, LNCaP^APIPC^(28), MSKCC EF1 and NCI-H660) and non-prostate control lines (HepG2, U2OS, WIL2-NS and HFF) (**Figure 1A**). From the data we see that androgen receptor (AR) positive lines LNCaP, LNCaP95, C4-2, C4-2B, LAPC4 and VCaP as well as AR negative cell lines LNCaP^APIPC^ (LNCaP cell engineered to grow without AR(28)) and the neuroendocrine (NE) lines MSKCC EF1 and NCI-H660 respond to BKIDC-1553. Thus, the responsiveness to BKIDC-1553 was found in both AR positive and AR negative PCa cell lines. The concentration to inhibit cell proliferation by 50% ranged from 2.5-20 µM in these cell lines; concentrations achievable in experimental animals without toxicity. No response to BKIDC-1553 is seen in the AR-negative and non-NE PCa lines (PC3 and DU145). We also saw no response to BKIDC-1553 from non-PCa cell lines including a human bone cancer line (U2OS), human foreskin fibroblasts (HFF), a lymphoblastoid (WIL2-NS) or hepatoma (HepG2) cell lines (**Figure 1A**). We performed crystal violet growth assays three times with LNCaP95 and PC3 cells in dose response to BKIDC-1553 (**Supplementary Figure S3**). They clearly demonstrate inhibition of cell growth in the BKIDC sensitive cell line, LNCaP95 cells, but not the BKIDC resistant line, PC3. These experiments show that the effects seen with the MTS assay are due to cell proliferation and not due to metabolic responses. In summary, BKIDC-1553 was active in suppressing the growth of most AR positive and AR negative metastatic PCa cell lines but was not effective in suppressing the growth of normal fibroblast cells or bone, lymphocyte, or hepatocyte derived cancer cell lines that are tested in this paper.

**Figure 1.**
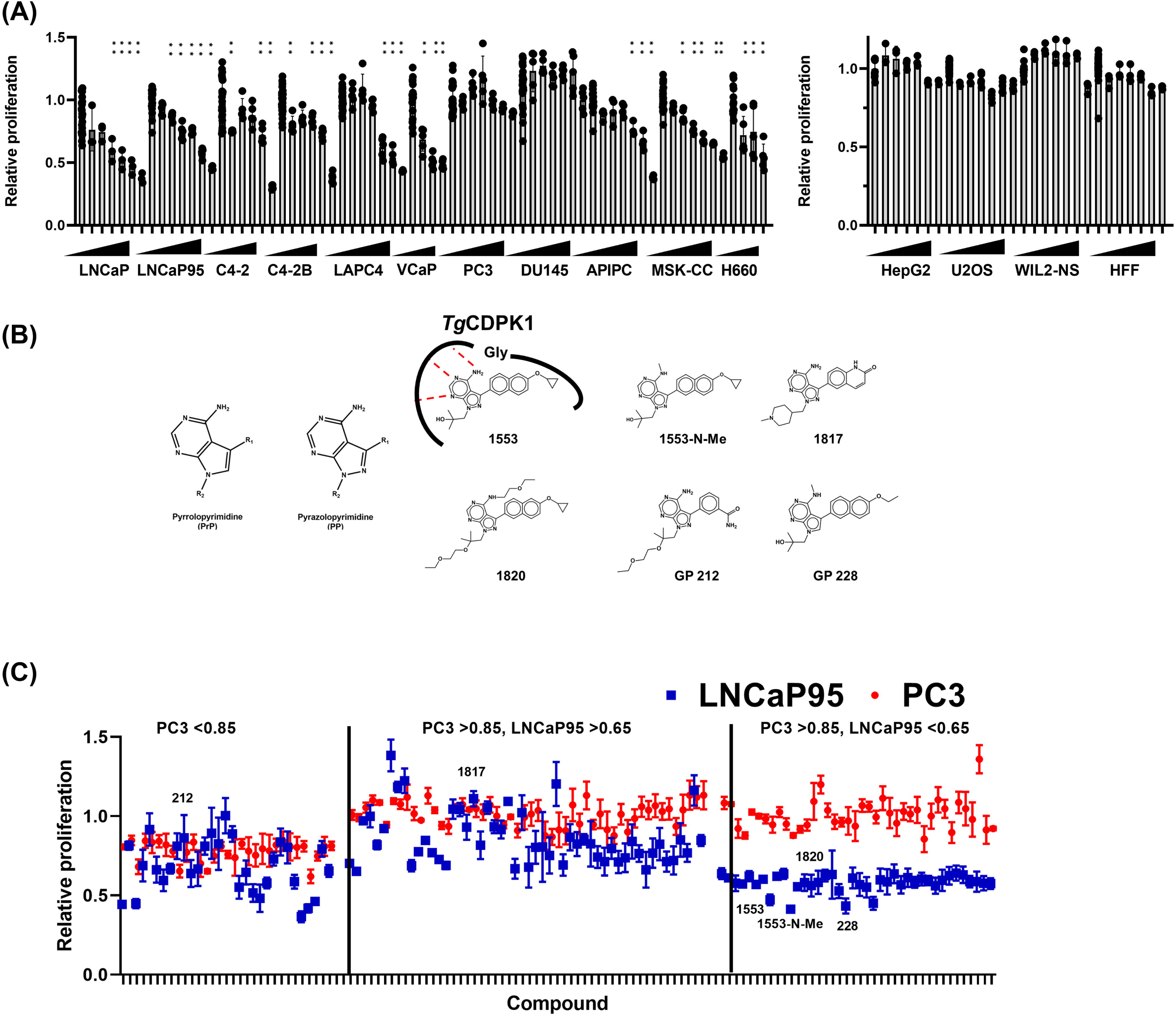
BKIDC effects on Prostate Cancer Cell lines and other Cell Lines. **(A)** Cell proliferation was evaluated by MTS assay at 72 h after BKIDC-1553 was added at the range between 0.63-20 µM, except for VCaP at 120 h (DMSO as a vehicle control). The data shown are representative of at least two independent experiments, normalized to the values obtained from cells grown in medium containing 0.1% DMSO vehicle and presented as mean ± standard deviation (SD) (n=3-5). One-way ANOVA followed by Dunnett’s test for multiple comparison: \*\**p* <0.01. **(B)** Structures of representative BKIDCs. Lead BKIDCs have two main scaffolds: Pyrrolopyrimidines (PrP) and pyrazolopyrimidines (PP) are two main scaffolds of BKIDCs. *Toxoplasma gondii* calcium dependent protein kinase 1 (*Tg*CDPK1) has the small gatekeeper residue glycine and can hold BKIDC-1553 in the ATP binding pocket. BKIDC-1553-N-Me, 1820, and GP228 have distinct R2 groups while commonly lacking the protein kinase binding and thus inhibition activity. **(C)** Graphical presentation of screening results of over 123 BKIDC compounds (**Table S1**) tested at 10 μM in LNCaP95 and PC3 (**Table S2**). Each dot represents one compound and the data are shown as mean ± SD. Compounds were categorized into three groups depending on the degree of inhibition of proliferation in LNCaP95 and PC3.

We next evaluated if PKD was the target of BKIDCs. We first noted that unlike previously reported PKD inhibitors(26, 27), BKIDC-1553 lacked activity against PC3, an AR negative and NE negative cell line. To gain insight into the targets of BKIDC-1553, we performed structure–activity relationship studies with small molecules based on pyrrolopyrimidine (PrP) or pyrazolopyrimidine (PP) scaffolds (**Figure 1B**) with LNCaP95 and PC3 as a representative of responders and non-responders, respectively (**Table S1 and S2**). A series of compounds with variable R1 and R2 substituents still displayed antiproliferative activity in LNCaP95 but not PC3 (**Figure 1C and Table S1 and S2**). We created BKIDCs, represented by BKIDC-1553-N-Me that contain substituents (alkyl groups on the exocyclic nitrogen of pyrimidine) that disrupt canonical hinge region hydrogen bonding interactions in the ATP-binding site of protein kinases (**Figure 1B**). The protein targets of BKIDC-1553-N-Me were determined with competitive proteomic profiling experiments and no protein kinases, including PKD, MEK2, and RIPK2, were identified as hits (**Figure S1**). Yet, BKIDC-1553-N-Me maintained its antiproliferative activity against LNCaP95 (**Figure S2**). This demonstrated PKD. MEK2, and RIPK2 were not BKIDC targets and suggested the antiproliferative actions of BKIDCs on PCa cells acts via a mechanism not requiring protein kinase inhibition. Indeed, we confirmed that BKIDC-1553-N-Me showed a similar antiproliferative spectrum across the cell lines tested with BKIDC-1553 (**Figure S2**). In this screen, we also found a closely-related molecule, BKIDC-1817 (**Figure 1B**: a 1H-pyrazolo[3,4-d]pyrimidin-4-amine compound)(46), had no inhibitory activity in LNCaP95 or PC3 cells (**Figure S2**). BKIDC-1817 served as a negative control in subsequent studies.

### BKIDC effects on cell cycle and cell-signaling pathways

We carried out a series of experiments to characterize cell cycle effects, cell-signaling pathways, and energy sources to investigate the cellular processes inhibited by BKIDC-1553 and -1553-N-Me. Thirty-hour exposure to BKIDC-1553 and 1553-N-Me commonly resulted in G1 cell cycle arrest of responders (LNCaP and LNCaP95), but not a non-responder line (PC3) which had no enrichment of the G1 population (**Figure 2A, Figure S4A and S4C**). In addition, our negative control BKIDC-1817 did not affect the cell cycle of any cell line. Similar patterns of cell cycle profiles were observed at 48- and 72-h time points with no obvious appearances of the sub-G1 population in LNCaP95 (47)(**Figure S4B**), suggesting that cytostatic rather than cytopathic actions of BKIDCs are responsible for their antiproliferative activities.

**Figure 2.**
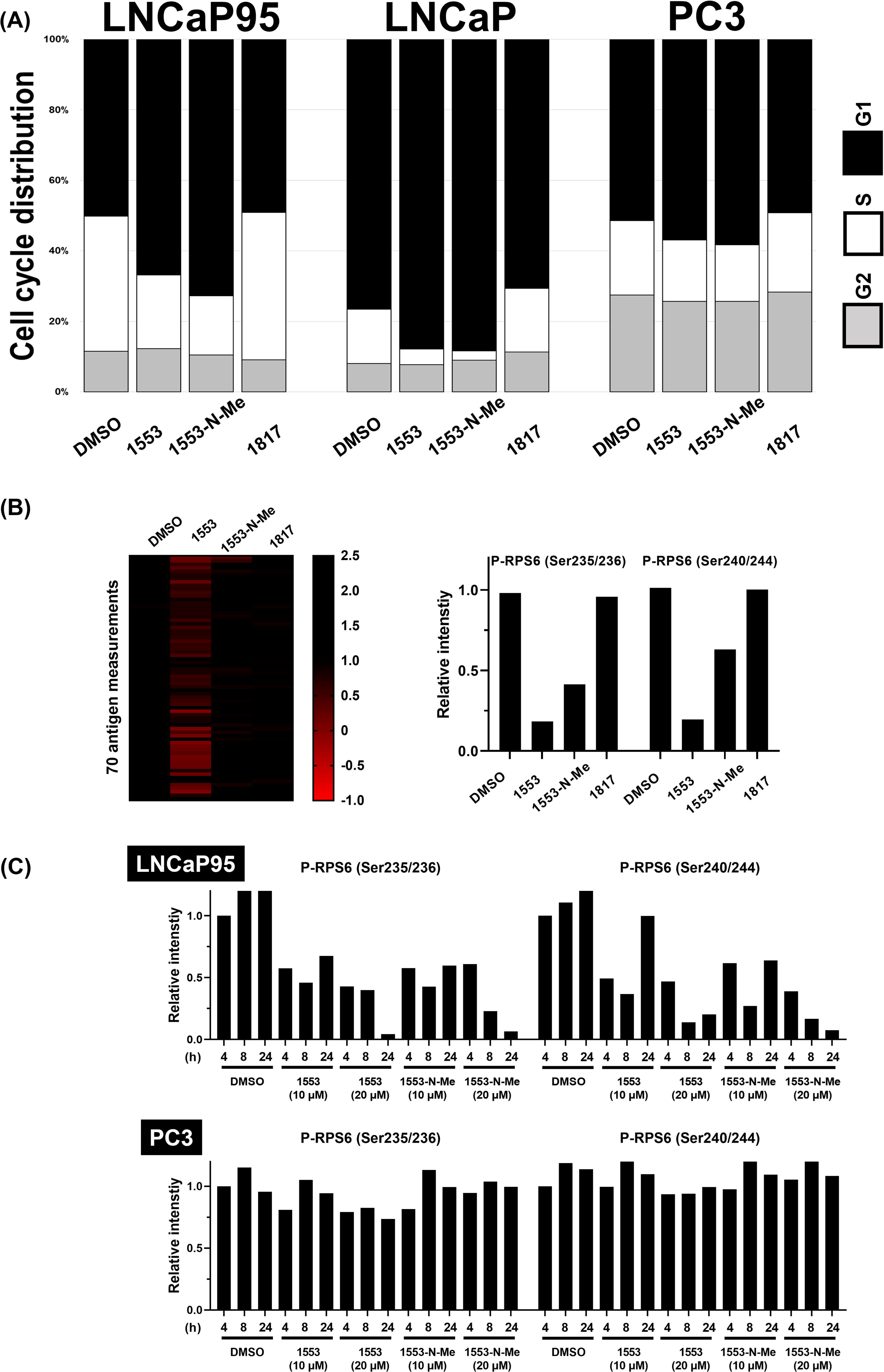
Cytostatic effects of anti-proliferative BKIDCs. **(A)** The percentage of cells at G1, S, and G2 phase of cell cycle at 30 h after treatment with the indicated BKIDCs. Cell cycle analysis showing G1 cell cycle arrest in LNCaP and LNCaP95 cells compared to PC3 when treated with BKIDC-1553 and 1553-N-Me. These results were confirmed in 2 subsequent independent experiments (**Figure S4**) **(B)** Heat map of antigen levels detected by RPPA. Cellular lysates of LNCaP95 treated with the indicated BKIDC at 20 µM for 24 h were subjected to RPPA with 70 distinct antibodies (Antibodies and the corresponding RPPA signal intensities relative to β-actin signals are listed in **Table S3**). Note BKIDC-1553 and 1553-N-Me commonly affected only abundance of ribosomal S6 protein (RPS6) phosphorylated at S235/S236 or S240/S244, the first two bands seen with BKIDC-1553 and the only bands seen with BKIDC-1553-N-Me. The response to BKIDC-1817 was comparable to the one to DMSO. **(C)** Time course and dose dependent studies of BKIDCs in LNCaP95 and PC3. Time- and dose-dependent decrease in phosphorylated RPS6 signals in RPPA in LNCaP95 but not PC3 as early as at 4 h after treatment with BKIDC-1553 and 1553-N-Me.

Since G1 but not G2/M arrest was dominantly associated with BKIDC-induced changes in cell cycle profiles, we took advantage of reverse-phase protein arrays (RPPA) to characterize robust biochemical alterations leading to G1 cell cycle arrest(48) (**Figure 2B and 2C, Figure S5, Table S3**). This high-throughput antibody-based targeted proteomics platform allowed us to observe a time- and dose-dependent decrease in phosphorylated ribosomal protein S6 at serine 235/S236 (P-RPS6^S235/S236^) and serine 240/S244 (P-RPS6^S240/S244^) in LNCaP95 but not PC3 cells as early as 4 h after treatment with BKIDC-1553 and 1553-N-Me (**Figure 2C, Figure S6**). RPS6 is a canonical downstream target of mTORC1 kinase which is negatively regulated by the metabolic sensor AMP-activated protein kinase (AMPK)(49, 50). Indeed, antiproliferative BKIDC-1553-N-Me, but not our negative control BKIDC-1817, consistently activated AMPK in LNCaP95 cells as reflected by increased enzymatic activity and increased phosphorylation of acetyl-CoA carboxylase (ACC) at Ser79 (the major substrate for AMPK)(49) (**Figure 3A and 3B**).

**Figure 3.**
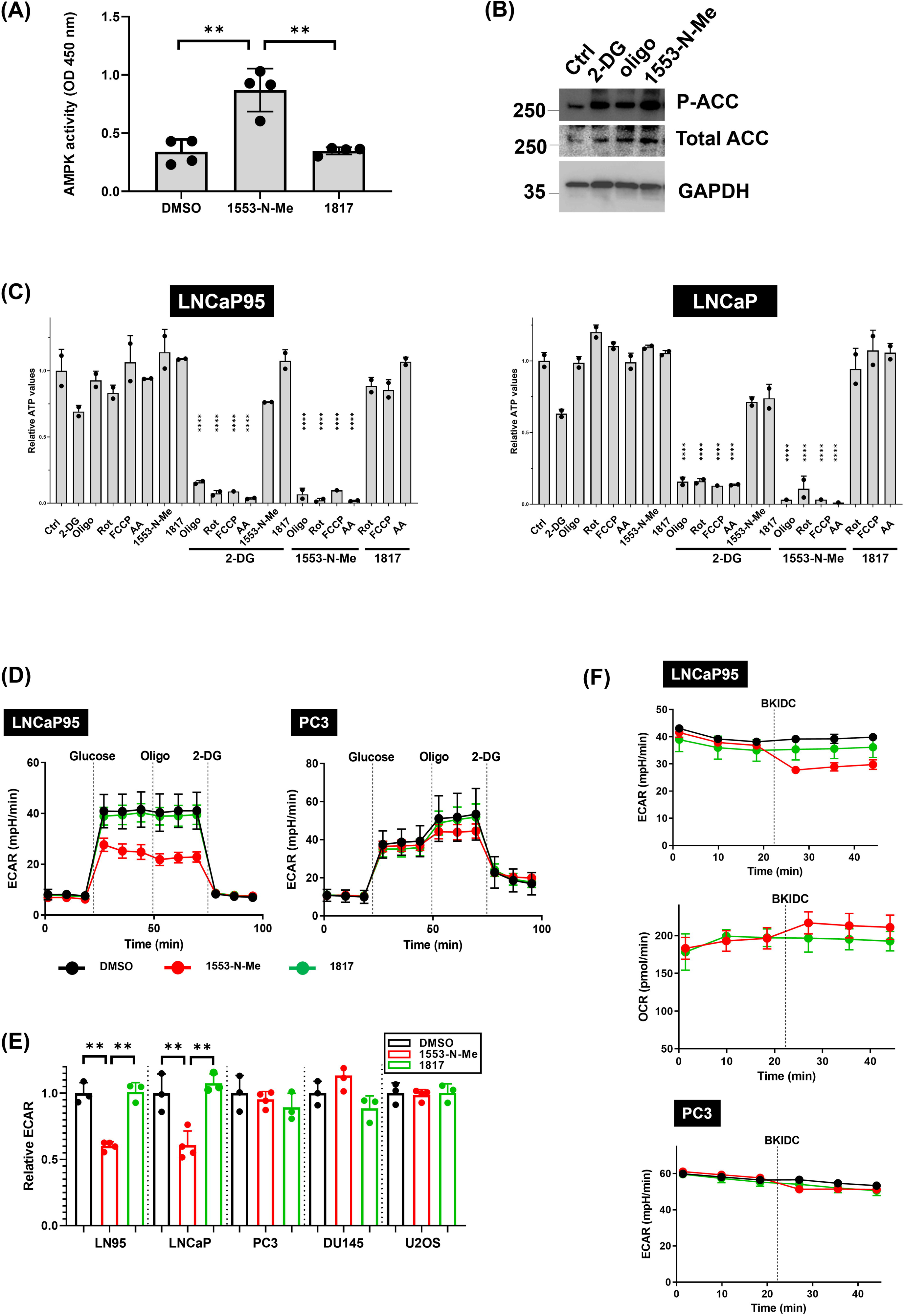
BKIDCs are antiglycolytic agents. **(A)** AMPK activation in LNCaP95 treated with BKIDC-1553-N-Me. AMPK activity in total LNCaP95 lysates was measured after 1 h exposure to BKIDC-1553-N-Me or 1817. One-way ANOVA followed by Tukey’s test for multiple comparison: \*\**p* <0.01. **(B)** Phosphorylated acetyl-CoA carboxylase (P-ACC) at Ser79 was increased after treatment with BKIDC-1553 but not 1817. Metabolic inhibitors 2-deoxyglucose (2-DG) and oligomycin (oligo) served as positive controls for AMPK activation to increase P-ACC. **(C)** Effects of BKIDCs on intracellular ATP levels in LNCaP and LNCaP95 cells. Cells were treated with BKIDC-1553-N-Me, glycolysis inhibitor 2-DG, and OXPHOS inhibitors (OXPHOSi) singly or in combination for 120 min. The data shown are representative of at least two independent experiments, normalized to the values obtained from cells grown in medium containing 0.1% DMSO vehicle and presented as mean ± SD (n=2). Rot: rotenone, FCCP: carbonyl cyanite-4 (trifluoromethoxy) phenylhydrazone, AA: Antimycin A. One-way ANOVA followed by Dunnett’s test for multiple comparison (vs. Oligomycin alone). **(D)** Extracellular acidification rates (ECAR) were determined in cultured cells by the Seahorse XFe24 analyzer. Cells were pre-treated with 20 µM BKIDC for 1 h prior to application to Seahorse analyzer. Increase in ECAR by glucose injection (glycolytic rates) was inhibited by BKIDC-1553-N-Me in LNCaP95 but not PC3. Maximum glycolytic capacity was evaluated by oligomycin injection. 2-DG injection shuts down glycolysis as reflected by return of ECAR to the basal levels. **(E)** The effect of BKIDC-1553-N-Me on glycolytic rates. The data shown are normalized to the values obtained from cells pre-treated with DMSO vehicle and presented as mean ± SD (n=3). One-way ANOVA followed by Tukey’s test for multiple comparison: \*\**p* <0.01. **(F)** Cells were exposed to BKIDC at the indicated time by injection at the final concentration of 10 µM. Decrease in ECAR was observed in LNCaP95 but not PC3 immediately after injection of BKIDC-1553-N-Me. There was a compensatory increase in oxygen consumption rates (OCR) which reflects mitochondrial oxidative respiration.

We next performed an in vitro enzyme assay using purified human AMPKα1β1γ1 complex to examine whether BKIDC-1553 and 1553-N-Me could be direct activators of AMPK. A well-established allosteric activator A-769662 caused 2-fold increase in AMPK activity (51) while BKIDCs failed to do so, suggesting BKIDCs do not stimulate AMPK activity directly (**Table S4**).

Since AMPK serves as a cellular sensor of ATP-ADP-AMP balance in cells, the alteration in ATP metabolism could be upstream of pathways that affect ATP levels and produce AMPK activation(49). We examined BKIDC’s effects on ATP supply from two major pathways, glycolysis in the cytosol and oxidative phosphorylation (OXPHOS) in mitochondria. OXPHOS inhibition, which was achieved by oligomycin (inhibitor for F0 component of H^+^-ATP-synthase), antimycin A (AA: inhibitor for complex III), rotenone (rot: complex I inhibitor), or carbonyl cyanite-4 (trifluoromethoxy) phenylhydrazone (FCCP: uncoupler)(52), cooperated with BKIDCs to deplete intracellular ATP (**Figure 3C, Figure S7**). This suggested that BKIDCs inhibit glycolysis-derived ATP production, and indeed a similar effect was seen with the glycolysis inhibitor 2-deoxyglucose (2-DG). Also, similar to 2-DG, BKIDC-1553 and -1553-N-Me cooperated with oligomycin to exhibit cytopathic, apoptotic actions in LNCaP as judged by increased PARP cleavage and cell detachment (47) (**Figure S8**).

### BKIDC effects on glycolysis

To examine the effects of BKIDCs on glycolysis directly, we employed the Seahorse assay to measure the extracellular acidification rate (ECAR), a proxy for glycolysis rate in cells(53). ECAR was suppressed by BKIDCs in LNCaP and LNCaP95 cells but not in BKIDC non-responder cell lines, including DU145, PC3, and U2OS (**Figure 3D and 3E, Figure S9**). Mitochondrial respiration as judged by oxygen consumption rate (OCR) (54) was not affected by BKIDC-1553-N-Me in both responder (LNCaP, LNCaP95) and non-responder cell lines (PC3, DU145)(**Figure S9**). The rapid decrease in ECAR with BKIDCs suggests that BKIDCs have a direct antiglycolytic effect (**Figure 3F**). Under the same conditions, BKIDCs induced a moderate compensatory increase in the oxygen consumption rate (OCR) in LNCaP95(**Figure 3F**), suggesting aerobic, mitochondrial oxygen consumption partially compensates for inhibition of glycolysis.

In order to determine which glycolytic step is interrupted by BKIDCs, we performed a metabolic flux assay with D-[U-^13^C] glucose(55) (**Figure 4A and 4B**). We exposed cells to labelled glucose with different concentrations of BKIDC-1553-N-Me and DMSO as a solvent control for 30 min. We did not see significant changes in glucose levels among cells lines and treatment. In LNCaP and LNCaP95 cells, we clearly saw a decrease in the levels of 6 carbon sugar phosphate which includes glucose 6-phosphate and fructose 6-phosphate. Consistently we observed marked reduction in levels of downstream metabolites, including dihydroxyacetone phosphate (DHAP), phosphoglycerate (PG), phosphoenolpyruvate (PEP), and pyruvate (**Figure 4B**). Importantly, the non-responder cell line DU145 did not display such metabolic alterations. These experiments suggested that either glucose uptake or glucose phosphorylation were affected by BKIDC exposure in susceptible PCa cells(52). Supporting this, results from the Glucose Uptake-Glo Assay also were consistent with BKIDC-1553-N-Me blocking glucose uptake and/or phosphorylation (**Figure S10A**).

**Figure 4.**
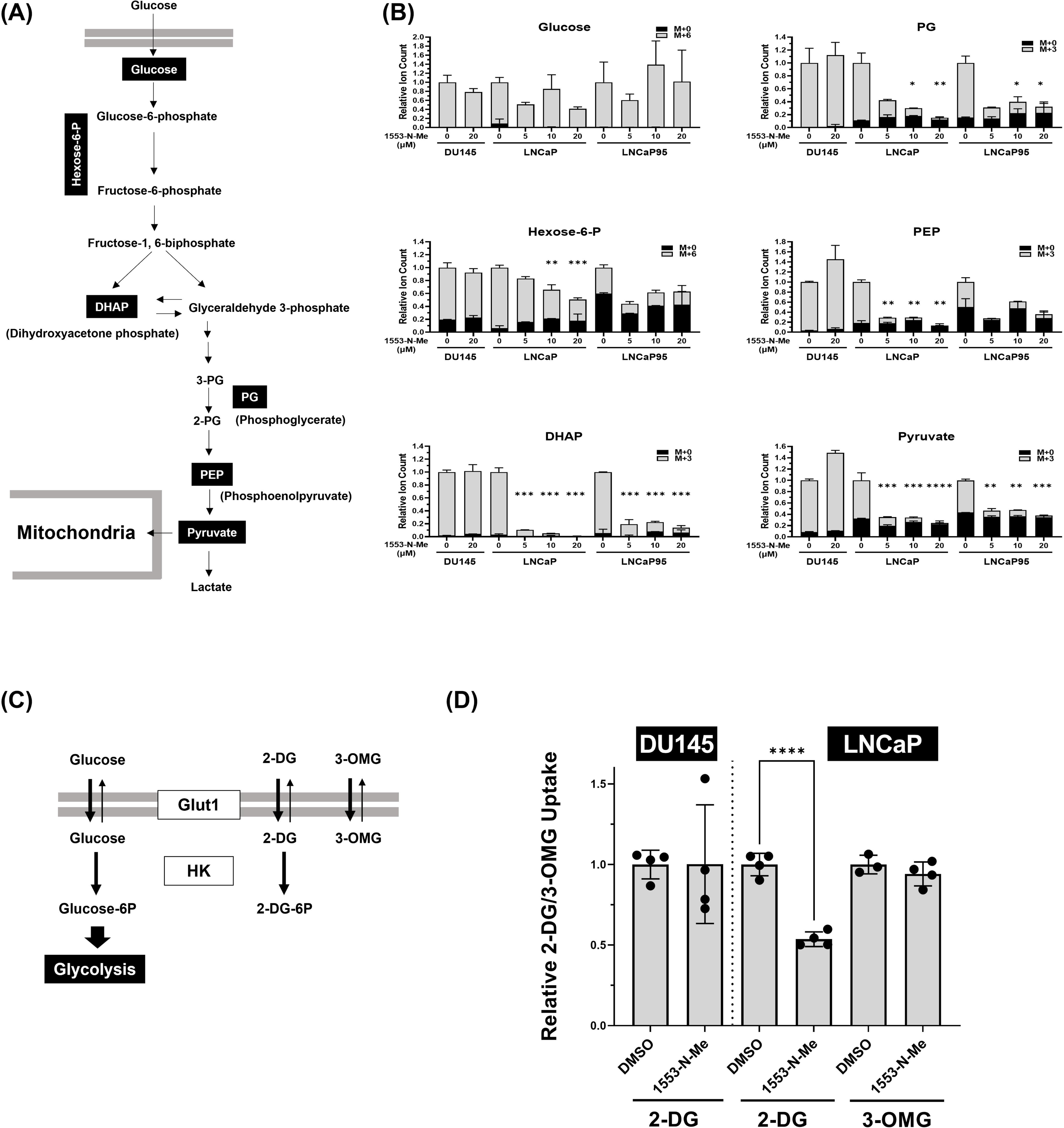
Glucose phosphorylation is the primary target for BKIDC as an antiglycolytic agent. **(A)** Schematic flowchart of glycolysis and the metabolites measured in this metabolic flux assay with ^13^C-labeled glucose with BKIDC-1553-N-Me this experiment (dark boxes). **(B)** Metabolic flux assay with ^13^C-labeled glucose. ^13^C enrichment of glycolytic intermediates was determined by liquid chromatography coupled with mass spectrometry (LC-MS) in DU145, LNCaP, and LNCaP95 cells at 30 min after exposure to [U-^13^C]-glucose in the presence and absence of BKIDC-1553-N-Me. The X-axis is the concentration of BKIDC-1553-N-Me, with 0-20 μM added to the indicated cell lines. Black columns indicate where unlabeled carbon (^12^C, thus M+0 [M=mass expected for ^12^C]) is found and grey columns originate from the ^13^C-labeled glucose (^13^C, thus M+6 or M+3). In LNCaP and LNCaP95 cells, addition of BKIDC-1553-N-Me inhibited glucose being metabolized to 3-carbon glycolytic intermediates, DHAP, PG, PEP, or pyruvate, indicating a block in glycolysis, at or prior to the six-carbon to 3-carbon fructose-bisphosphate aldolase step. Note that in BKIDC non-responsive DU145 cells there is no BKIDC-1553-N-Me effect on hexose-6-phosphate and 3-carbon glycolytic intermediates (DHAP, PEP, PG, or pyruvate). The results trended to indicate that BKIDC blocked effects at hexose-6-phosphate levels. One-way ANOVA followed by Dunnett’s test for multiple comparison for M +3 and M + 6 isotopomers: \**p* <0.05, \*\**p* <0.01, \*\*\**p* <0.001, \*\*\*\**p* <0.0001. **(C)** To distinguish between glucose uptake and phosphorylation steps, two different radiolabeled glucose analogs were used for radiometric assays in LNCaP and DU145 cells. Both 2-DG and 3-O-methylglucose (3-OMG) are taken up into cells in the same manner as glucose. Unlike 2-DG which is trapped by hexokinase (HK)-mediated phosphorylation and accumulated in the cells as a form of 2-DG-6P, 3-OMG is not phosphorylated, and its intracellular levels reflect the cells’ capability of Glut1-mediated glucose uptake itself. **(D)** 2-DG 2-[1,2-^3^H(N)] accumulation was inhibited by the presence of BKIDC-1553-N-Me at 20 μM in LNCaP but not DU145 cell lines. 3-O-[methyl-^3^H] D-glucose uptake was not affected by BKIDC-1553-N-Me in LNCaP. These data support the conclusion that BKIDC block phosphorylation but not uptake of glucose in BKIDC-susceptible cell line LNCaP cell line. Two-tailed t tests: \*\*\*\**p* <0.0001

To distinguish between glucose uptake and glucose phosphorylation steps, we used two different radiolabeled glucose analogs, [^3^H] 2-DG and [^3^H] 3-O-methyl D-glucose (3-OMG), which are taken up into cells in the same manner as glucose (**Figure 4C and 4D**). 2-DG undergoes phosphorylation to become trapped and accumulates in the cells as a form of 2DG-6P. On the other hand, 3-OMG cannot be phosphorylated and its intracellular levels reflect the cell’s capability of glucose uptake alone(30). We confirmed time-dependent accumulation of 2-DG signals in both LNCaP and DU145 (**Figure S10B**). This accumulation was inhibited by the presence of BKIDC-1553-N-Me in LNCaP but not DU145 cells. BKIDC-1553-N-Me did not inhibit 3-OMG uptake in LNCaP cells, suggesting that BKIDC inhibits the glucose phosphorylation step, not glucose uptake, in BKIDC susceptible cells (**Figure 4D**).

Hexokinases carry out the phosphorylation of glucose and we wanted to learn if BKIDCs were dependent on a specific isoform of hexokinase (HK). Among five isoforms of hexokinases expressed in humans (HK1, HK2, HK3, GCK1/HK4, and HKDC1)(56), HK1 and HK2 are reportedly predominant in LNCaP cells(57). We used a CRISPR-Cas9 system to generate HK1 and HK2 knock-out (KO) LNCaP clones. We successfully established several independent HK1 and HK2 KO clones, demonstrating that HK1 and HK2 are dispensable, and deletion of each gene did not affect cell proliferation and survival (**Figure 5 and Figure S11**). We then characterized their sensitivity to BKIDCs. We evaluated intracellular ATP levels, in the presence of oligomycin (OXPHOSi), to determine BKIDC’s antiglycolytic activity in the HK1 and HK2 KO lines. Like the parental LNCaP line, individual HK1 KO clones were sensitive to both BKIDC-1553 and 1553-N-Me, demonstrating lower ATP levels in the presence of oligomycin. In sharp contrast, HK2 KO clones displayed remarkable resistance to antiglycolytic action of BKIDCs, in that ATP levels were not suppressed in the presence of oligomycin. Pooled KO lines with different sgRNAs also displayed resistance to BKIDC only with HK2 KO pools, but not with HK1 KO (**Figure S12**). This suggested that BKIDCs exert their antiglycolytic activity through HK2, but not HK1, in susceptible PCa cell lines. Collectively, these findings implicate HK2-mediated glucose phosphorylation as the primary target of BKIDC. An unexpected finding was that both HK1 and HK2 KO clones retained antiproliferative sensitivity to BKIDCs (**Figure S11**). This suggests that BKIDCs may exert antiproliferative effects beyond effects on glycolysis. But from these experiments, it is clear that BKIDCs require HK2, but not HK1, for their effects on ATP generation from glycolysis. We explored whether HK2 could be directly targeted by BKIDCs. We were unable to demonstrate that BKIDCs inhibit human recombinant HK2 (hrHK2) enzymatic activity, in phosphorylation of glucose to glucose-6-phosphate (**Figure S13**). Similarly, HK2 expression does not determine whether a cell line is sensitive. BKIDC-resistant cell lines PC3 and DU145 express HK2 and its expression levels are comparable irrespective of susceptibility to BKIDC (**Figure S14**). This suggests that BKIDCs do not act directly on HK2, but likely act on regulatory molecules (pathway’s) that are known to modulate HK2 function(58).

**Figure 5.**
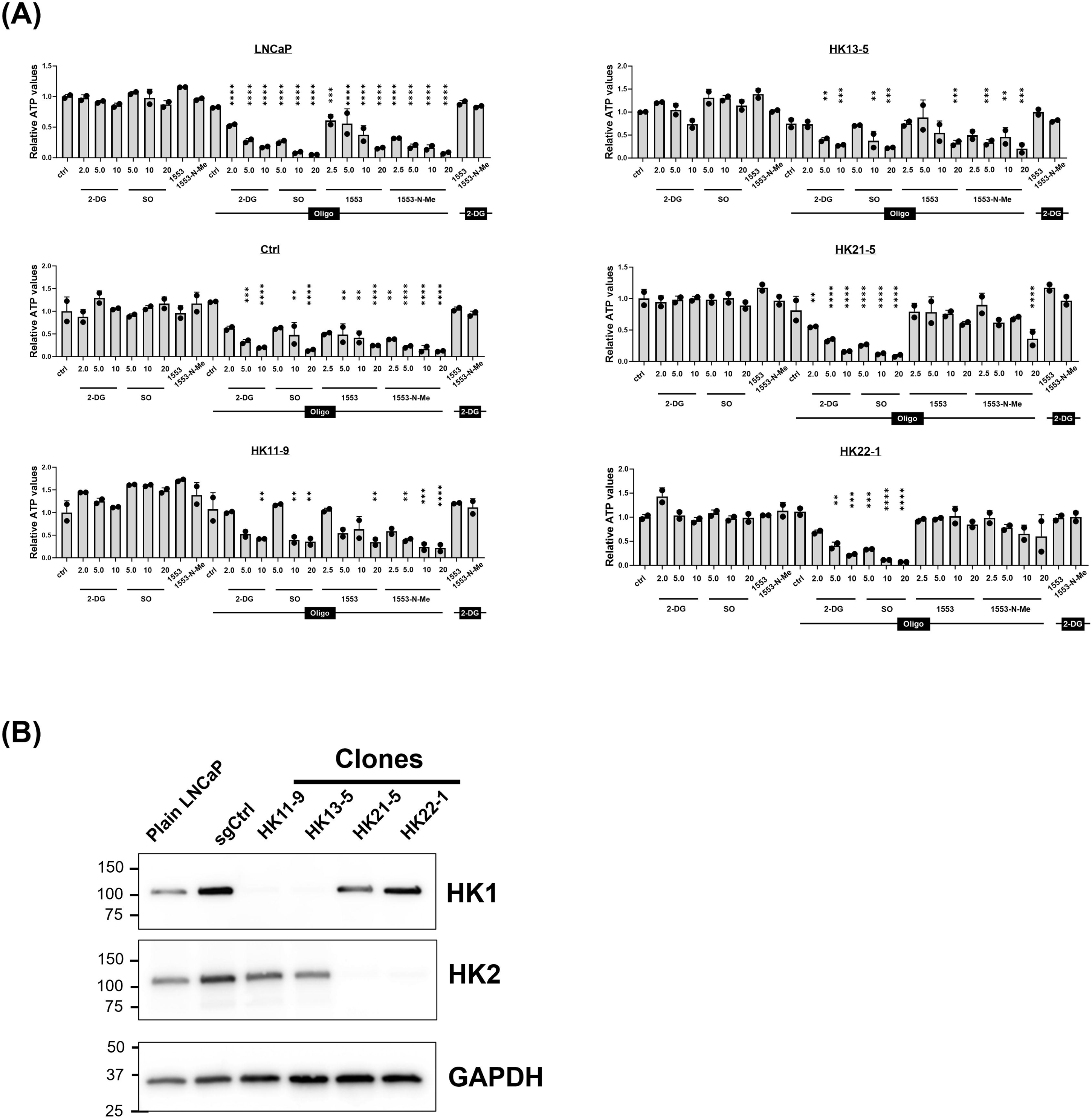
The effect of HK deficiency on antiglycolytic actions of BKIDC-1553-N-Me. **(A)** CRISPR-Cas9 system was used to generate HK1 and HK2 knock-out (KO) LNCaP clones and two independently derived KO clones for HK1 and HK2 were tested. Intracellular ATP levels, in the presence of a mitochondrial respiration blocker oligomycin (Oligo), were used as a proxy to monitor the activity of glycolysis to generate ATP. 2-Deoxyglucose (2-DG) and sodium oxamate (SO) were employed to block glycolysis at different steps. SO is an inhibitor of lactate dehydrogenase thus blocking the final step of glycolysis conversion of pyruvate into lactate. Note HK2 KO clones displayed remarkable resistance to decreasing ATP when co-treated with Oligo and BKIDC-1553-N-Me while they were sensitive to 2-DG and SO. One-way ANOVA followed by Dunnett’s test for multiple comparison (vs. Oligomycin alone). **(B)** Western blot validation of HK1 and HK2 deficiency in LNCaP KO clones. GAPDH served as a loading control.

### BKIDC effects in preclinical models of prostate cancer

We next tested the therapeutic potential of BKIDC-1553 in in vivo preclinical PCa models. The slow clearance of BKIDC-1553 from the systemic circulation of mice led to exposures of over 2 µM for an extended period of up to 72 h post-dose, which allowed a Monday, Wednesday, Friday dosing regimen (**Supplementary Figure S15**). BKIDC-1553, administered 20 mg/kg orally, 3 times/week, decreased tumor growth after 5 weeks in a human PCa patient-derived xenograft (PDX), LuCaP 35 (59) while PC3 tumors derived from adeno-PCa cell-line derived xenografts were resistant to BKIDC-1553 (**Figure 6A**). Intratumoral concentrations of BKIDC-1553 48 h after the end of dosing were similar in PC3 and LuCaP 35 tumors (about 8 µM, **Table S5**), ruling out that observed drug resistance of PC3 is due to low exposure to BKIDC-1553. Importantly, antitumor activities of BKIDC-1553 were comparable to those of standard ARSi enzalutamide treatment in AR+ LuCaP 35 model. As demonstrated in the responder line LNCaP95 cells in vitro (**Figure 3A**), we observed increased P-ACC immunoreactivities in 1553-treated LuCaP 35 tumors but not enzalutamide-treated tumors (**Figure 6B**). This implies 1553 also elicited metabolic rewiring in association with AMPK activation in vivo. These observations are consistent with our in vitro studies and may reflect the association of antiglycolytic action of BKIDC-1553 with antitumor activity. These results demonstrate that BKIDC-1553 has good systemic exposure and tissue penetration needed to achieve therapeutic levels. Furthermore, we tested organoid cultures derived from neuroendocrine prostate cancer that is androgen insensitive and resistant to AR-directed therapy. BKIDC-1553 elicited antiproliferative activities in LTL331R organoids derived from treatment-induced neuroendocrine prostate cancer xenograft model LTL331R(29) (**Figure 6C**). Taken together, these preclinical studies suggest BKIDC-1553 could be developed as a safe and effective therapy of advanced PCa.

**Figure 6.**
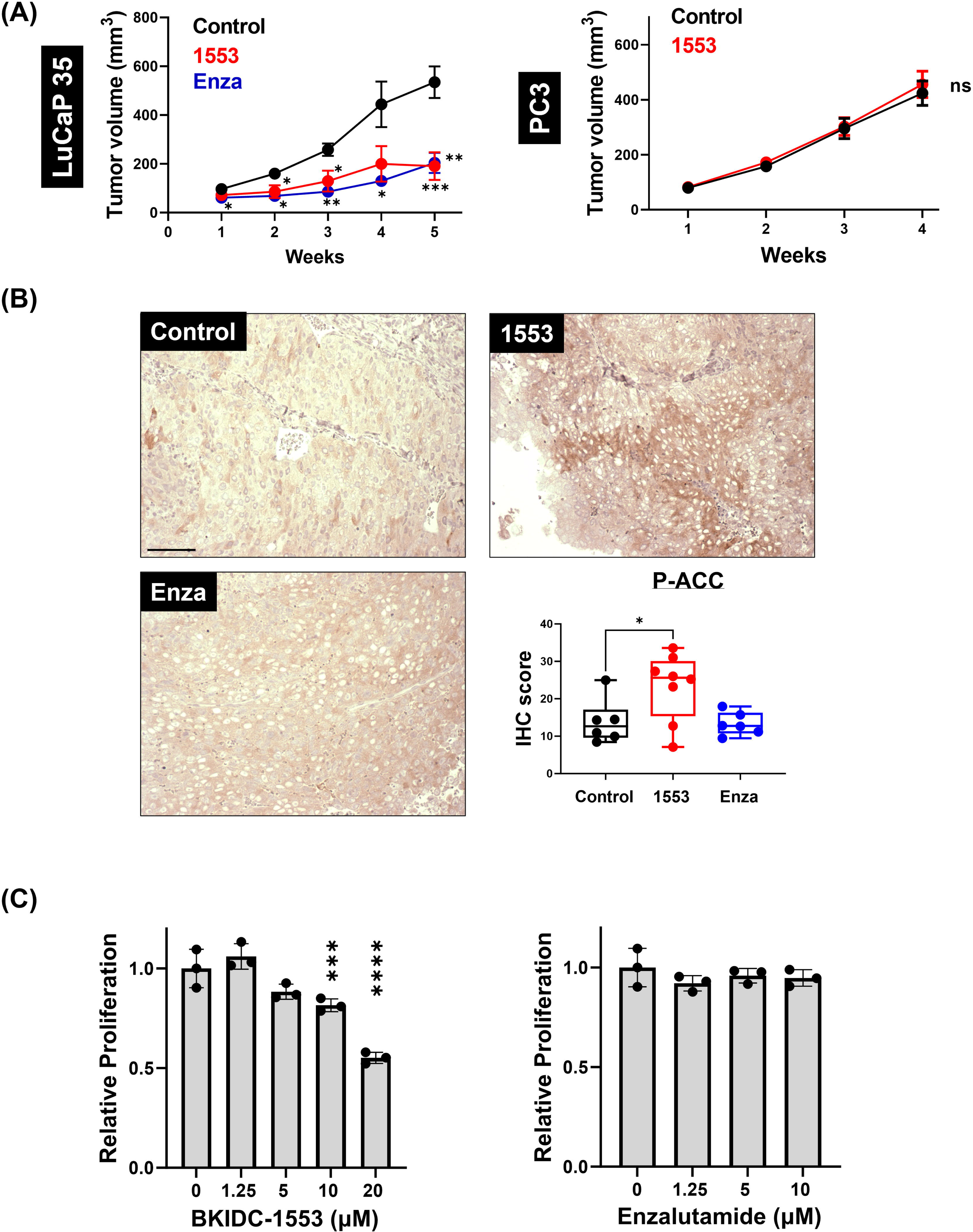
Preclinical anti-prostate cancer activity of BKIDC-1553. **(A)** Growth Curves of Prostate Cancer Xenografts Implanted in SCID Mice. The BKIDC-resistant human prostate cancer cell line PC3 and a human patient-derived xenograft (PDX) model (LuCaP 35) were injected subcutaneously in SCID mice. Mice were treated for 4-5 weeks with 20 mg/kg BKIDC-1553 po three times a week. The LuCaP 35 tumors responded to BKIDC-1553 to a similar degree as enzalutamide (Enza) and no decrease was seen in PC3 xenografts. The data are presented as mean ± standard error of mean (SEM) (n=7-10). One-way ANOVA followed by Tukey’s test for multiple comparison (LuCaP35) and two-tailed t tests (PC3): \**p* <0.05, \*\**p* <0.01, ns: not significant. **(B)** Immunohistochemical analysis of P-ACC in LuCaP 35 tumors. Representative images of LuCaP 35 tumors stained with an anti-P-ACC antibody are shown (at 20x magnification). Box and whisker plots show scoring analysis of anti-PACC immunoreactivities in tumors in the specified treatment. One-way ANOVA followed by Dunnett’s test for multiple comparison (vs. Control: *p <0.05). The number of animals used for IHC study was: 6 for control, 8 for 1553, 6 for enzalutamide. IHC score is defined in Materials and Methods. **(C)** Dose response of LTL331R line (patient-derived in vitro model of neuroendocrine prostate cancer) to BKIDC-1553 and enzalutamide using CellTiter-Glo® 3D Cell Viability Assay. The data shown are normalized to the values obtained from cells treated with DMSO vehicle and presented as mean ± SD (n=3). One-way ANOVA followed by Dunnett’s test for multiple comparison (vs. Control).

### BKIDC-1553 is an advanced preclinical candidate

BKIDC-1553 has the properties of a suitable advanced preclinical candidate in models of pharmacokinetics (PK), in vitro adsorption-distribution-metabolism-excretion (ADME), and toxicology in vitro and in vivo.

#### PK

We previously reported BKIDC-1553 (compound 32) has excellent PK parameters when it is administered in a single oral dose to mice, rats, calves, dogs and monkeys (35). We have reanalyzed these data, summarized in **Supplementary Data Table 6**, and demonstrate >65% oral bioavailability in three species and slow clearance in all species. The exposure obtained in each species is in accordance with allometric scaling, suggesting oral dosing will be successful in humans, and allometric scaling can be used to estimate the human dose for efficacy. To establish the efficacious exposure, we dosed SCID mice for 2 weeks with a dose that would allow for systemic exposures equivalent to concentrations successful in inhibiting human PDX LuCaP 35 in vitro, namely, 20 mg/kg BKIDC given orally on Monday, Wednesday, and Friday. Plasma samples were collected to assess for BKIDC levels. The AUC_0-288h_ (12 days) was estimated to be 3,190 μmol·h/L. The highest exposure over any 24 h period was estimated to be 381.8 μmol·h/L with a C_max_ of 19.4 μM, and these values were used to compare to the highest exposure over a 24 h period during toxicology experiments for determination of a safety index. BKIDC-1553 is predicted to have a ∼17 h half-life in humans based on an estimated Cl of 1.1 mL/min/kg and V of 1.6 L/kg in humans. Using the average BKIDC concentration (11.1 μM) and C_max_ (19.4 μM) associated with efficacy in the preclinical mouse prostate cancer model, the estimated efficacious oral dose in a 74 kg man will be 500 mg per day.

#### ADME

We previously reported that BKIDC-1553 has moderate solubility and oral absorption consistent with an oral preclinical candidate(35). BKIDC-1553 binds to human plasma proteins at 97%, which correlates with approximately 3% free fraction in plasma. Studies of permeability in Caco-2 cells and MDR1-MDCKII cells demonstrated high permeability (Caco-2 P_app A-B_: 61 ± 9.6 x 10^-6^ cm/s and P_app B-A_: 35 ± 1.0 x 10^-6^ cm/s; MDR1-MDCKII P_app A-B_: 15.6 ± 0.3 x 10^-6^ cm/s and P_app B-A_: 4.2 ± 0.6 x 10^-6^ cm/s) and no apparent efflux pump activity (Caco-2 efflux ratio: 0.6 ± 0.1; MDR1-MDCKII efflux ratio: 0.27 ± 0.03). Hepatocyte metabolism with human, rats, and dog microsomes demonstrated similar metabolites for all three species (**Figure S16**). Inhibition of cytochrome P450 (CYP) was assessed in vitro, and 20 μM BKIDC-1553 had no effect on CYPs 1A2, 2C9, 2C19, 2D6, or 3A4. In separate studies, 10 μM BKIDC-1553 resulted in <30% inhibition of CYP2B6, but did inhibit CYP2C8 by a mean of 56%. These data indicate that there is low potential for drug-drug-interactions associated with CYP 1A2, 2B6, 2C9, 2C19, 2D6, or 3A4. Follow up dose response studies are planned in order to determine the IC_50_ for BKIDC-1553 for CYP2C8 metabolism and better predict the possible effects on CYP2C8 drug metabolism before submission for Investigative New Drug (IND) status with the U.S. Food and Drug Administration (FDA).

#### Toxicology

As noted above (**Figure 1**), BKIDC-1553 has little to no effect on normal mammalian cells (HFF) in vitro. We previously reported BKIDC-1553 has no discernable effect on the cardiac risk channel, hERG, and an 80 human protein kinase screen revealed possible interaction only with PKD3 (IC_50_ of 0.12 μM), and MEK2 (IC_50_ of 0.97 μM)(35). BKIDC-1553 was not genotoxic in a bacterial mutagenicity assay (microAmes assay) or an in vitro micronucleus study. BKIDC-1553 was assessed at 10 μM for possible binding or inhibition to 44 receptors and enzymes in a Safety 44 Screen (https://cdnmedia.eurofins.com/corporate-eurofins/media/1069358/safetyscreen44_epdsfl420june16.pdf)(42). Of the 44 receptors and enzymes, BKIDC-1553 had activity in excess of 66% of control for only one target, Angiotensin Converting Enzyme (ACE), which was 100% inhibited at 10 μM. We described previously that mice tolerated a single 500 mg/kg oral dose and 5 days of daily dosing at 100 mg/kg using a polyetheleneglycol-ethanol-saline vehicle(35). We conducted an additional toxicology experiment for 7 days in mice with a lipid-based (Phosal 53 MCT) vehicle and found that mice tolerated 35 mg/kg/day and 70 mg/kg/day without clinical signs, CBC or biochemical changes, or histologic abnormalities, but did not tolerate 100 mg/kg/day, which led to clinically moribund animals after 3 days of dosing. Investigation of these mice revealed plasma glucose levels averaging >300 mg/dL, which presumably resulted in hyperglycemia. No other signs or sources of toxicity could be found in these animals. BKIDC-1553 blood exposures at this level were >5-times the exposure necessary for therapeutic effects on PDX, and thus a 5X margin of safety exists for therapy. Similar dose acceleration experiments were conducted in Beagle dogs treated with 1.7, 5, or 15 mg/kg BKIDC-1553 for 14 days, and no clinical, hematologic, or biochemical abnormalities were observed. However, the exposure (AUC_last 24 hours_) from the highest dose (15 mg/kg) was estimated to yield only 60% of exposure estimated in therapeutic dosing in mice. We subsequently investigated a dosing regimen of 30 mg/kg administered orally twice a day for 4 doses (2 days), followed by 15 mg/kg administered twice a day in a lipidic vehicle for 6 doses (3 days), which yielded exposures approximately equal to therapeutic exposures measured in mice. In this experiment, after administration of 2 doses of 30 mg/kg twice a day to dogs, two subsequent doses resulted in emesis, leading to the dose reduction to 15 mg/kg twice a day, noted above. Similar vomiting was not seen dogs treated with 5-day twice a day administration of vehicle alone. This suggests that vomiting was not due to the vehicle, and vomiting may limit dose escalation in dogs. Moreover, plasma glucose levels, serum chemistry, hematology, and coagulation and urinalysis were unremarkable. Advanced cardiac testing was performed in anesthetized rats with accelerating IV infusions of BKIDC-1553 to monitor drug induced changes in mean arterial blood pressure, heart rate, and cardiac contractility (**Table S7**). There were no significant changes in the cardiovascular parameters at IV doses of 3 mg/kg and 10 mg/kg but at 30 mg/kg there was a 25% increase in cardiac contractility. Plasma BKIDC-1553 levels at the 30 mg/kg were 55 µM, far surpassing the exposure necessary for therapy, thus the compound has a good cardiovascular safety index in rats. Altogether, in vivo assessments in mice, rats, and dogs demonstrate tolerability of BKIDC-1553 at doses that are pharmacologically active indicating potential therapeutic benefit to patients without drug-induced toxicity.

## DISCUSSION

Despite considerable efforts, precision oncology in advanced PCa has faced challenges, including drug resistance and limited populations who benefit from the related therapeutics(8). Metabolic reprogramming is observed in emergent CRPC after initial therapies fail(60). In particular, emergent CRPC becomes highly glycolytic as demonstrated by positive ^18^F-fluorodoxyglucose (^18^F-FDG) labeling in metastatic CRPC but not primary PCas(61–63). Strong avidity to FDG in advanced PCa suggests that targeting glycolysis would have a major impact on PCa progression and would be especially important in treating those tumors that currently are untreatable or have limited potential treatments(64).

However, the therapeutic targeting of metabolic complexes or enzymes in glycolysis is challenging, affecting not only tumor cells but also normal proliferating cells. Multiple enzymatic isoforms can also perform overlapping functions, and metabolic flexibility can divert processes away from inhibited pathways. Mammals possess five HK isoenzymes, including HK1, HK2, HK3, HK4/GCK1, and HKDC1 with overlapping and specific tissue and cellular distribution patterns to fix glucose as the first step in glycolysis(56). Along with catalytic domains, HK1 and HK2 contain N-terminal motifs to interact with VDAC1 for association with mitochondria and the glucose-6-phosphate (G6P)-binding sties for feedback allosteric inhibition(56). HK2 is required for oncogenic transformation and progression of some cancers, but HK2 is minimally expressed in most adult tissues and its systemic deletion in adult mice does not lead to overt side effects(22). Thus, HK2 garners much attention as an ideal and selective target in cancer therapeutics(65, 66). Target-based screening has identified HK2 selective inhibitors that competitively or allosterically inhibit catalytic functions or promote dissociation from the mitochondria, though they are still in early phases of clinical development(67).

As demonstrated here, using phenotypic screening and target deconvolution, we identified a series of compounds that fulfil a major unmet need for development of metabolism-based therapeutic interventions(15). It is clear from Seahorse ECAR measurements that BKIDCs interrupt glycolysis almost immediately after exposure in susceptible PCa cells. This is accompanied by a rapid reduction in glycolysis metabolites. Experiments showed that 2-DG accumulation, but not 3-OMG accumulation, was interrupted by BKIDCs. This demonstrated that hexokinase activity, but not glucose importation, was targeted by BKIDCs. Finally, knockout of HK2, but not HK1, abrogated the reduced ATP levels observed with BKIDCs combined with oligomycin. This demonstrated that BKIDCs interrupt ATP generation by glycolysis, and appear to do so in a HK2 dependent fashion. This makes sense, in that PCa is known to activate HK2 expression and become dependent on it for glycolysis(24).

We explored whether HK2 could be directly targeted by BKIDCs. BKIDC-1553 did not inhibit recombinant-HK2 (**Figure S13**). In addition to this, HK2 expression does not determine whether a cell line is sensitive. BKIDC-resistant cell lines PC3 and DU145 express HK2 protein (**Figure S14**). This suggests that BKIDCs do not act directly on HK2, but could act on the many molecules that are known to modulate HK2 function(58). We hypothesize the regulatory pathways of BKIDC-responsive prostate cancers are different from non-BKIDC responsive PC3 and DU145 cells, and other cell lines. This difference suppressing the HK2 function of susceptible PCa cells, leading to the suppression of glycolysis immediately after exposure of BKIDCs (Seahorse experiments, **Figure 3F**).

As noted, both HK2 and HK1 KO cell lines retained sensitivity to antiproliferative effects of BKIDCs. Observed antiproliferative phenotypes may reflect polypharmacological effects of BKIDCs. Indeed, it is still a matter of debate whether HK2 inhibition is sufficient to elicit antitumor activity(24, 68–71). HK1 and HK2 are dispensable in LNCaP cells as knockout of each gene did not affect cell proliferation and survival (this study) (69). However, metabolic reprogramming of the HK1 or HK2 KO population may change the coupling of glycolysis to antiproliferative effects. The lack of effect on proliferation may be due to BKIDC effects on other targets or may reflect the plasticity of PCa, allowing proliferation after complete KO of HK2(72, 73).

BKIDCs elicit cell-cycle arrest and AMPK activation, presumably due to the integrated stress response(74). Depending on the context of AMPK expression, e.g. spatial location in a cell, AMPK activation can be associated with a proliferation response(75). Indeed, AMPK may work together with HK2 inhibition in the BKIDCs’ effects on PCa. As we demonstrated here, BKIDCs synergize with OXPHOS inhibition (oligomycin) to promote cell death. A similar effect to BKIDCs could be seen with the non-specific glycolysis inhibitor, 2-DG. However, BKIDCs did not exhibit the toxicity to multiple cell lines that 2-DG demonstrates, making BKIDCs a good candidate for clinical therapy. HK2 inhibition reportedly renders therapeutic vulnerability to interference with OXPHOS or glutamine metabolism(76–78). This warrants exploration of synthetic lethal pairings for metabolism-targeted combined therapies to improve clinical results of BKIDC actions.

BKIDC-1553-N-Me and BKIDC-1553 were evaluated as possible leads to advance as preclinical candidates. BKIDC-1553-N-Me would be advantageous, potentially, through its lack of demonstrated “off-target” activity on protein kinases PKD3, RIPK2, and MEK2, and BKIDC-1553 has weak inhibitory activity towards these kinases. However, BKIDC-1553-N-Me was found to be rapidly converted to BKIDC-1553 after oral dosing in mice, and thus was not advantageous to BKIDC-1553 for oral administration.

BKIDC-1553 has the properties of a suitable advanced preclinical candidate with its ability to arrest prostate cancer xenograft growth in the SCID implantation model, as well as its PK, in vitro ADME, and toxicology attributes in vitro and in vivo. In the SCID xenograft LuCaP 35 model, 20 mg/kg of BKIDC-1553 delivered orally three times weekly was efficacious; even after 8 weeks of therapy, no toxicity was observed in the BKIDC-1553-treated animals, though vehicle controls with tumors needed to be euthanized per animal care committee regulations. The PK parameters of BKIDC-1553 have been examined in mice, rats, dogs, and monkeys, and excellent oral bioavailability has been found in all species tested. In vitro ADME testing supports that CYP metabolism likely leads to clearance of BKIDC-1553, but consistent with its slow clearance in multiple species by liver microsomes and hepatocytes, it is slowly cleared in all species tested in vitro. Moreover, BKIDC-1553 is highly bound to plasma protein (97% human plasma protein bound), which likely contributes to its slow clearance (only unbound drug is subject to hepatic clearance) and its safety (less free drug to contribute to safety issues in vivo). This slow clearance explains why such low doses (20 mg/kg), intermittently orally administered (3 times per week) in mice are successful for therapy of the PDX model. Comparative in vitro metabolism testing demonstrates that human, rat, and dog hepatocytes metabolize BKIDC-1553 to form identical metabolites, demonstrating that rats and dogs are excellent to study for final GLP-toxicology required for application for an IND status for BKIDC-1553. In vitro ADME studies suggest low probability of drug-drug interactions, perhaps with the exception of drugs that interact with CYP2C8, but this needs confirmatory dose response studies to adequately assess this risk. In vitro safety studies demonstrate lack of binding to hERG cardiac toxicity ion channels at 10 µM; and 10 µM is far beyond the unbound free levels measured during successful therapy. An 80 human kinase off-target screen revealed possible interaction only with PKD3 (IC_50_ of 0.12 μM), and MEK2 (IC_50_ of 0.97 μM)(35). However, these studies were carried out without ATP. ATP is a competitor for inhibition of BKIDC-1553. Therefore, given the high intracellular concentration of ATP (mM range) and high plasma protein binding of BKIDC-1553 (only free levels, ∼3% of plasma levels, should drive off-target effects), it is not likely that these protein kinases will be inhibited during therapy. BKIDC-1553 gave no signals for genotoxicity (microAmes test or in vitro micronucleus assays) or for interaction with a 44 off-target screen of receptors and enzymes of toxicological concern, except for 100% inhibition of angiotensin converting enzyme (ACE) at 10 µM. ACE inhibitors are generally considered safe, but follow on dose response experiments will be carried out to further explore this result prior to IND application. Finally, preliminary experiments in mice and dogs with short term, multiple day oral administration experiments demonstrate safety at observed therapeutic levels. Mouse experiments suggest a therapeutic window of about 5-fold, with hyperglycemia being the limiting toxicity. Dog experiments suggest vomiting could be an issue at exposures above therapeutic exposures. Both hyperglycemia and vomiting are toxicity issues that can be monitored and managed easily in human clinical trials, and during therapy after regulatory approval. A pre-IND assessment with the U.S. FDA suggests that pending a final Good Laboratory Practice (GLP) Toxicology and Chemistry Manufacturing and Controls (CMC) development study, BKIDC-1553 is on track for a successful IND application. Thus, we conclude BKIDC-1553 is an outstanding preclinical candidate for development against prostate cancer.

In summary, we overcame the hurdle of targeting tumor metabolism by developing a series of druggable small molecules, BKIDCs, that elicit antiproliferative effects selectively in PCa models. BKIDCs rapidly shut down glycolytic energy production, an effect that appears dependent on HK2 expression. BKIDC-1553 was effective in vitro and in vivo in shutting down prostate cancer growth, and demonstrates outstanding PK, ADME, and safety properties consistent with a late preclinical lead. Only CMC/Formulation/Stability and GLP toxicology studies remain before an IND status can be requested from the U.S. FDA, to advance this promising molecule to clinical trials of advanced prostate cancer. Since this compound has a unique mechanism of action, it will be a welcome addition to the chemotherapy of advanced prostate cancer, which generally becomes resistant to the available androgen-directed therapy, and may be an addition or alternative to rescue cytotoxic chemotherapy regimens that are very toxic and fail to deliver lasting response.

## Abbreviations

AA: antimycin A
ACC: acetyl-CoA Carboxylase
ACN: acetonitrile
adeno-PCa: adenocarcinoma prostate cancer
ADME: adsorption-distribution-metabolism-excretion
AMPK: AMP-activated protein kinase
AR: androgen receptor
ARSi: androgen receptor signaling inhibitors
AUC: area under the curve
BKI: bumped kinase inhibitor
BKIDCs: bumped kinase inhibitor (BKI) derived compounds
BSA: bovine serum albumin
CAR: cell-associated radioactivity
Cl: clearance
CMC: Chemistry Manufacturing and Controls
CRPC: castration-resistant prostate cancer
CYP: cytochrome P450
CytB: cytochalasin B
DAPI: 4, 6-diamidino-2-phenylindole
DAB: 3,3’-diaminobenzidine
2-DG: 2-deoxyglucose
2-DG-6-P: 2-DG-6-phosphate
DHAP: dihydroxyacetone phosphate
DPBS: Dulbecco’s phosphate-buffered saline
ECAR: extracellular acidification rate
FBS: fetal bovine serum
FCCP: carbonyl cyanide 4-(trifluoromethoxy)phenylhydrazone
FDA: Food and Drug Administration
FDG: fluorodoxyglucose
GLP: Good Laboratory Practice
G6P: glucose-6-phosphate
hERG: human ether-a-go-go related gene
HK: hexokinase
hr: human recombinant
IC_50_: 50% inhibition values
IND: Investigative New Drug
IRS-1: insulin receptor substrate-1
ITS: Insulin-Transferrin-Selenium
KO: knock-out
KRB: Krebs-Ringer bicarbonate solution
LC-MS: liquid chromatography coupled with mass spectrometry
MTS: 3-(4,5-dimethylthiazol-2-yl)-5-(3-carboxymethoxyphenyl)-2-(4-sulfophenyl)-2H-tetrazolium, inner salt
NE: neuroendocrine
OCR: oxygen consumption rate
Oligo: oligomycin
3-OMG: 3-O-methyl D-glucose
OXPHOS: oxidative phosphorylation
OXPHOSi: OXPHOS inhibitors
P_app_: apparent permeability coefficient
PARP: poly ADP ribose polymerase
PCa: prostate cancer
PDL: poly-D-lysine
PDX: patient-derived xenograft
PEP: phosphoenolpyruvate
PG: phosphoglycerate
PK: pharmacokinetics
PKM2: pyruvate kinase M2
PP: pyrazolopyrimidine
PrP: pyrrolopyrimidine
RLU: relative luminescence units
RPPA: reverse-phase protein arrays
RPS6: ribosomal S6 protein
SCID: severe combined immunodeficient
SD: standard deviation
SEM: standard error of mean
SO: sodium oxamate
UPLC-HRMS: ultra-high-performance liquid chromatography-high resolution mass spectrometry
V: volume of distribution

## Acknowledgements

We thank Dr. Taran Gujral and Stella Shin (Fred Hutchinson Cancer Center) for help with the RPPA data, Michael S. Dey PhD and D. Bruce Burlington PhD for their consultation, Dr. Michael Schweizer for discussion and manuscript editing, and Mika Munari for technical assistance. We are grateful to Prof. Susan Charman and the team within the Centre for Drug Candidate Optimisation, Monash Institute of Pharmaceutical Sciences, Monash University, Parkville, Australia for testing the ADME properties of BKIDC-1553. We are grateful to Medicines for Malaria Venture (MMV) for providing access to testing for ADME for BKIDC-1553. Funding: Lopker Family Foundation in honor of Karl Lopker, who died of prostate cancer in 2018; WE-REACH, supported by University of Washington and National Institutes of Health grant 5U01HL152401; US Department of Defense grant W81XWH-17-1-0323 (SRP, DM, WCVV); US Department of Defense grant W81XWH-20-1-0146 (SRP); US Department of Veterans Affairs Merit Review grant I01BX0033248 (SRP); US Department of Veterans Affairs Service Directed Project number SDR-PLYMATE (SRP); National Institutes of Health/National Cancer Institute grant CA255830 (TU) and National Institutes of Health/National Cancer Institute grant P50CA097186 (KKO).

## Notes

**Conflict of interest disclosure statement:** SRP is the President, WCVV is the Chief Executive Officer, DJM is the Chief Scientific Officer, and KKO is the Secretary of ProsTech Inc. ProsTech Inc. was created to commercialize therapies for prostate cancer, but as yet, has no commercial claims on BKIDCs or any other potential commercial products. Other authors declare that they have no competing interests. The patent numbers related to this article are as follows. Use Patent in Cancer Field: US 10350211 issued, Composition of Matter Patents: US 10632122 and US 10307425 both issued; and US patent application 17/758,577 pending. None of these four are sublicensed for cancer or producing royalties.

### Competing Interest Statement

SRP is the President, WCVV is the Chief Executive Officer, DJM is the Chief Scientific Officer, and KKO is the Secretary of ProsTech Inc. ProsTech Inc. was created to commercialize therapies for prostate cancer, but as yet, has no commercial claims on BKIDCs or any other potential commercial products. Other authors declare that they have no competing interests. The patent numbers related to this article are as follows. Use Patent in Cancer Field: US 10350211 issued, Composition of Matter Patents: US 10632122 and US 10307425 both issued; and US patent application 17/758,577 pending. None of these four are sublicensed for cancer or producing royalties.

### Summary of Updates

This revision has additional data, including updated version of Figure 6 (organoid cultures, immunohistochemistry), Supplemental Figure S3 (crystal violet assay), Supplemental Figure S4C (cell cycle profile), and Supplemental Figure S14 (data mining).

