## Supplemental information for "A Compound that Inhibits Glycolysis in Prostate Cancer Controls Growth of Advanced Prostate Cancer"

**Supplementary Materials**

**Supplementary Figures**

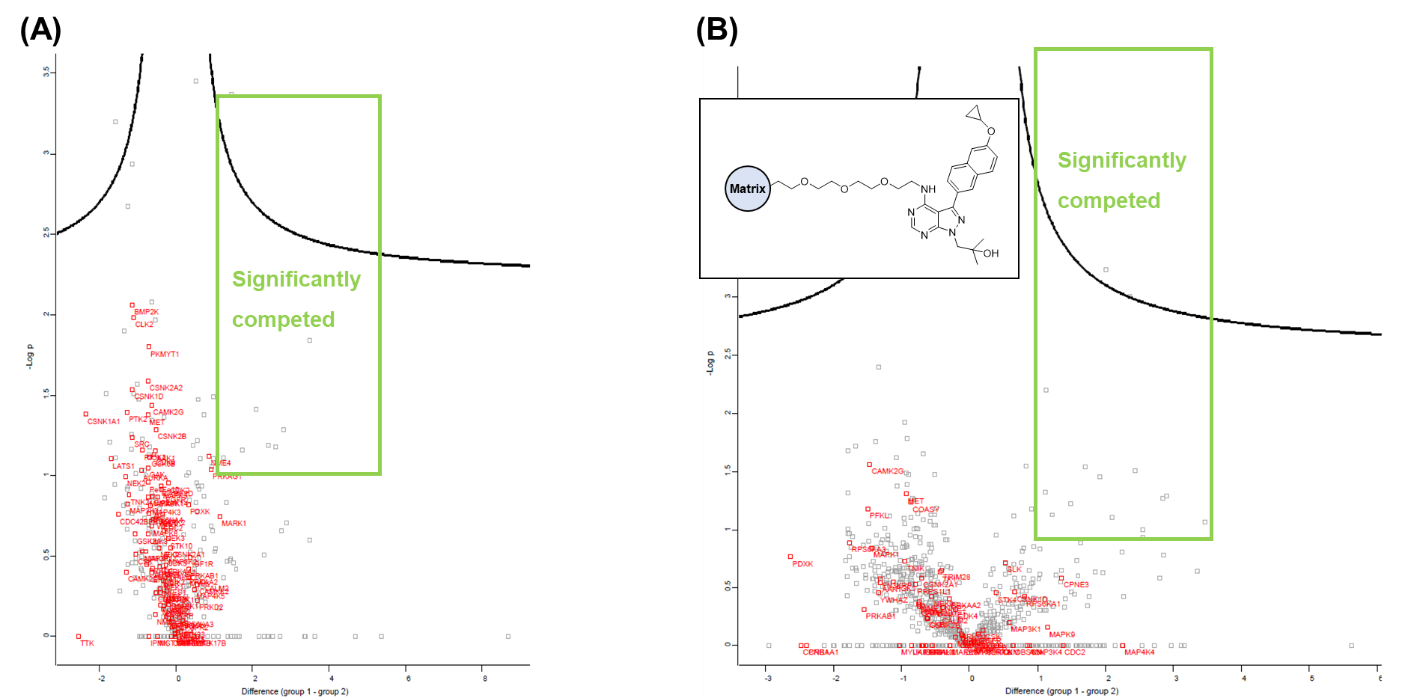

**Figure S1. Competitive proteomic profiling of 1553-N-Me. (A)** Kinobead profiling of 1553-N-Me. 20 µM of 1553-N-Me or DMSO was incubated with a HEK293/HCT116 lysate mixture and then subjected to a competitive kinobead matrix pulldown. Enriched proteins were quantified by LCMS and significantly competed (Log2Difference (1553-N-Me compared to DMSO) > 1 and -LogP value >1) targets were identified. Of the ~120 proteins with kinase activity that were identified, none (including RIPK2 and PRKD2) were significantly competed by 1553-N-Me. Pulldowns and mass spectrometric quantification were performed in quadruplicate. Significantly competed proteins were within the rectangle surrounded by green line. **(B)** Competitive profiling of 1553-N-Me with an immobilized 1826 (structure shown in the inset) matrix. 20 µM of 1553-N-Me or DMSO was incubated with a HEK293/HCT116 lysate mixture and then subjected to a competitive pulldown with immobilized 1826. Enriched proteins were quantified by LCMS and significantly competed (Log2Difference (1553-N-Me compared to DMSO) > 1 and -LogP value >1) targets were identified. No protein kinases were significantly competed by 1553-N-Me. Pulldowns and mass spectrometric quantification were performed in quadruplicate. (n=4). Significantly competed proteins were within the rectangle surrounded by green line.

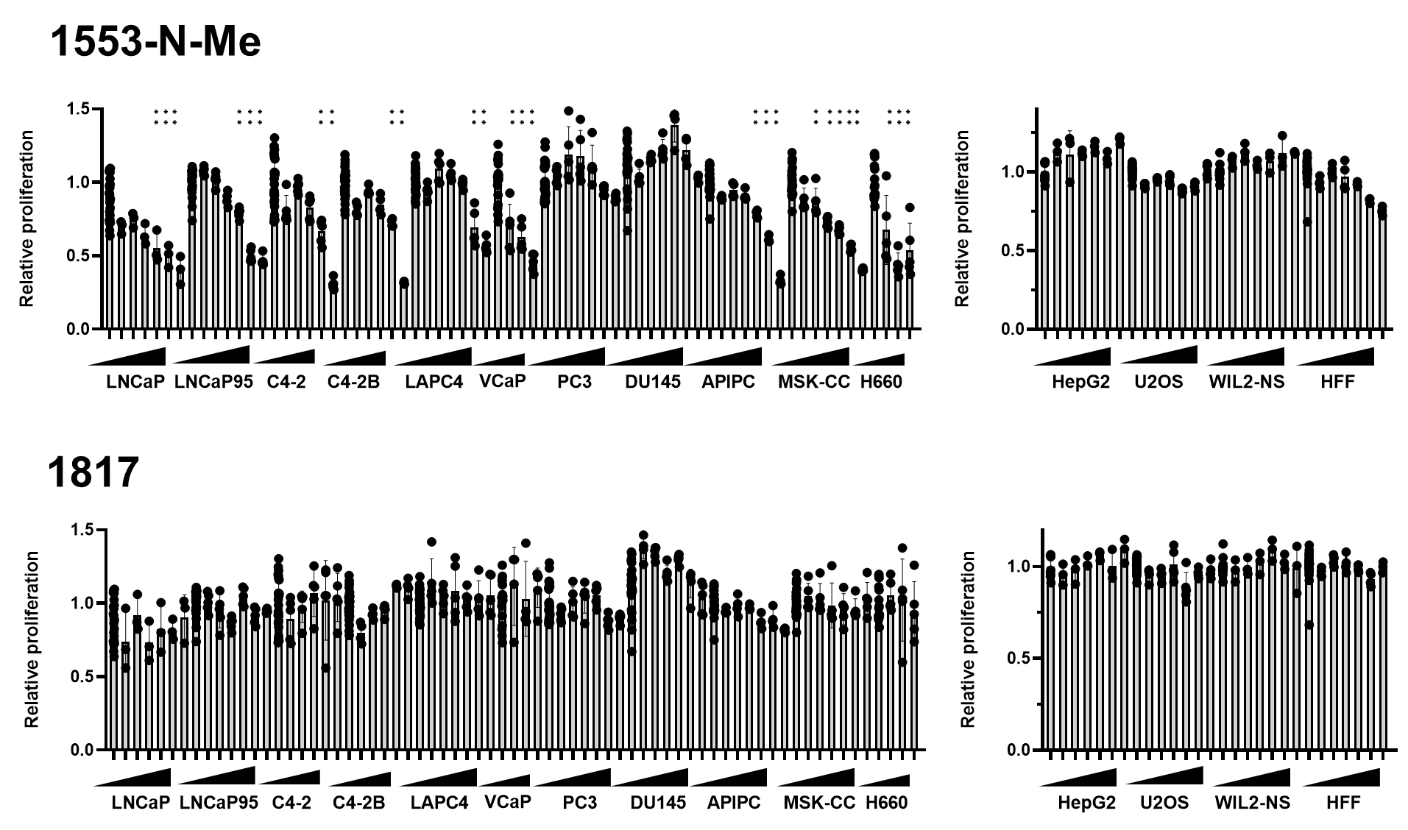

**Figure S2. Dose response of multiple cancer cell lines to BKIDC-1553-N-Me and 1817 using the MTS proliferation assay**. The effect of BKIDC on cell proliferation was evaluated in the same manner as described in **Figure 1**. One-way ANOVA followed by Dunnett’s test for multiple comparison: ***p* <0.01.

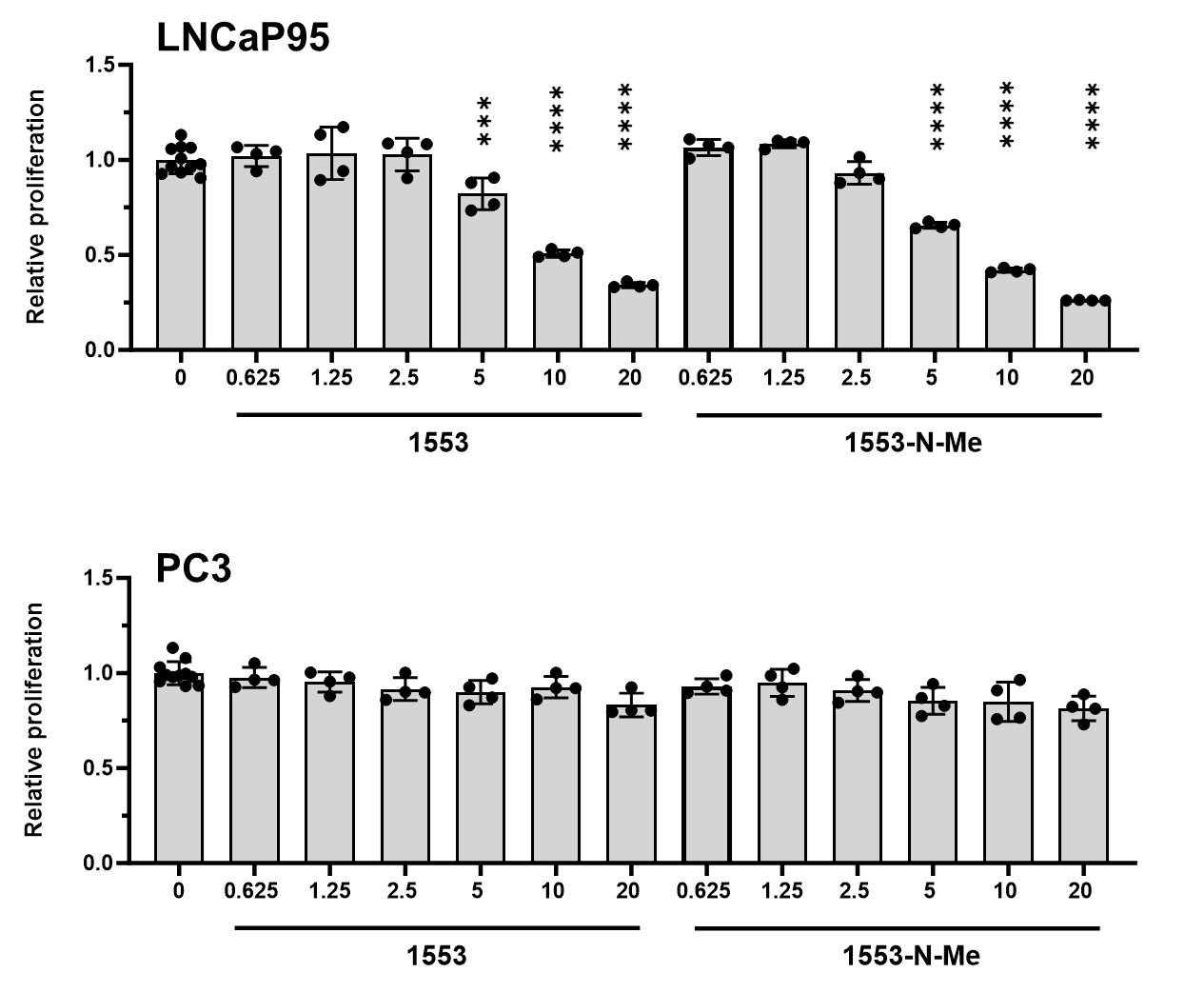

**Figure S3. Crystal violet-based cell proliferation assay**

Cell proliferation was evaluated by crystal violet assay at 72 h after BKIDC-1553 or 1553-N-Me was added at the range between 0.625-20 µM (DMSO as a vehicle control). The data shown are representative of at least three independent experiments, normalized to the values obtained from cells grown in medium containing 0.1% DMSO vehicle and presented as mean ± standard deviation (SD) (n=3-5). One-way ANOVA followed by Dunnett's test for multiple comparison (vs. Control). Note crystal violet assay, which is insensitive to changes in cell metabolic activity, provided the similar results as MTS assay (**Figure 1**, **Figure S2**) to reflect antiproliferative activity of BKIDC in LNCaP95 but not PC3.

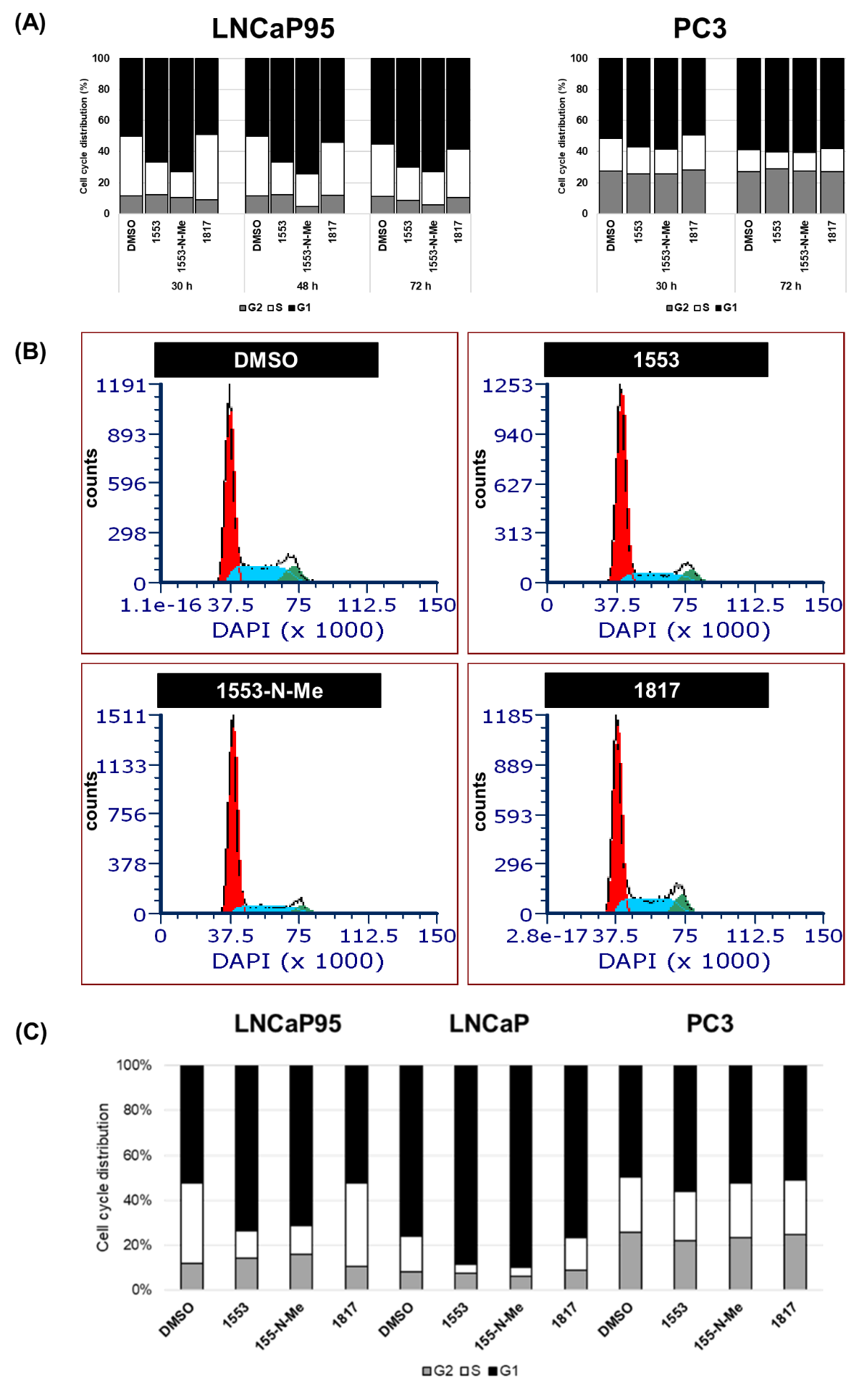

**Figure S4. Cell cycle profiles and histograms of DAPI-stained LNCaP95 and PC3 treated with BKIDCs** **(A)** Cell cycle profiles of LNCaP95 and PC3 treated with BKIDCs for 30, 48, and 72 h. The same profile data used for **Figure 2** are shown for cells treated for 30 h. Alterations in cell cycle distribution were maintained over 72 h for LNCaP95 cells. There were no differences in cell cycle distribution between control (DMSO and 1817) and treatment (1553 and 1553-N-Me) for PC3 cells at early and later time points. **(B)** Histograms of 4',6-diamidino-2-phenylindole (DAPI)-stained LNCaP95 cells treated with BKIDC for 72 h. A lack of sub-G1 peaks indicates no sign of apoptotic cell death in cells treated with antiproliferative BKIDC-1553 and 1553-N-Me. The number of cells located in S phase is reduced following treatment with BKIDC-1553 or 1553-N-Me compared with DMSO or BKIDC-1817 (negative control). Red: G1 phase, light blue: S phase, green: G2 phase. **(C)** Reproducibility of **Figure 2A** result was confirmed by another independent experiment. The percentage of cells at G1, S, and G2 phase of cell cycle at 30 h after treatment with the indicated BKIDCs. Cell cycle analysis showing G1 cell cycle arrest in LNCaP and LNCaP95 cells compared to PC3 when treated with BKIDC-1553 and 1553-N-Me.

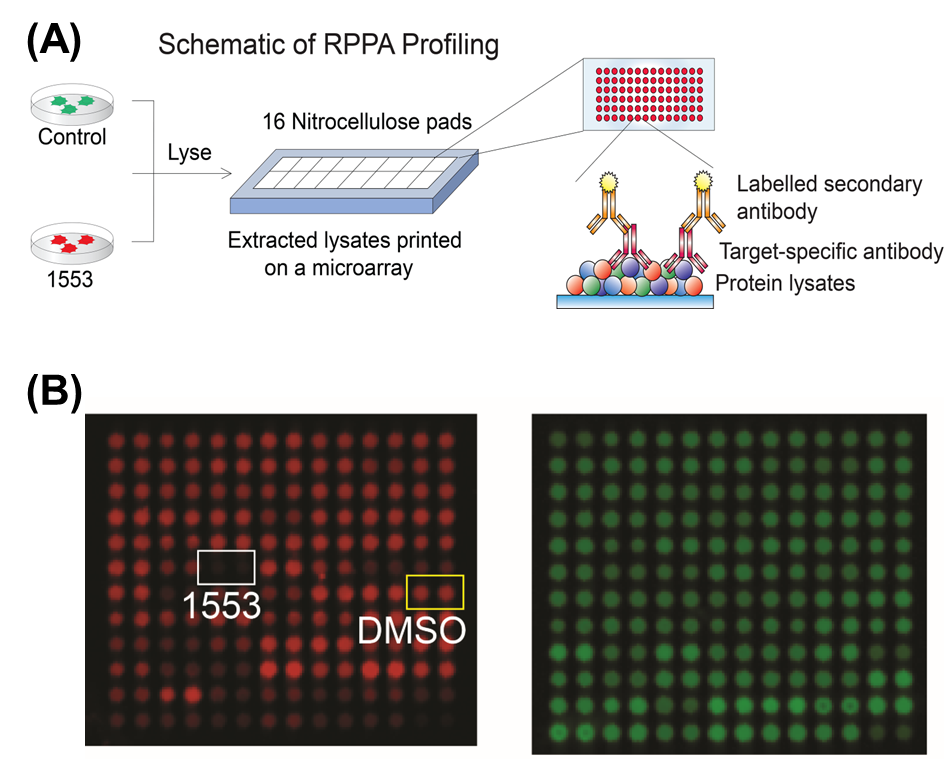

**Figure S5. RPPA Profiling (A)** A schematic of RPPA profiling showing that lysates from inhibitor treated cells are printed onto nitrocellulose covered microarrays and probed with a panel of validated antibodies (see **Table S3**). **(B)** Representative images of a lysate-printed microarray probed with an antibody to phosphorylated RPS6 at S235/S236 (left) and β-actin (right) as the total protein control.

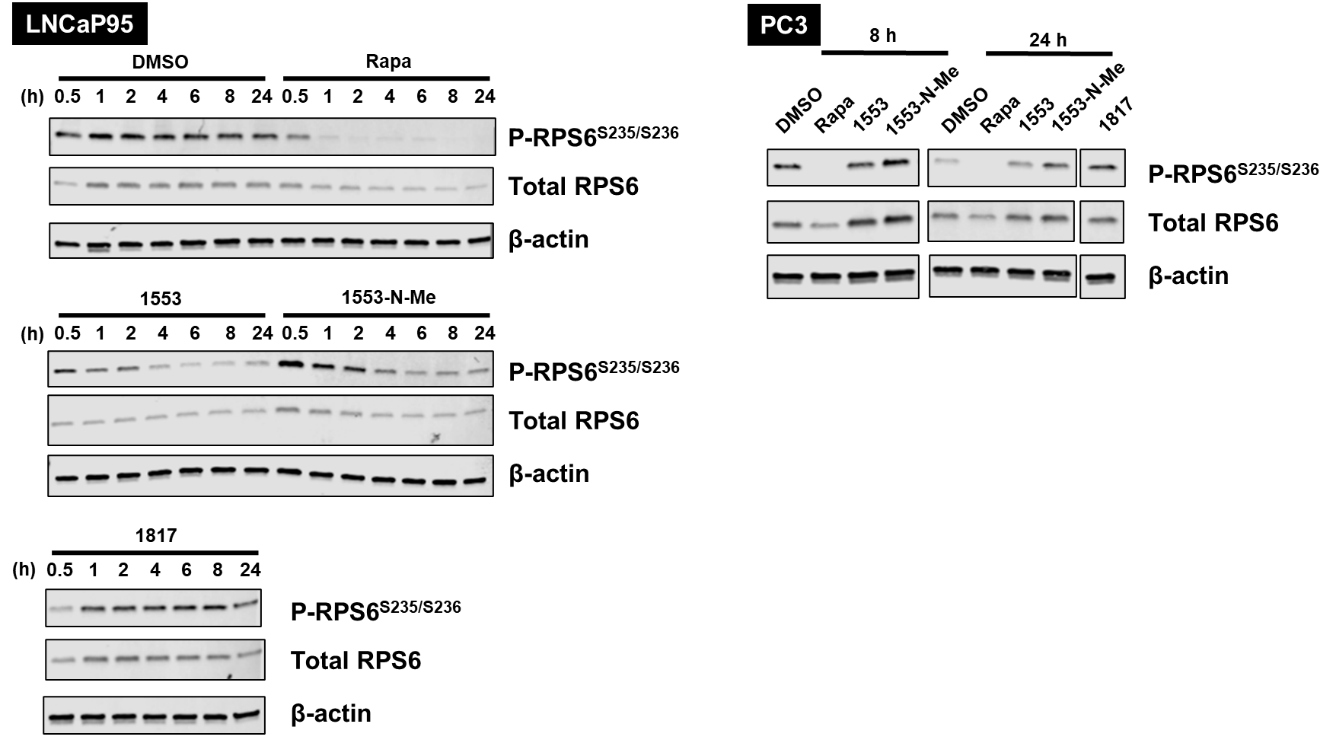

**Figure S6. Western blot analysis confirms the findings with RPPA on susceptible cells treated with BKIDCs.** LNCaP95 and PC3 were treated with BKIDC at 20 µM or mTORC1 inhibitor rapamycin (Rapa) at 50 nM for the indicated time. Phosphorylated RPS6 at S235/S236 was decreased within 4 h in LNCaP95 cells after treatment with BKIDC-1553 and 1553-N-Me while it was unchanged in PC3. BKIDC-1817 and Rapa served as negative and positive control in both cell lines, respectively.

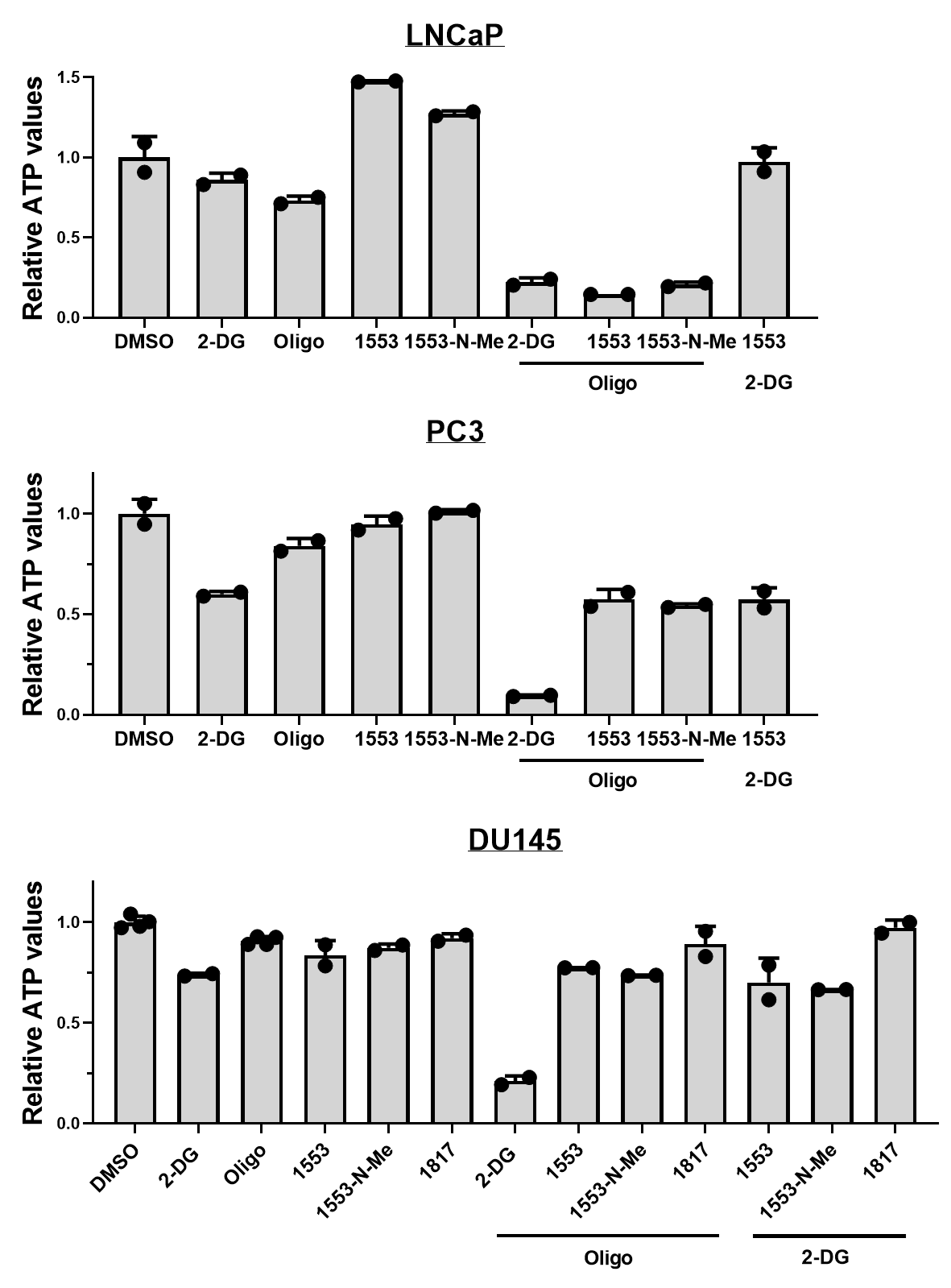

**Figure S7. Effects of BKIDC-1553 and 1553-N-Me on intracellular ATP levels in LNCaP, PC3, and DU145 cells**. Cells were treated with BKIDC, 2-DG, and oligomycin (Oligo) singly or in combination for 120 min. Intracellular ATP levels were evaluated with the Celltiter-Glo™ luminescent assay kit. The data shown are normalized to the values obtained from cells grown in medium containing 0.1% DMSO vehicle and presented as mean ± SD (n=2). Note PC3 and DU145 were resistant to decreasing ATP when co-treated with Oligo and antiproliferative BKIDC while they were sensitive to 2-DG.

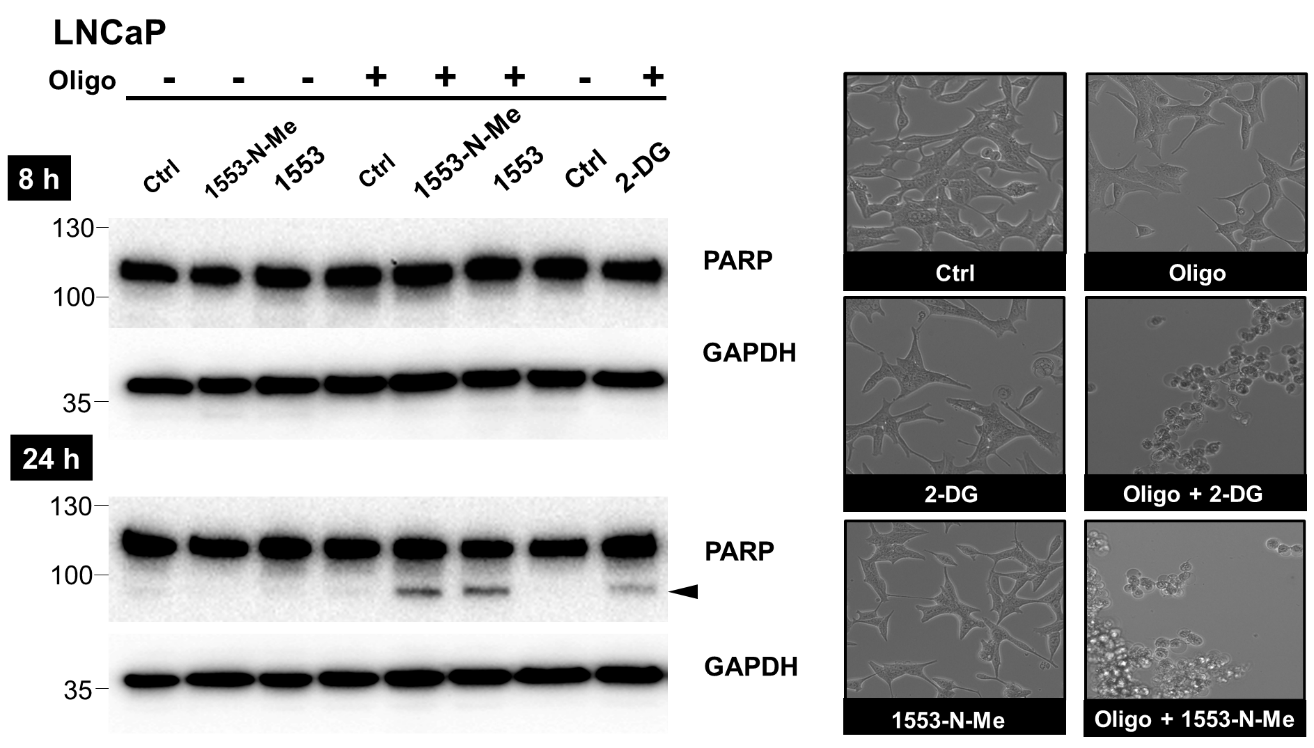

**Figure S8. BKIDCs cooperate with oligomycin to induce apoptotic cell death.** LNCaP cells were treated with BKIDCs, 2-DG, and oligomycin (Oligo) singly or in combination for 8 and 24 h. BKIDCs and 2-DG cooperated with Oligo to promote cleavage of PARP at 24 h but not 8 h-treatment in LNCaP. This biochemical event was reflected by appearances of detached cells in the phase-contrast microscopic images.

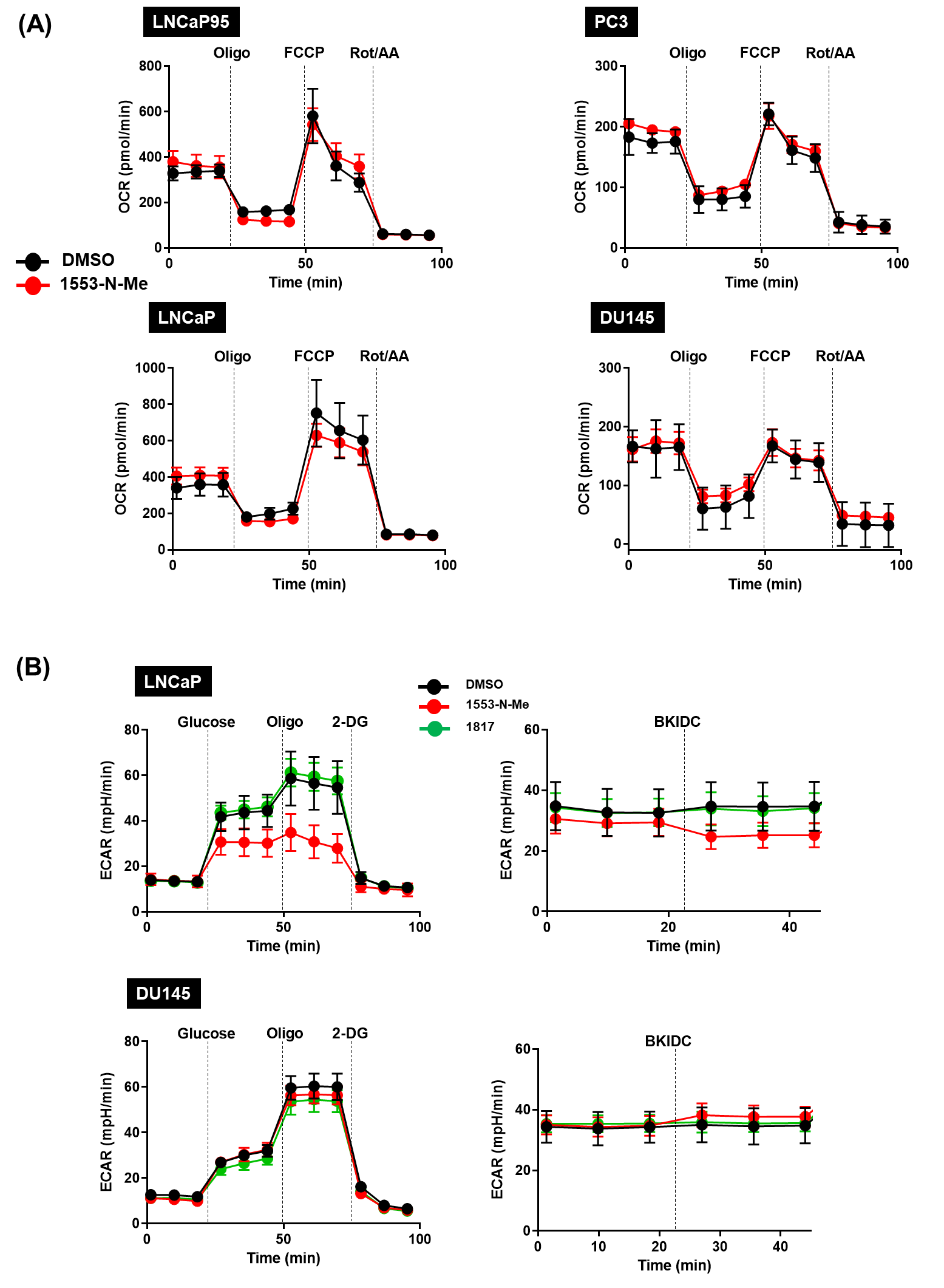

**Figure S9. Measure of the oxygen consumption rate (OCR) and** **extracellular acidification rate (ECAR) using the Seahorse assay (A)** Mitochondrial respiration was evaluated by the Seahorse XFe24 analyzer. Profiles of oxygen consumption rate (OCR) across time are shown for LNCaP95, LNCaP, PC3, and DU145. Arrows indicate the addition of OXPHOSi (oligomycin; FCCP; rotenone/antimycin A). OCR profiles were similar between DMSO and BKIDC-1553-N-Me treated cell lines for all four cell types. **(B)** ECAR was determined in LNCaP and DU145 cells in the same manner as described in **Figure 3**.

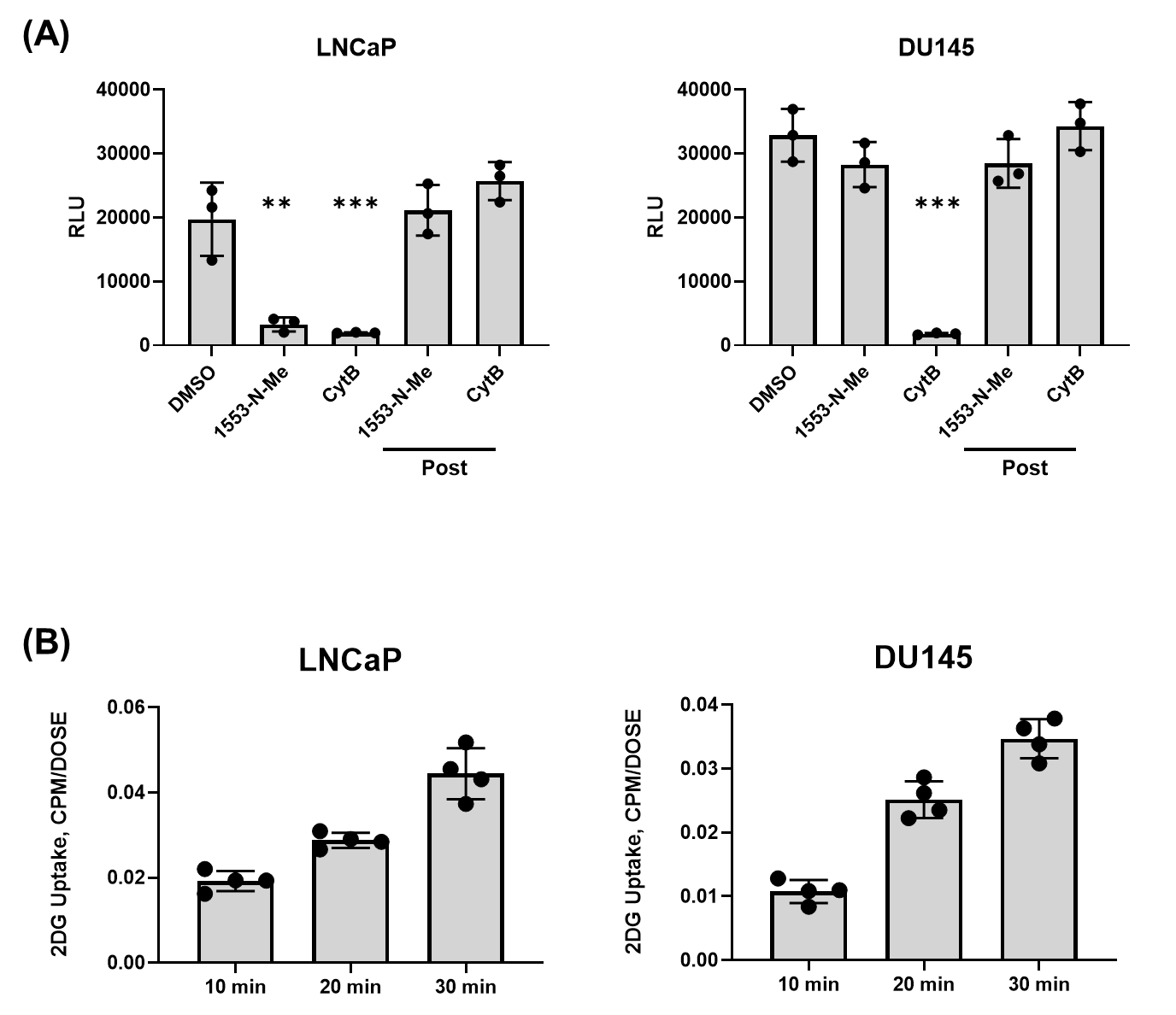
**Figure S10. Glucose uptake assays. (A)** Glucose Uptake-Glo Assay^TM^ (Promega) was used to determine if glucose import and phosphorylation was affected by BKIDC-1553-N-Me. The assay is based on the uptake of 2-DG and the enzymatic detection of intracellular accumulation of 2-DG-6-phosphate. Cytochalasin B (CytB: 50 µM) is a positive control that inhibits glucose uptake into cells and inhibited signal (RLU: relative luminescence unit) in both LNCaP and DU145 cell lines. In contrast, the addition of 20 μM BKIDC-1553-N-Me inhibited signal only in LNCaP, but not DU145 cells. Post BKIDC-1553-N-Me and Post CytB are controls where the compounds were added after cellular harvest, to demonstrate these compounds did not interfere with the assay to detect 2-DG-6-phosphate. One-way ANOVA followed by Tukey’s test for multiple comparison: ***p* <0.01, ****p* <0.001. **(B)** Time-dependent accumulation of 2-DG-derived signal in both LNCaP and DU145.

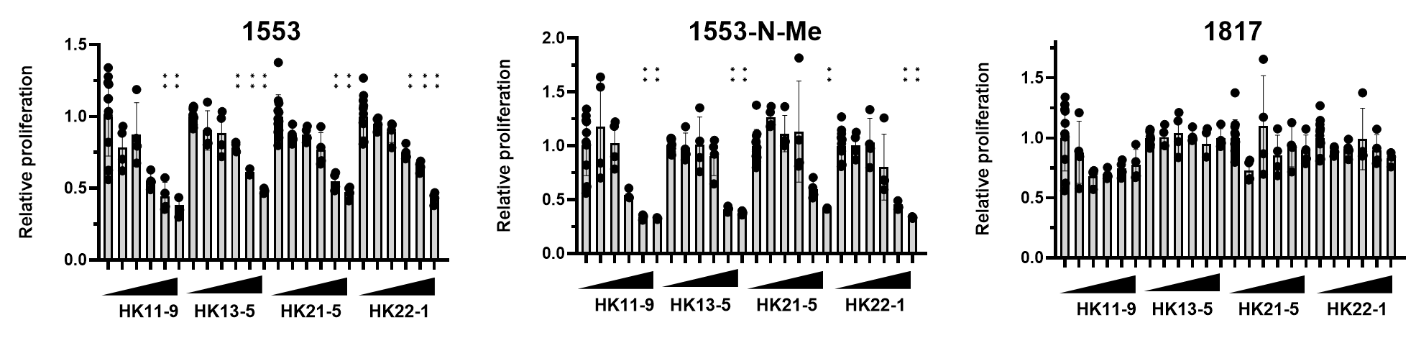

**Figure S11. Antiproliferative activity of BKIDCs on HK1 and HK2 KO LNCaP cell lines.** Cell proliferation was evaluated by MTS assay at 72 h after BKIDC was added at the range between 1.25-20 µM. DMSO was used as a vehicle control. The data shown are normalized to the values obtained from cells grown in medium containing 0.1% DMSO vehicle and presented as mean ± standard deviation (SD) (n=4). One-way ANOVA followed by Dunnett’s test for multiple comparison: ***p* <0.01.

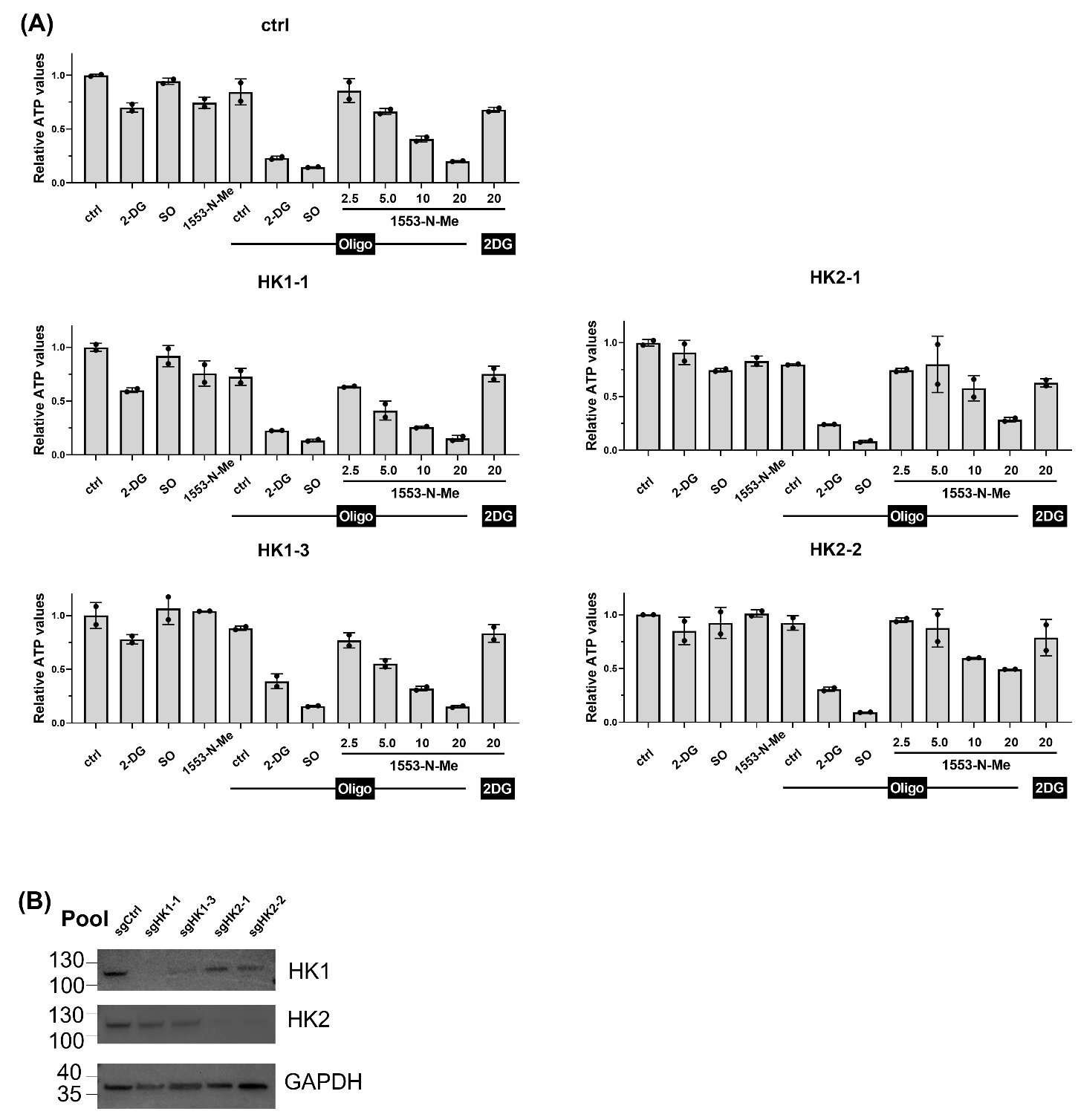

**Figure S12. Effects of BKIDCs on intracellular ATP levels in HK1 and HK2 KO LNCaP cell lines.** **(A)** Pooled LNCaP knockout lines with different sgRNAs displayed resistance to BKIDC and oligomycin only when HK2 is knocked out, but not with HK1 knock out. Note all lines were sensitive to 2-DG and sodium oxamate (SO). **(B)** Western blot validation of reduced levels of HK1 and HK2 in the corresponding LNCaP KO pools. GAPDH serves as a loading control.

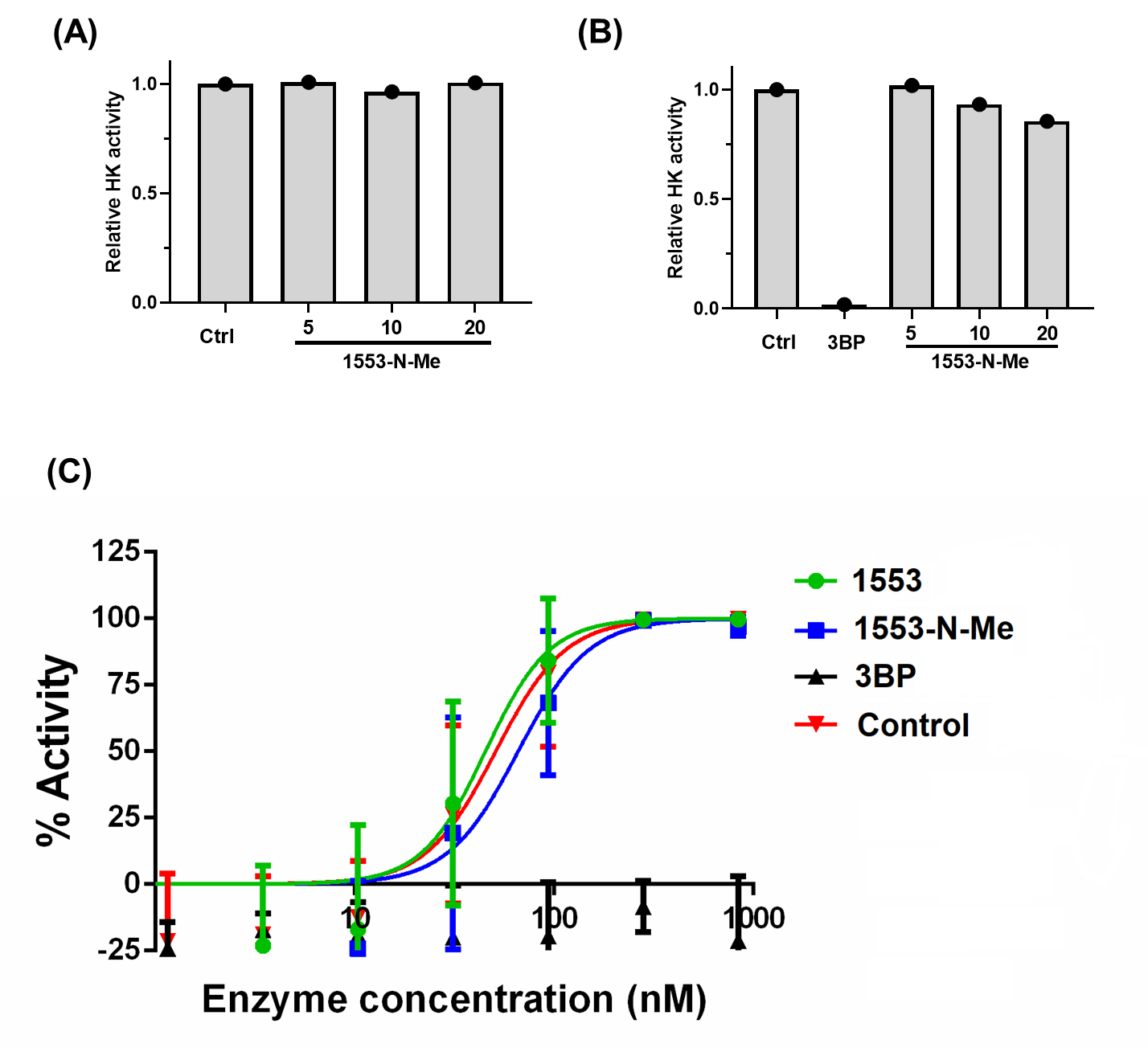

**Figure S13. Evaluation of Hexokinase activity using LNCaP crude extracts and recombinant enzyme. (A)** Hexokinase activity was assayed in crude extracts from subconfluent LNCaP cells in the presence and absence of BKIDC-1553-N-Me with the Hexokinase Colorimetric Assay Kit according to the manufacturer’s instructions. Hexokinase activity was normalized to the value from the assay with DMSO as a vehicle control. **(B)** Hexokinase activity was assayed with 11.7 ng of recombinant human HK2 protein in the presence of BKIDC-1553-N-Me or 10 µM 3-bromopyruvate (3BP), a positive control known inhibitor of hexokinase. **(C)** The activity of HK2 was measured with or without the indicated compound (10 µM for 1553 and 1553-N-Me, 100 µM for 3BP) using a serial dilution of the enzyme in the range between 70 and 840 nM. Activities are expressed as percentage of those with the control reaction containing 840 nM HK2 and shown as mean ± SD. 3BP inhibited 100% at every concentration.

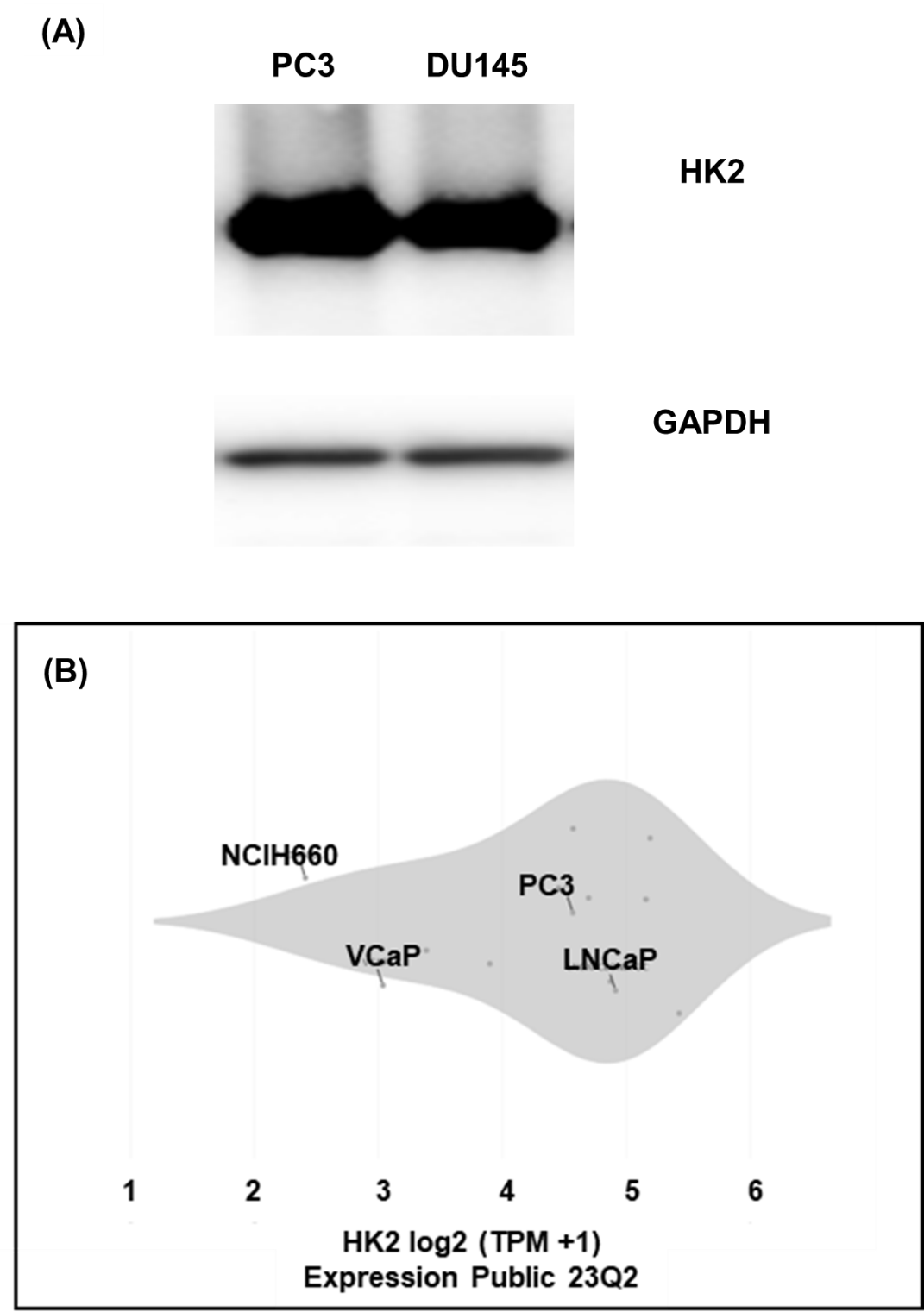
**Figure S14. Expression of HK2 in prostate cancer cell lines.**

**(A)** Western blot analysis to demonstrate that PC3 and DU145 express HK2. **(B)** Expression levels of HK2 in prostate cancer lines used in this study. The RNA-seq data were extracted from publicly available database DepMap portal (<https://depmap.org/portal/interactive/?x=slice%2Fexpression%2F11218%2Fentity_id>).

Selected cell lines are shown to represent the non-responder (PC3) and responder lines (NCI-H660, VCaP, and LNCaP). TPM: transcription per million

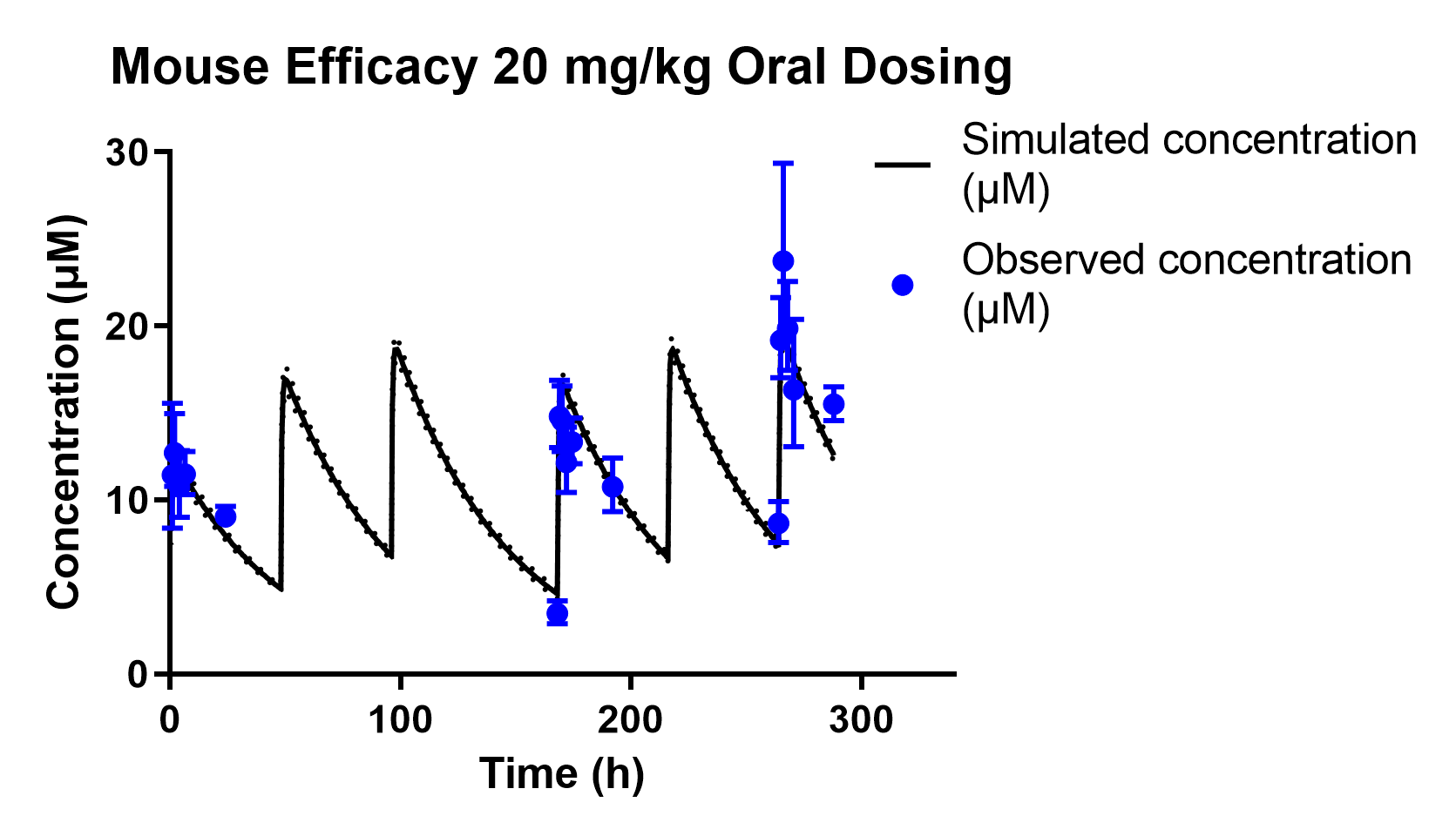

| **Average AUC last day (µmol*h/L)** | **Average AUC 0-12d (µmol*h/L)** |
| --- | --- |
| **381.8** | **3,190.0** |

**Figure S15. Measured and simulated plasma levels in SCID mice with 20 mg/kg oral dosing three times weekly document exposures in plasma obtained for therapeutic growth arrest of prostate cancer in vivo.** Three SCID mice, without tumors implanted, were dosed with BKIDC-1553 by oral gavage on Monday, Wednesday and Friday for two weeks, and plasma was obtained at the time points noted. Plasma BKIDC-1553 levels were obtained by LC-MS/MS and the mean plotted as micromoles per liter (µM) and the standard deviation of the mean as error bars. AUC last day refers to AUC over the last dosing day, 264-288 h, and AUC 0-12d refers to AUC over the study period (0-288 h).

**Figure S16**. Metabolites of BKIDC-1553 produced by liver microsomes from rats, dogs, and humans. The metabolite scheme above depicts the potential relationships between the various metabolites observed. These proposed interrelationships are based only on inference. BKIDC-1553 exhibited low turnover in rat, dog, and human liver microsome incubations. The major metabolic pathways were O-dealkylation (M1) and mono-oxidation (M2, M3, and M5). Other metabolic pathways were di-oxidation and hydrogenation (M6), di-oxidation and dehydrogenation (M4). All metabolites detected in human were also detected in rat and dog.

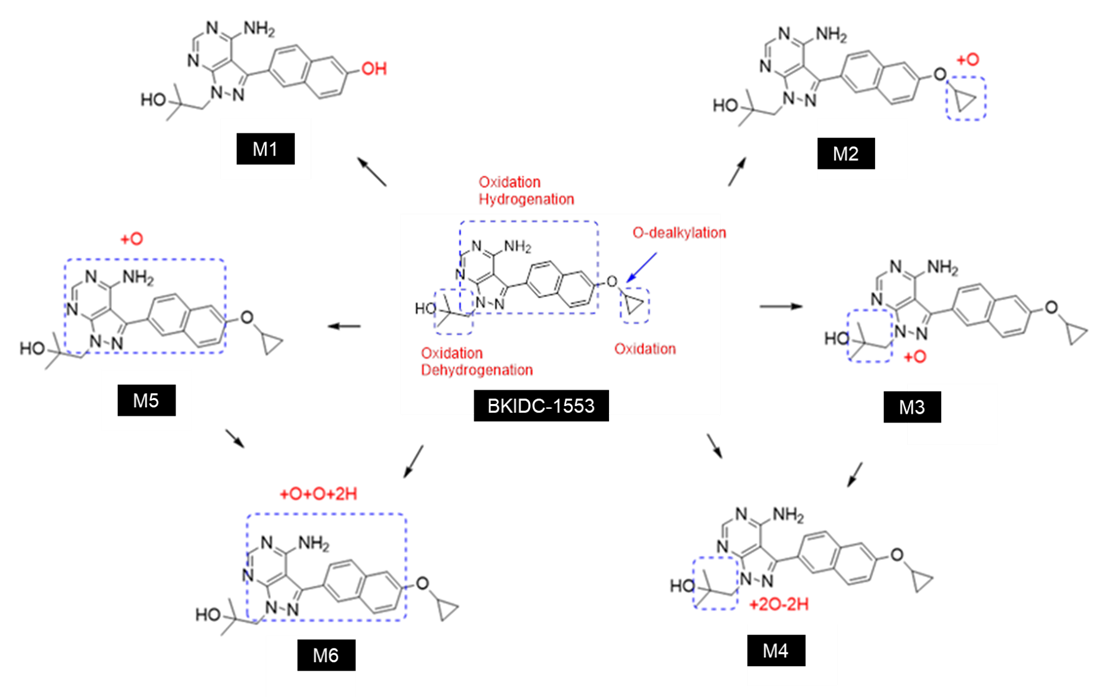

**Supplementary Tables**

**Table S1. PrP and PP-based BKIDC chemical structures.**

| **BKIDC** | **Structure** | **MW** | **Chemical Formula** | **SMILES** | **IUPAC Name** |
| --- | --- | --- | --- | --- | --- |
| **1553** | 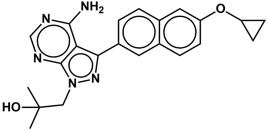 | **389.5** | **C22H23N5O2** | **NC1=C2C(N(CC(O)(C)C)N=C2C3=CC=C(C=C(OC5CC5)C=C4)C4=C3)=NC=N1** | **1-(4-amino-3-(6-cyclopropoxynaphthalen-2-yl)-1H-pyrazolo[3,4-d]pyrimidin-1-yl)-2-methylpropan-2-ol** |
| **1649** | 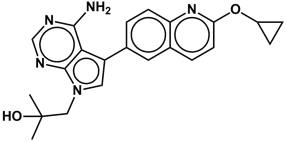 | **389.19** | **C22H23N5O2** | **OC(C)(C)CN(C=C1C2=CC=C(N=C(OC3CC3)C=C4)C4=C2)C5=C1C(N)=NC=N5** | **1-(4-amino-5-(2-cyclopropoxyquinolin-6-yl)- 7H-pyrrolo[2,3-d]pyrimidin-7-yl) -2-methylpropan-2-ol** |
| **1553-N-Me** | 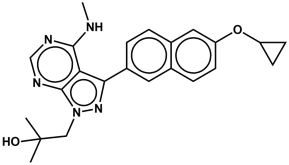 | **403.5** | **C23H25N5O2** | **NC1=C2C(N(CC(C)(OC)C)N=C2C3=CC=C(C=C(OC5CC5)C=C4)C4=C3)=NC=N1** | **1-(3-(6-cyclopropoxynaphthalen-2-yl)-4-(methylamino)-1H-pyrazolo[3,4-d]pyrimidin-1-yl)-2-methylpropan-2-ol** |
| **1811** | 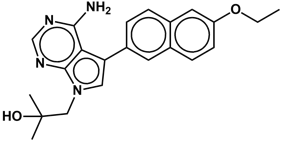 | **376.4** | **C22H24N4O2** | **NC1=C2C(N(CC(C)(C)O)C=C2C3=CC=C(C=C(OCC)C=C4)C4=C3)=NC=N1** | **1-(4-amino-5-(6-ethoxynaphthalen-2-yl)-7H-pyrrolo[2,3-d]pyrimidin-7-yl)-2-methylpropan-2-ol** |
| **1812** | 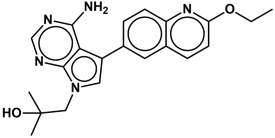 | **377.4** | **C21H23N5O2** | **NC1=C2C(N(CC(C)(C)O)C=C2C3=CC=C(N=C(OCC)C=C4)C4=C3)=NC=N1** | **1-(4-amino-5-(2-ethoxyquinolin-6-yl)-7H-pyrrolo[2,3-d]pyrimidin-7-yl)-2-methylpropan-2-ol** |
| **1813** | 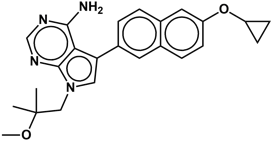 | **402.2** | **C24H26N4O2** | **NC1=C2C(N(CC(C)(C)OC)C=C2C3=CC=C(C=C(OC5CC5)C=C4)C4=C3)=NC=N1** | **5-(6-cyclopropoxynaphthalen-2-yl)-7-(2-methoxy-2-methylpropyl)-7H-pyrrolo[2,3-d]pyrimidin-4-amine** |
| **1814** | 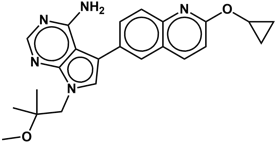 | **403.4** | **C23H25N5O2** | **NC1=C2C(N(CC(C)(C)OC)C=C2C3=CC=C(N=C(OC5CC5)C=C4)C4=C3)=NC=N1** | **5-(2-cyclopropoxyquinolin-6-yl)-7-(2-methoxy-2-methylpropyl)-7H-pyrrolo[2,3-d]pyrimidin-4-amine** |
| **1815** | 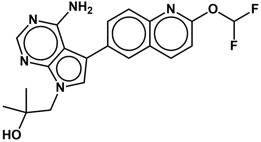 | **399.4** | **C20H19F2N5O2** | **NC1=C2C(N(CC(C)(C)O)C=C2C3=CC=C(N=C(OC(F)F)C=C4)C4=C3)=NC=N1** | **1-(4-amino-5-(2-(difluoromethoxy)quinolin-6-yl)-7H-pyrrolo[2,3-d]pyrimidin-7-yl)-2-methylpropan-2-ol** |
| **1817** | 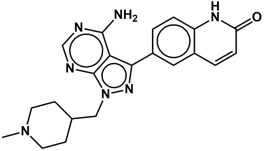 | **389.45** | **C21H23N7O** | **NC1=C2C(N(CC4CCN(C)CC4)N=C2C3=CC=C(N=C(O)C=C5)C5=C3)=NC=N1** | **6-(4-amino-1-((1-methylpiperidin-4-yl)methyl)-1H-pyrazolo[3,4-d]pyrimidin-3-yl)quinolin-2-ol** |
| **1818** | 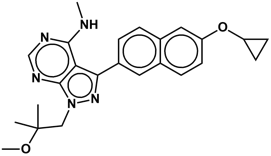 | **417.5** | **C24H27N5O2** | **COC(C)(C)CN2C1=NC=NC(NC)=C1C(C3=CC=C(C=C(OC5CC5)C=C4)C4=C3)=N2** | **3-(6-cyclopropoxynaphthalen-2-yl)-1-(2-methoxy-2-methylpropyl)-N-methyl-1H-pyrazolo[3,4-d]pyrimidin-4-amine** |
| **1819** | 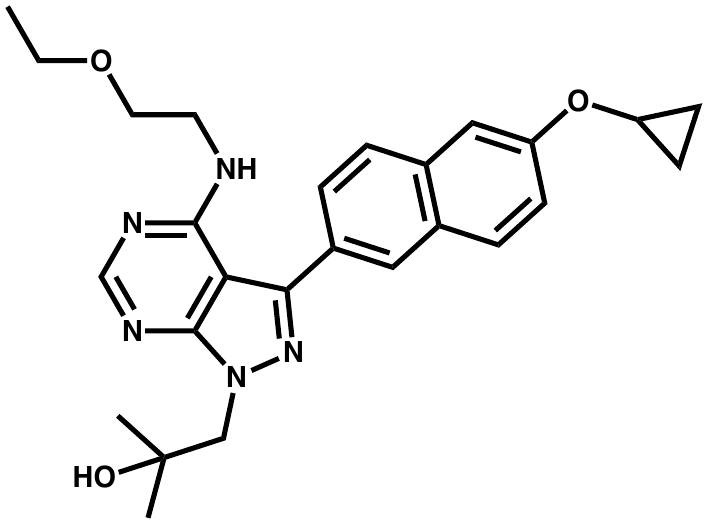 | **461.5** | **C26H31N5O3** | **NC1=C2C(N(CC(C)(C)OCCOCC)N=C2C3=CC=C(C=C(OC5CC5)C=C4)C4=C3)=NC=N1** | **3-(6-cyclopropoxynaphthalen-2-yl)-1-(2-(2-ethoxyethoxy)-2-methylpropyl)-1H-pyrazolo[3,4-d]pyrimidin-4-amine** |
| **1820** | 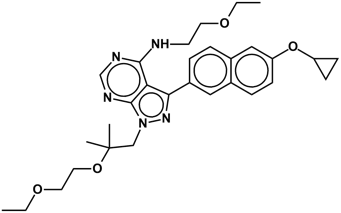 | **533.7** | **C30H39N5O4** | **CC(OCCOCC)(C)CN2C1=NC=NC(NCCOCC)=C1C(C3=CC=C(C=C(OC5CC5)C=C4)C4=C3)=N2** | **3-(6-cyclopropoxynaphthalen-2-yl)-1-(2-(2-ethoxyethoxy)-2-methylpropyl)-N-(2-ethoxyethyl)-1H-pyrazolo[3,4-d]pyrimidin-4-amine** |
| **1826** | 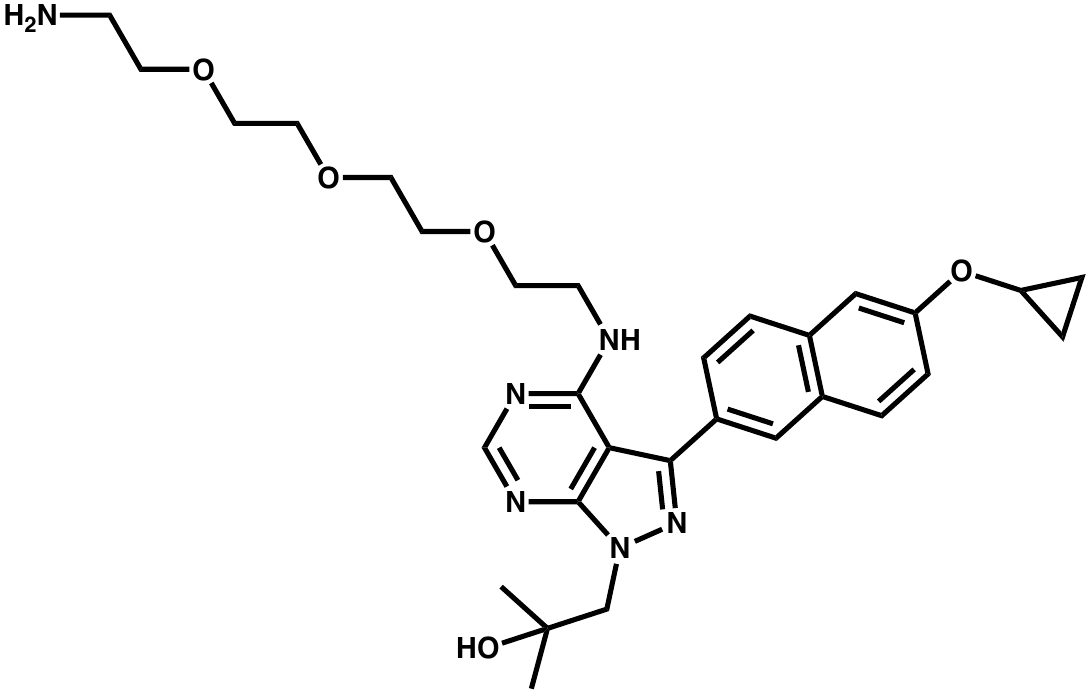 | **564.6** | **C30H40N6O5** | **NC1=C2C(N(CC(OCCOCCOCCOCCN)(C)C)N=C2C3=CC=C(C=C(OC5CC5)C=C4)C4=C3)=NC=N1** | **1-(2-(2-(2-(2-(2-aminoethoxy)ethoxy)ethoxy)ethoxy)-2-methylpropyl)-3-(6-cyclopropoxynaphthalen-2-yl)-1H-pyrazolo[3,4-d]pyrimidin-4-amine** |
| **GP223** | 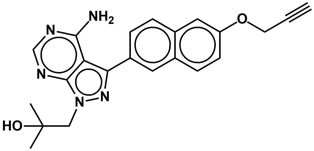 | **387.43** | **C22H21N5O2** | **NC1=C2C(N(N=C2C3=CC=C4C(C=CC(OCC#C)=C4)=C3)CC(O)(C)C)=NC=N1** | **1-(4-amino-3-(6-(prop-2-yn-1-yloxy)naphthalen-2-yl)-1H-pyrazolo[3,4-d]pyrimidin-1-yl)-2-methylpropan-2-ol** |
| **GP224** | 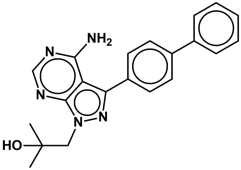 | **359.42** | **C21H21N5O** | **NC1=C2C(N(N=C2C3=CC=C(C4=CC=CC=C4)C=C3)CC(O)(C)C)=NC=N1** | **1-(3-([1,1'-biphenyl]-4-yl)-4-amino-1H-pyrazolo[3,4-d]pyrimidin-1-yl)-2-methylpropan-2-ol** |
| **GP225** | 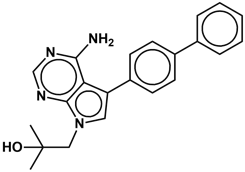 | **358.44** | **C22H22N4O** | **NC1=C2C(N(C=C2C3=CC=C(C4=CC=CC=C4)C=C3)CC(O)(C)C)=NC=N1** | **1-(5-([1,1'-biphenyl]-4-yl)-4-amino-7H-pyrrolo[2,3-d]pyrimidin-7-yl)-2-methylpropan-2-ol** |
| **GP226** | 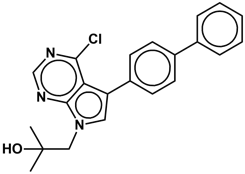 | **377.87** | **C22H20ClN3O** | **ClC1=C2C(N(C=C2C3=CC=C(C4=CC=CC=C4)C=C3)CC(O)(C)C)=NC=N1** | **1-(5-([1,1'-biphenyl]-4-yl)-4-chloro-7H-pyrrolo[2,3-d]pyrimidin-7-yl)-2-methylpropan-2-ol** |
| **GP227** | 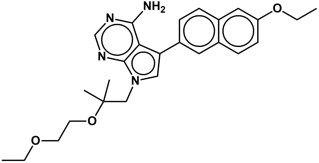 | **448.56** | **C26H32N4O3** | **CCOC1=CC2=CC=C(C3=CN(CC(OCCOCC)(C)C)C4=NC=NC(N)=C43)C=C2C=C1** | **7-(2-(2-ethoxyethoxy)-2-methylpropyl)-5-(6-ethoxynaphthalen-2-yl)-7H-pyrrolo[2,3-d]pyrimidin-4-amine** |
| **GP228** | 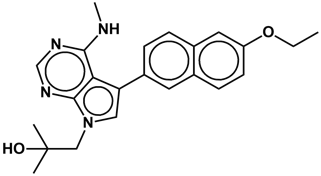 | **390.48** | **C23H26N4O2** | **CCOC1=CC2=CC=C(C3=CN(CC(O)(C)C)C4=NC=NC(NC)=C43)C=C2C=C1** | **1-(5-(6-ethoxynaphthalen-2-yl)-4-(methylamino)-7H-pyrrolo[2,3-d]pyrimidin-7-yl)-2-methylpropan-2-ol** |
| **GP229** | 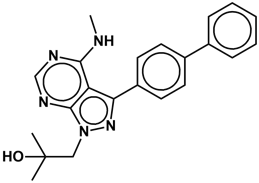 | **373.45** | **C22H23N5O** | **CC(O)(C)CN1C2=NC=NC(NC)=C2C(C3=CC=C(C4=CC=CC=C4)C=C3)=N1** | **1-(3-([1,1'-biphenyl]-4-yl)-4-(methylamino)-1H-pyrazolo[3,4-d]pyrimidin-1-yl)-2-methylpropan-2-ol** |
| **GP230** | 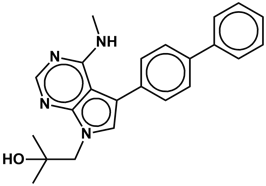 | **372.46** | **C23H24N4O** | **CC(O)(C)CN1C2=NC=NC(NC)=C2C(C3=CC=C(C4=CC=CC=C4)C=C3)=C1** | **1-(5-([1,1'-biphenyl]-4-yl)-4-(methylamino)-7H-pyrrolo[2,3-d]pyrimidin-7-yl)-2-methylpropan-2-ol** |
| **GP231** | 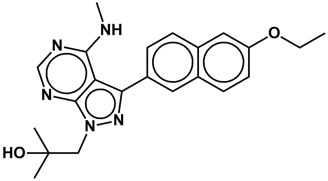 | **391.47** | **C22H25N5O2** | **CCOC1=CC2=CC=C(C3=NN(CC(O)(C)C)C4=NC=NC(NC)=C43)C=C2C=C1** | **1-(3-(6-ethoxynaphthalen-2-yl)-4-(methylamino)-1H-pyrazolo[3,4-d]pyrimidin-1-yl)-2-methylpropan-2-ol** |
| **GP232** | 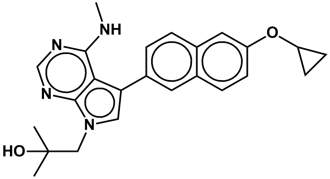 | **402.49** | **C24H26N4O2** | **CC(O)(C)CN1C2=NC=NC(NC)=C2C(C3=CC=C4C(C=CC(OC5CC5)=C4)=C3)=C1** | **1-(5-(6-cyclopropoxynaphthalen-2-yl)-4-(methylamino)-7H-pyrrolo[2,3-d]pyrimidin-7-yl)-2-methylpropan-2-ol** |
| **GP233** | 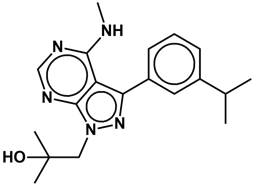 | **339.43** | **C19H25N5O** | **CC(O)(C)CN1C2=NC=NC(NC)=C2C(C3=CC=CC(C(C)C)=C3)=N1** | **1-(3-(3-isopropylphenyl)-4-(methylamino)-1H-pyrazolo[3,4-d]pyrimidin-1-yl)-2-methylpropan-2-ol** |
| **GP234** | 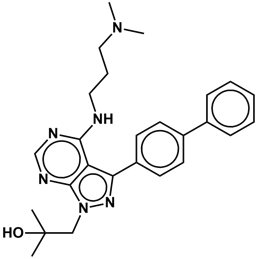 | **444.57** | **C26H32N6O** | **CC(O)(C)CN1C2=NC=NC(NCCCN(C)C)=C2C(C3=CC=C(C4=CC=CC=C4)C=C3)=N1** | **1-(3-([1,1'-biphenyl]-4-yl)-4-((3-(dimethylamino)propyl)amino)-1H-pyrazolo[3,4-d]pyrimidin-1-yl)-2-methylpropan-2-ol** |
| **GP235** | 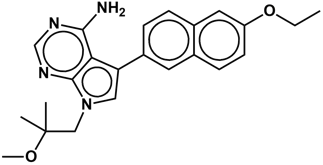 | **390.48** | **C23H26N4O2** | **CCOC1=CC2=CC=C(C3=CN(CC(OC)(C)C)C4=NC=NC(N)=C43)C=C2C=C1** | **5-(6-ethoxynaphthalen-2-yl)-7-(2-methoxy-2-methylpropyl)-7H-pyrrolo[2,3-d]pyrimidin-4-amine** |
| **GP236** | 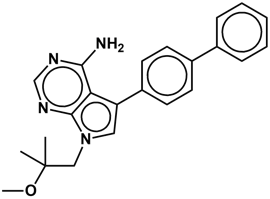 | **372.46** | **C23H24N4O** | **NC1=C2C(N(C=C2C3=CC=C(C4=CC=CC=C4)C=C3)CC(OC)(C)C)=NC=N1** | **5-([1,1'-biphenyl]-4-yl)-7-(2-methoxy-2-methylpropyl)-7H-pyrrolo[2,3-d]pyrimidin-4-amine** |
| **GP237** | 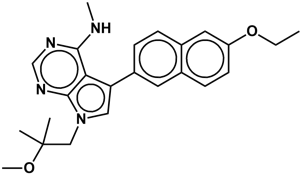 | **404.5** | **C24H28N4O2** | **CCOC1=CC2=CC=C(C3=CN(CC(OC)(C)C)C4=NC=NC(NC)=C43)C=C2C=C1** | **5-(6-ethoxynaphthalen-2-yl)-7-(2-methoxy-2-methylpropyl)-N-methyl-7H-pyrrolo[2,3-d]pyrimidin-4-amine** |
| **GP238** | 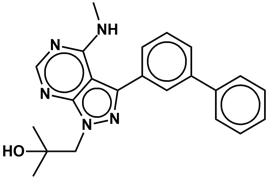 | **373.45** | **C22H23N5O** | **CC(O)(C)CN1C2=NC=NC(NC)=C2C(C3=CC=CC(C4=CC=CC=C4)=C3)=N1** | **1-(3-([1,1'-biphenyl]-3-yl)-4-(methylamino)-1H-pyrazolo[3,4-d]pyrimidin-1-yl)-2-methylpropan-2-ol** |
| **GP239** | 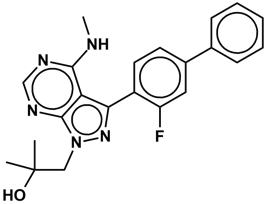 | **391.44** | **C22H22FN5O** | **CC(O)(C)CN1C2=NC=NC(NC)=C2C(C3=CC=C(C4=CC=CC=C4)C=C3F)=N1** | **1-(3-(3-fluoro-[1,1'-biphenyl]-4-yl)-4-(methylamino)-1H-pyrazolo[3,4-d]pyrimidin-1-yl)-2-methylpropan-2-ol** |
| **GP240** | 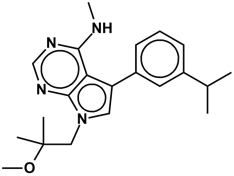 | **352.47** | **C21H28N4O** | **CNC1=C2C(N(C=C2C3=CC=CC(C(C)C)=C3)CC(C)(C)OC)=NC=N1** | **5-(3-isopropylphenyl)-7-(2-methoxy-2-methylpropyl)-N-methyl-7H-pyrrolo[2,3-d]pyrimidin-4-amine** |
| **GP241** | 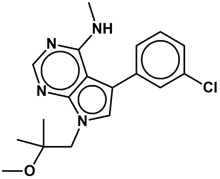 | **344.84** | **C18H21ClN4O** | **CNC1=C2C(N(C=C2C3=CC=CC(Cl)=C3)CC(C)(C)OC)=NC=N1** | **5-(3-chlorophenyl)-7-(2-methoxy-2-methylpropyl)-N-methyl-7H-pyrrolo[2,3-d]pyrimidin-4-amine** |
| **GP242** | 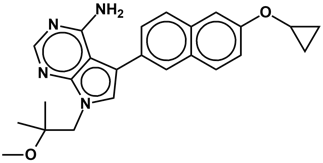 | **402.49** | **C24H26N4O2** | **NC1=C2C(N(C=C2C3=CC=C4C(C=CC(OC5CC5)=C4)=C3)CC(OC)(C)C)=NC=N1** | **5-(6-cyclopropoxynaphthalen-2-yl)-7-(2-methoxy-2-methylpropyl)-7H-pyrrolo[2,3-d]pyrimidin-4-amine** |
| **GP243** | 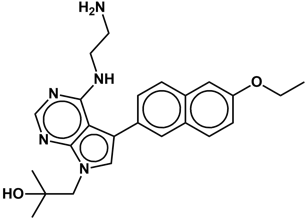 | **419.52** | **C24H29N5O2** | **NCCNC1=C2C(N(CC(C)(C)O)C=C2C3=CC(C=CC(OCC)=C4)=C4C=C3)=NC=N1** | **1-(4-((2-aminoethyl)amino)-5-(6-ethoxynaphthalen-2-yl)-7H-pyrrolo[2,3-d]pyrimidin-7-yl)-2-methylpropan-2-ol** |
| **GP244** | 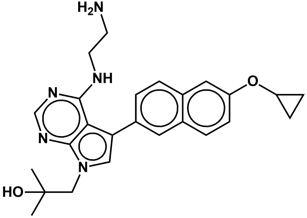 | **431.53** | **C25H29N5O2** | **NCCNC1=C2C(N(CC(C)(C)O)C=C2C3=CC(C=CC(OC4CC4)=C5)=C5C=C3)=NC=N1** | **1-(4-((2-aminoethyl)amino)-5-(6-cyclopropoxynaphthalen-2-yl)-7H-pyrrolo[2,3-d]pyrimidin-7-yl)-2-methylpropan-2-ol** |
| **GP245** | 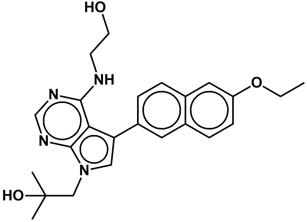 | **420.5** | **C24H28N4O3** | **OCCNC1=C2C(N(CC(C)(C)O)C=C2C3=CC(C=CC(OCC)=C4)=C4C=C3)=NC=N1** | **1-(5-(6-ethoxynaphthalen-2-yl)-4-((2-hydroxyethyl)amino)-7H-pyrrolo[2,3-d]pyrimidin-7-yl)-2-methylpropan-2-ol** |
| **GP246** | 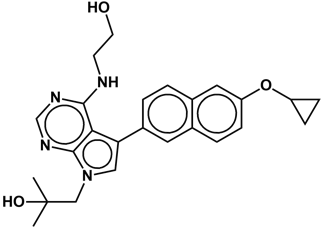 | **432.51** | **C25H28N4O3** | **OCCNC1=C2C(N(CC(C)(C)O)C=C2C3=CC(C=CC(OC4CC4)=C5)=C5C=C3)=NC=N1** | **1-(5-(6-cyclopropoxynaphthalen-2-yl)-4-((2-hydroxyethyl)amino)-7H-pyrrolo[2,3-d]pyrimidin-7-yl)-2-methylpropan-2-ol** |
| **GP247** | 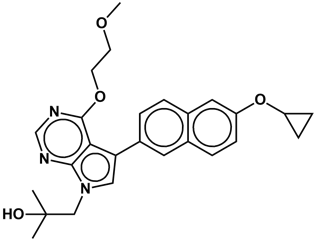 | **447.53** | **C26H29N3O4** | **OC(C)(C)CN1C2=NC=NC(OCCOC)=C2C(C3=CC(C=CC(OC4CC4)=C5)=C5C=C3)=C1** | **1-(5-(6-cyclopropoxynaphthalen-2-yl)-4-(2-methoxyethoxy)-7H-pyrrolo[2,3-d]pyrimidin-7-yl)-2-methylpropan-2-ol** |
| **GP248** | 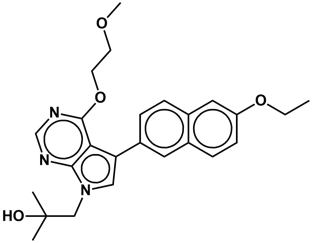 | **435.52** | **C25H29N3O4** | **OC(C)(C)CN1C2=NC=NC(OCCOC)=C2C(C3=CC(C=CC(OCC)=C4)=C4C=C3)=C1** | **1-(5-(6-ethoxynaphthalen-2-yl)-4-(2-methoxyethoxy)-7H-pyrrolo[2,3-d]pyrimidin-7-yl)-2-methylpropan-2-ol** |
| **GP249** | 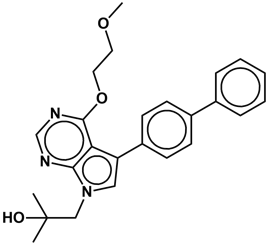 | **417.5** | **C25H27N3O3** | **OC(C)(C)CN1C2=NC=NC(OCCOC)=C2C(C3=CC=C(C4=CC=CC=C4)C=C3)=C1** | **1-(5-([1,1'-biphenyl]-4-yl)-4-(2-methoxyethoxy)-7H-pyrrolo[2,3-d]pyrimidin-7-yl)-2-methylpropan-2-ol** |
| **GP250** | 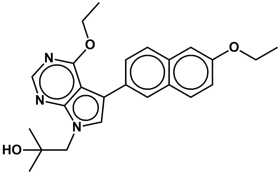 | **405.49** | **C24H27N3O3** | **OC(C)(C)CN1C2=NC=NC(OCC)=C2C(C3=CC(C=CC(OCC)=C4)=C4C=C3)=C1** | **1-(4-ethoxy-5-(6-ethoxynaphthalen-2-yl)-7H-pyrrolo[2,3-d]pyrimidin-7-yl)-2-methylpropan-2-ol** |
| **GP251** | 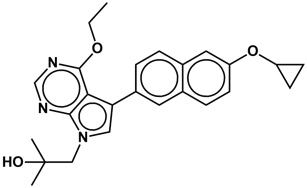 | **417.5** | **C25H27N3O3** | **CCOC1=C2C(N(CC(C)(C)O)C=C2C3=CC(C=CC(OC4CC4)=C5)=C5C=C3)=NC=N1** | **1-(5-(6-cyclopropoxynaphthalen-2-yl)-4-ethoxy-7H-pyrrolo[2,3-d]pyrimidin-7-yl)-2-methylpropan-2-ol** |
| **GP252** | 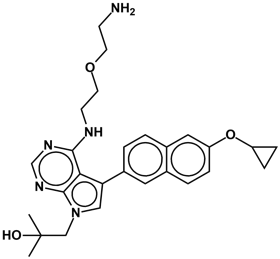 | **475.58** | **C27H33N5O3** | **CC(C)(O)CN1C2=NC=NC(NCCOCCN)=C2C(C3=CC(C=CC(OC4CC4)=C5)=C5C=C3)=C1** | **1-(4-((2-(2-aminoethoxy)ethyl)amino)-5-(6-cyclopropoxynaphthalen-2-yl)-7H-pyrrolo[2,3-d]pyrimidin-7-yl)-2-methylpropan-2-ol** |
| **GP253** | 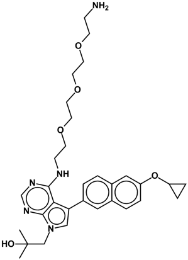 | **563.69** | **C31H41N5O5** | **NCCOCCOCCOCCNC1=C2C(N(CC(C)(O)C)C=C2C3=CC(C=CC(OC4CC4)=C5)=C5C=C3)=NC=N1** | **1-(4-((2-(2-(2-(2-aminoethoxy)ethoxy)ethoxy)ethyl)amino)-5-(6-cyclopropoxynaphthalen-2-yl)-7H-pyrrolo[2,3-d]pyrimidin-7-yl)-2-methylpropan-2-ol** |
| **GP254** | 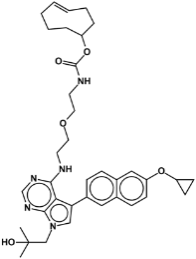 | **627.77** | **C36H45N5O5** | **CC(C)(O)CN1C2=NC=NC(NCCOCCNC(OC3CC/C=C/CCC3)=O)=C2C(C4=CC(C=CC(OC5CC5)=C6)=C6C=C4)=C1** | **(E)-cyclooct-4-en-1-yl (2-(2-((5-(6-cyclopropoxynaphthalen-2-yl)-7-(2-hydroxy-2-methylpropyl)-7H-pyrrolo[2,3-d]pyrimidin-4-yl)amino)ethoxy)ethyl)carbamate** |
| **GP255** | 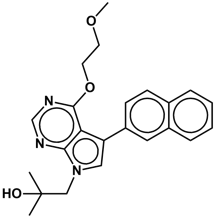 | **391.46** | **C23H25N3O3** | **OC(C)(C)CN1C2=NC=NC(OCCOC)=C2C(C3=CC(C=CC=C4)=C4C=C3)=C1** | **1-(4-(2-methoxyethoxy)-5-(naphthalen-2-yl)-7H-pyrrolo[2,3-d]pyrimidin-7-yl)-2-methylpropan-2-ol** |
| **GP256** | 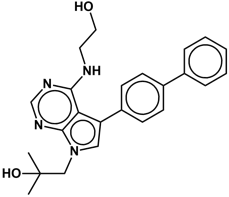 | **402.49** | **C24H26N4O2** | **OC(C)(C)CN1C2=NC=NC(NCCO)=C2C(C3=CC=C(C4=CC=CC=C4)C=C3)=C1** | **1-(5-([1,1'-biphenyl]-4-yl)-4-((2-hydroxyethyl)amino)-7H-pyrrolo[2,3-d]pyrimidin-7-yl)-2-methylpropan-2-ol** |
| **GP260** | 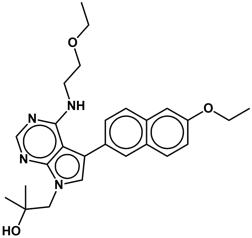 | **448.56** | **C26H32N4O3** | **OC(C)(C)CN1C2=NC=NC(NCCOCC)=C2C(C3=CC(C=CC(OCC)=C4)=C4C=C3)=C1** | **1-(4-((2-ethoxyethyl)amino)-5-(6-ethoxynaphthalen-2-yl)-7H-pyrrolo[2,3-d]pyrimidin-7-yl)-2-methylpropan-2-ol** |
| **GP261** | 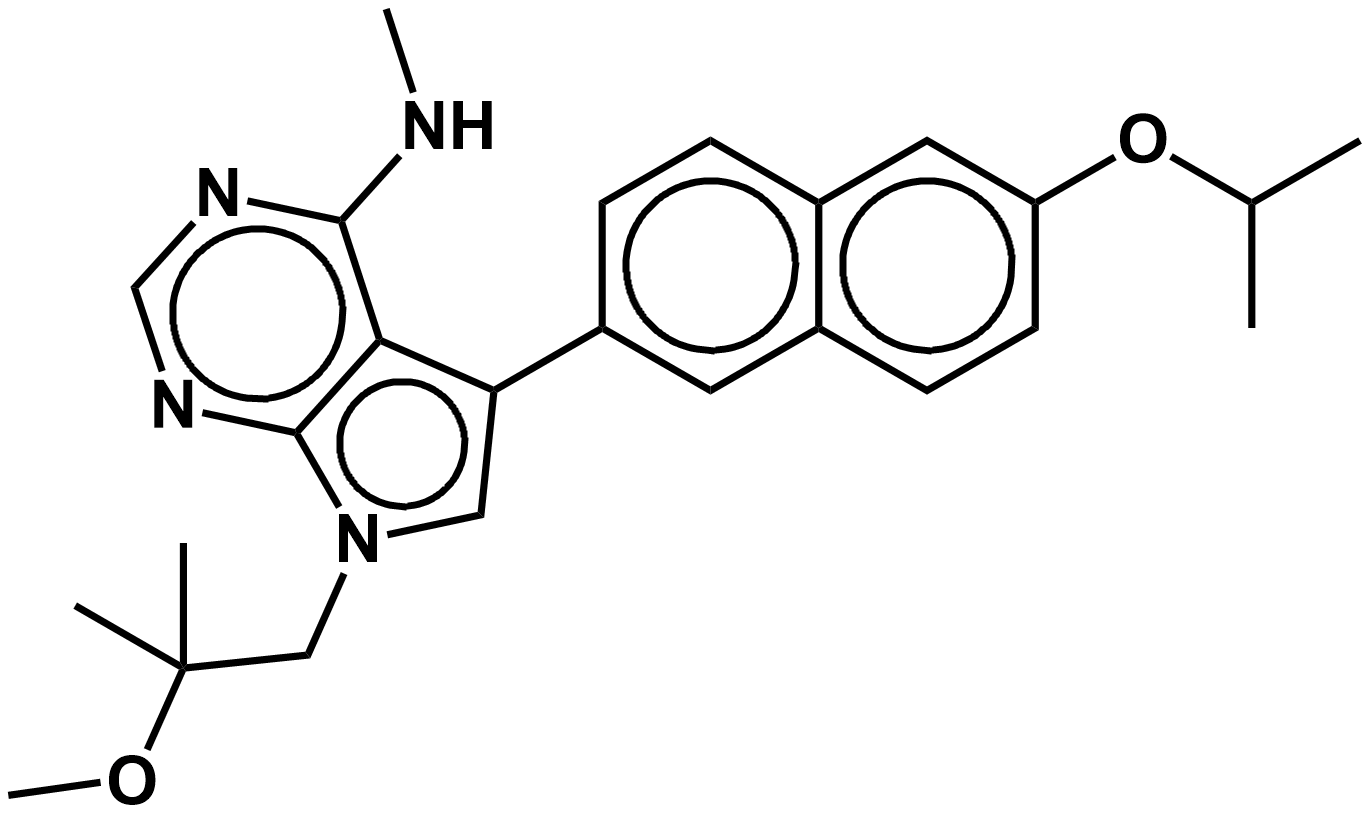 | **418.24** | **C25H30N4O2** | **CNC1=C2C(N(CC(C)(C)OC)C=C2C3=CC(C=CC(OC(C)C)=C4)=C4C=C3)=NC=N1** | **5-(6-isopropoxynaphthalen-2-yl)-7-(2-methoxy-2-methylpropyl)-N-methyl-7H-pyrrolo[2,3-d]pyrimidin-4-amine** |
| **GP262** | 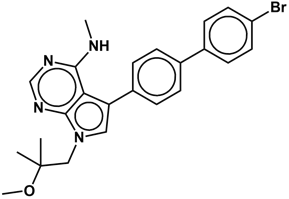 | **464.12** | **C24H25BrN4O** | **CNC1=C2C(N(CC(C)(C)OC)C=C2C3=CC=C(C4=CC=C(Br)C=C4)C=C3)=NC=N1** | **5-(4'-bromo-[1,1'-biphenyl]-4-yl)-7-(2-methoxy-2-methylpropyl)-N-methyl-7H-pyrrolo[2,3-d]pyrimidin-4-amine** |
| **GP263** | 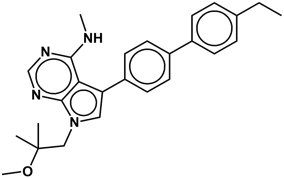 | **414.24** | **C26H30N4O** | **CNC1=C2C(N(CC(C)(C)OC)C=C2C3=CC=C(C4=CC=C(CC)C=C4)C=C3)=NC=N1** | **5-(4'-ethyl-[1,1'-biphenyl]-4-yl)-7-(2-methoxy-2-methylpropyl)-N-methyl-7H-pyrrolo[2,3-d]pyrimidin-4-amine** |
| **GP264** | 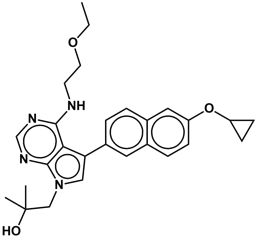 | **460.25** | **C27H32N4O3** | **OC(C)(C)CN1C2=NC=NC(NCCOCC)=C2C(C3=CC(C=CC(OC4CC4)=C5)=C5C=C3)=C1** | **1-(5-(6-cyclopropoxynaphthalen-2-yl)-4-((2-ethoxyethyl)amino)-7H-pyrrolo[2,3-d]pyrimidin-7-yl)-2-methylpropan-2-ol** |
| **GP265** | 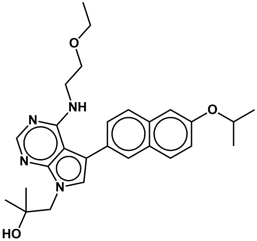 | **462.26** | **C27H34N4O3** | **OC(C)(C)CN1C2=NC=NC(NCCOCC)=C2C(C3=CC(C=CC(OC(C)C)=C4)=C4C=C3)=C1** | **1-(4-((2-ethoxyethyl)amino)-5-(6-isopropoxynaphthalen-2-yl)-7H-pyrrolo[2,3-d]pyrimidin-7-yl)-2-methylpropan-2-ol** |
| **GP266** | 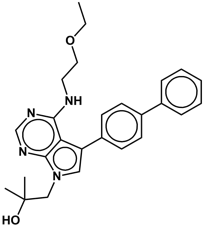 | **430.24** | **C26H30N4O2** | **OC(C)(C)CN1C2=NC=NC(NCCOCC)=C2C(C3=CC=C(C4=CC=CC=C4)C=C3)=C1** | **1-(5-([1,1'-biphenyl]-4-yl)-4-((2-ethoxyethyl)amino)-7H-pyrrolo[2,3-d]pyrimidin-7-yl)-2-methylpropan-2-ol** |
| **GP267** | 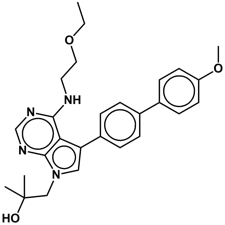 | **460.25** | **C27H32N4O3** | **OC(C)(C)CN1C2=NC=NC(NCCOCC)=C2C(C3=CC=C(C4=CC=C(OC)C=C4)C=C3)=C1** | **1-(4-((2-ethoxyethyl)amino)-5-(4'-methoxy-[1,1'-biphenyl]-4-yl)-7H-pyrrolo[2,3-d]pyrimidin-7-yl)-2-methylpropan-2-ol** |
| **GP268** | 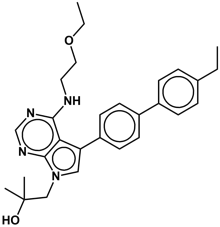 | **458.27** | **C28H34N4O2** | **OC(C)(C)CN1C2=NC=NC(NCCOCC)=C2C(C3=CC=C(C4=CC=C(CC)C=C4)C=C3)=C1** | **1-(4-((2-ethoxyethyl)amino)-5-(4'-ethyl-[1,1'-biphenyl]-4-yl)-7H-pyrrolo[2,3-d]pyrimidin-7-yl)-2-methylpropan-2-ol** |
| **GP269** | 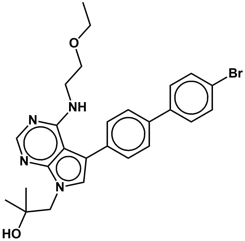 | **508.15** | **C26H29BrN4O2** | **OC(C)(C)CN1C2=NC=NC(NCCOCC)=C2C(C3=CC=C(C4=CC=C(Br)C=C4)C=C3)=C1** | **1-(5-(4'-bromo-[1,1'-biphenyl]-4-yl)-4-((2-ethoxyethyl)amino)-7H-pyrrolo[2,3-d]pyrimidin-7-yl)-2-methylpropan-2-ol** |
| **GP270** | 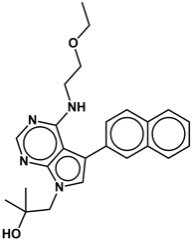 | **404.22** | **C24H28N4O2** | **OC(C)(C)CN1C2=NC=NC(NCCOCC)=C2C(C3=CC(C=CC=C4)=C4C=C3)=C1** | **1-(4-((2-ethoxyethyl)amino)-5-(naphthalen-2-yl)-7H-pyrrolo[2,3-d]pyrimidin-7-yl)-2-methylpropan-2-ol** |
| **GP271** | 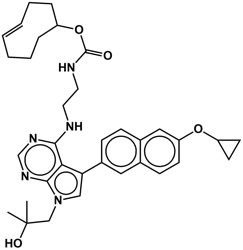 | **583.32** | **C34H41N5O4** | **CC(C)(O)CN1C2=NC=NC(NCCNC(OC3CC/C=C/CCC3)=O)=C2C(C4=CC(C=CC(OC5CC5)=C6)=C6C=C4)=C1** | **(E)-cyclooct-4-en-1-yl (2-((5-(6-cyclopropoxynaphthalen-2-yl)-7-(2-hydroxy-2-methylpropyl)-7H-pyrrolo[2,3-d]pyrimidin-4-yl)amino)ethyl)carbamate** |

**Table S2. Screening results of BKIDCs tested at 10 µM in LNCaP95 and PC3 cells in the MTS assay.**

|  | **LNCaP95** | | **PC3** | |
| --- | --- | --- | --- | --- |
|  | **mean** | **stdev** | **mean** | **stdev** |
| BKIDC-1318 | 0.7017 | 0.0166 | 1.0071 | 0.0306 |
| BKIDC-1369 | 0.6517 | 0.0209 | 0.9909 | 0.0229 |
| BKIDC-1417 | 0.9702 | 0.0153 | 1.0542 | 0.0318 |
| BKIDC-1527 | 0.9991 | 0.0698 | 1.1010 | 0.0295 |
| BKIDC-1537 | 0.6374 | 0.0345 | 1.0836 | 0.0389 |
| BKIDC-1539 | 0.6102 | 0.0054 | 1.0762 | 0.0124 |
| BKIDC-1541 | 0.8185 | 0.0281 | 1.0857 | 0.0133 |
| BKIDC-1553 | 0.5763 | 0.0488 | 0.9214 | 0.0589 |
| BKIDC-1556 | 0.9214 | 0.0000 | 0.9455 | 0.0176 |
| BKIDC-1561 | 1.3836 | 0.0999 | 1.0951 | 0.0180 |
| BKIDC-1614 | 1.1843 | 0.0322 | 1.0759 | 0.0299 |
| BKIDC-1624 | 1.2232 | 0.0770 | 1.1189 | 0.0801 |
| BKIDC-1649 | 0.5717 | 0.0172 | 0.8779 | 0.0198 |
| BKIDC-1651 | 0.4429 | 0.0154 | 0.8064 | 0.0160 |
| BKIDC-1659 | 0.6872 | 0.0354 | 1.0162 | 0.0413 |
| BKIDC-1660 | 0.6215 | 0.0317 | 1.0259 | 0.0161 |
| BKIDC-1662 | 0.5689 | 0.0195 | 1.0001 | 0.0146 |
| BKIDC-1663 | 0.6014 | 0.0133 | 0.9827 | 0.0281 |
| BKIDC-1664 | 0.7761 | 0.0158 | 0.9743 | 0.0065 |
| BKIDC-1666 | 0.8463 | 0.0218 | 1.1291 | 0.0492 |
| BKIDC-1672 | 0.7695 | 0.0147 | 1.0380 | 0.0181 |
| BKIDC-1553-N-Me | 0.4714 | 0.0336 | 0.9462 | 0.0503 |
| BKIDC-1677 | 0.7270 | 0.0189 | 0.9404 | 0.0223 |
| BKIDC-1693 | 0.6890 | 0.0232 | 0.9357 | 0.0472 |
| BKIDC-1700 | 1.0443 | 0.0614 | 1.0612 | 0.0104 |
| BKIDC-1701 | 1.0417 | 0.0417 | 1.0731 | 0.0404 |
| BKIDC-1703 | 0.9301 | 0.0794 | 1.0420 | 0.0116 |
| BKIDC-1729 | 1.1103 | 0.0484 | 1.0471 | 0.0526 |
| BKIDC-1730 | 0.8164 | 0.0871 | 1.0251 | 0.0498 |
| BKIDC-1739 | 1.0541 | 0.0392 | 1.0708 | 0.0492 |
| BKIDC-1741 | 0.9334 | 0.0408 | 1.0007 | 0.0573 |
| BKIDC-1753 | 0.6208 | 0.0020 | 1.0255 | 0.0312 |
| BKIDC-1759 | 0.8139 | 0.0262 | 0.8465 | 0.0300 |
| BKIDC-1785 | 0.9153 | 0.0562 | 0.9451 | 0.0217 |
| BKIDC-1789 | 1.0935 | 0.0057 | 0.9962 | 0.0138 |
| BKIDC-1811 | 0.6360 | 0.0319 | 0.9505 | 0.0360 |
| BKIDC-1812 | 0.4133 | 0.0139 | 0.8785 | 0.0146 |
| BKIDC-1813 | 0.5541 | 0.0117 | 0.9168 | 0.0211 |
| BKIDC-1814 | 0.5803 | 0.0528 | 0.9441 | 0.0355 |
| BKIDC-1815 | 0.6679 | 0.0628 | 0.9089 | 0.0427 |
| BKIDC-1817 | 1.0215 | 0.1100 | 0.9930 | 0.0760 |
| BKIDC-1819 | 0.5621 | 0.0686 | 1.0931 | 0.1159 |
| BKIDC-1820 | 0.5821 | 0.0733 | 1.1993 | 0.0557 |
| BKIDC-1826 | 0.4482 | 0.0293 | 0.6803 | 0.0463 |
| BKIDC-1829 | 0.6777 | 0.1281 | 1.0129 | 0.0736 |
| GP 223 | 0.5517 | 0.0719 | 0.8252 | 0.0304 |
| GP 224 | 0.6450 | 0.0752 | 0.7792 | 0.0882 |
| GP 225 | 0.5151 | 0.0555 | 0.7530 | 0.0707 |
| GP 226 | 0.8240 | 0.0756 | 0.9503 | 0.0647 |
| GP 227 | 0.4819 | 0.0860 | 0.7938 | 0.0959 |
| GP 228 | 0.4331 | 0.0480 | 0.9701 | 0.0871 |
| GP 229 | 0.6096 | 0.0655 | 0.9372 | 0.0944 |
| GP 230 | 0.5687 | 0.0716 | 1.0666 | 0.0402 |
| GP 231 | 0.5514 | 0.0644 | 1.0591 | 0.0310 |
| GP 232 | 0.4501 | 0.0404 | 0.9945 | 0.0401 |
| GP 233 | 0.8540 | 0.0330 | 1.1305 | 0.0878 |
| GP 234 | 0.5986 | 0.0707 | 1.1131 | 0.0748 |
| GP 235 | 0.5855 | 0.0804 | 1.0199 | 0.0961 |
| GP 236 | 0.6361 | 0.0594 | 0.9555 | 0.0812 |
| GP 237 | 0.5779 | 0.0338 | 0.7822 | 0.1012 |
| GP 238 | 0.8250 | 0.1435 | 0.9464 | 0.0299 |
| GP 238 | 0.7293 | 0.0415 | 0.8181 | 0.0404 |
| GP 239 | 0.8368 | 0.0855 | 0.8418 | 0.0613 |
| GP 240 | 0.7420 | 0.0689 | 1.0140 | 0.0898 |
| GP 241 | 0.7129 | 0.0874 | 0.9146 | 0.1144 |
| GP 242 | 0.5668 | 0.0555 | 1.0298 | 0.0527 |
| GP 243 | 0.8027 | 0.1003 | 0.8066 | 0.0855 |
| GP 244 | 0.7961 | 0.0655 | 0.8770 | 0.0525 |
| GP 245 | 0.6115 | 0.0582 | 1.0129 | 0.0443 |
| GP 246 | 0.5819 | 0.0344 | 1.0389 | 0.0826 |
| GP 247 | 0.5868 | 0.0432 | 0.8028 | 0.0672 |
| GP 248 | 0.6032 | 0.0395 | 0.8551 | 0.0817 |
| GP 249 | 0.7102 | 0.0701 | 1.0119 | 0.0646 |
| GP 250 | 0.5932 | 0.0295 | 1.0002 | 0.0790 |
| GP 251 | 0.7375 | 0.0905 | 0.9011 | 0.0746 |
| GP 252 | 0.8403 | 0.1079 | 0.9855 | 0.0576 |
| GP 253 | 0.5610 | 0.0509 | 1.1028 | 0.0800 |
| GP 254 | 0.5950 | 0.0668 | 1.0465 | 0.0569 |
| GP 255 | 0.7638 | 0.0526 | 1.0593 | 0.0639 |
| GP 256 | 0.6619 | 0.1127 | 1.0379 | 0.0573 |
| GP 257 | 0.7669 | 0.0925 | 1.0659 | 0.0727 |
| GP 260 | 0.6242 | 0.0555 | 0.8963 | 0.0653 |
| GP 261 | 0.8232 | 0.1018 | 1.0296 | 0.0523 |
| GP 262 | 0.7131 | 0.0640 | 1.0453 | 0.0676 |
| GP 263 | 0.6409 | 0.0472 | 1.0865 | 0.0685 |
| GP 264 | 0.6297 | 0.0522 | 1.0474 | 0.1061 |
| GP 265 | 0.7207 | 0.0886 | 0.9355 | 0.0530 |
| GP 266 | 0.8500 | 0.1056 | 1.0388 | 0.0970 |
| GP 267 | 0.5870 | 0.0360 | 0.9788 | 0.1086 |
| GP 268 | 0.7682 | 0.0558 | 1.1321 | 0.0920 |
| GP 269 | 1.1628 | 0.0953 | 1.1198 | 0.0697 |
| GP 270 | 0.8467 | 0.0343 | 1.1317 | 0.0887 |
| GP 271 | 0.5867 | 0.0635 | 1.3589 | 0.0887 |
| GP 274 | 0.3652 | 0.0340 | 0.8101 | 0.0334 |
| GP 275 | 0.4181 | 0.0260 | 0.6197 | 0.0434 |
| GP 276 | 0.4616 | 0.0090 | 0.7482 | 0.0286 |
| GP 277 | 0.7937 | 0.0499 | 0.8094 | 0.0600 |
| GP 278 | 0.6542 | 0.0377 | 0.8123 | 0.0271 |
| GP 279 | 0.5766 | 0.0451 | 0.9134 | 0.0884 |
| GP 280 | 0.5749 | 0.0337 | 0.9222 | 0.0116 |

**Table S3. Antibodies and corresponding RPPA signal intensities relative to β-actin signals.** Cellular lysates of LNCaP95 treated with the indicated BKIDC at 20 µM for 24 h were subjected to RPPA with 70 distinct antibodies.

| **Antibody** | | **Samples/Relative RPPA signals** | | | |
| --- | --- | --- | --- | --- | --- |
| **Cell Signaling Technology Cat #** | **Name** | **DMSO (24 hr)** | **20 µM 1553 (24 hr)** | **20 µM 1676 (24 hr)** | **20 µM 1817 (24 hr)** |
| 2211 | Phospho-S6 Ribosomal Protein (S235/236) | 0.9806 | 0.1837 | 0.4148 | 0.9576 |
| 2215 | Phospho-S6 Ribosomal Protein (S240/244) | 1.0138 | 0.1963 | 0.6316 | 1.0037 |
| 2231 | Phospho-EGFR(Y845) | 1.0616 | 0.6236 | 1.0015 | 1.0634 |
| 2338 | Phospho-MEK1/2 (S221) (166F8) | 1.1965 | 0.3559 | 1.2144 | 1.3313 |
| 2831 | Phospho-cPLA2 (S505) | 1.0998 | 0.6257 | 0.8486 | 0.9420 |
| 3582 | LDHA (C4B5) | 1.1248 | 0.7707 | 0.9491 | 0.9939 |
| 4058 | Phospho-Akt (S473) (193H12) | 1.4416 | 0.7823 | 2.4281 | 1.5434 |
| 5558 | Phospho-GSK-3β (S9) (D85E12) XP® | 1.4118 | 0.1939 | 2.3468 | 1.6107 |
| 9111 | Phospho-cdc2 (Y15) | 1.0466 | 0.6704 | 1.0615 | 1.5685 |
| 9145 | Phospho-Stat3 (Y705) | 1.1702 | 0.6944 | 0.9793 | 1.0407 |
| 9167 | Phospho-Stat1 (Y701) | 1.1072 | 0.5201 | 1.1710 | 1.1021 |
| 9198 | Phospho-CREB (S133) | 1.2263 | 0.5814 | 1.1148 | 1.0095 |
| 9582 | β-Catenin (6B3) | 1.0109 | 0.8860 | 1.1698 | 1.0290 |
| 9741 | Phospho-MARCKS (S152/156) | 1.1126 | 0.7664 | 1.0697 | 1.0240 |
| 2401 | Phospho-HSP27 (S82) | 0.9335 | 0.7848 | 1.0691 | 0.9064 |
| 2441 | Phospho-eIF4G (S1108) | 1.0653 | 0.7458 | 1.0709 | 1.0442 |
| 2541 | Phospho-Paxillin (Tyr118) | 1.0491 | 0.7536 | 0.9252 | 1.0110 |
| 2602 | PAK1 | 1.0976 | 0.7820 | 0.8911 | 0.9696 |
| 2605 | Phospho-PAK1 (S199/204)/PAK2 (S192/197) | 1.0681 | 0.5239 | 1.2403 | 1.1605 |
| 2741 | Phospho-eIF4E (S209) | 1.0986 | 0.8465 | 0.9852 | 0.8699 |
| 3033 | p-NF-KB p65 (S536) | 1.1810 | 0.5202 | 1.0334 | 0.9699 |
| 3671 | Phospho-Myosin Light Chain 2 (S19) | 1.1478 | 0.7167 | 1.1235 | 1.0512 |
| 9215 | p-p38 (T180/ Y182) | 1.0632 | 0.7407 | 1.1198 | 1.1403 |
| 9221 | Phospho-ATF-2 (T71) | 1.0521 | 0.6716 | 1.1923 | 1.1929 |
| 9251 | p-SAPK/ JNK (T183/ Y185) | 1.1428 | 0.4844 | 1.0278 | 1.1788 |
| 9261 | Phospho-c-Jun (S63) II | 1.1274 | 0.6195 | 1.0416 | 1.1154 |
| 9348 | Phospho-RSK3 (T356/S360) | 1.0901 | 0.5962 | 1.0567 | 1.0539 |
| 9371 | Phospho-PKC (pan) (βII S660) | 1.0230 | 0.6853 | 1.1024 | 1.0396 |
| 9718 | p-H2AX (S139) | 1.2404 | 0.4166 | 1.1419 | 1.6558 |
| 2630 | Atg5 | 1.0822 | 1.0185 | 1.0167 | 1.0158 |
| 2867 | Hexokinase II (C64G5) | 1.1200 | 0.8616 | 1.0191 | 0.9544 |
| 3195 | E-Cadherin (24E10) | 1.0964 | 1.1564 | 1.1216 | 1.0026 |
| 3205 | Pyruvate Dehydrogenase (C54G1) | 1.7156 | 0.8150 | 0.8541 | 1.0009 |
| 3810 | Enoalse-1 | 1.1969 | 0.8096 | 0.9065 | 0.9579 |
| 3820 | PDHK1 (C47H1) | 1.4547 | 0.5665 | 0.9524 | 1.0972 |
| 3879 | Snail (C15D3) | 1.2008 | 0.6185 | 1.1441 | 0.9638 |
| 4448 | AFP (D12C1) | 1.1541 | 0.5889 | 1.5694 | 1.3428 |
| 8164 | PFKP (D4B2) | 1.1763 | 0.7189 | 0.8668 | 0.9195 |
| 8171 | Enoalse-2 (D20H2) | 1.1322 | 1.0338 | 1.0006 | 0.9988 |
| 8175 | PFKL | 1.2417 | 0.8480 | 0.9862 | 1.0844 |
| 9010 | BRCA1 Antibody | 1.1554 | 0.6585 | 0.9936 | 1.0664 |
| 2971 | Phospho-mTOR (S2448) | 1.1896 | 0.5654 | 0.9373 | 1.0883 |
| 3415 | Atg3 | 1.1896 | 0.5089 | 0.9957 | 1.1645 |
| 3495 | Beclin-1 (D40C5) | 1.0562 | 0.9538 | 1.0388 | 1.0281 |
| 4407 | Phospho-EGFR (Y1173) | 1.2108 | 0.0000 | 1.2491 | 1.0472 |
| 5284 | p-BAD (S112) | 1.0416 | 0.7167 | 0.9366 | 1.0253 |
| 8060 | Aldolase A (D73H4) | 1.0265 | 0.8264 | 1.0213 | 0.9858 |
| 9258 | JNK2 (56G8) | 1.0479 | 0.8363 | 0.9814 | 0.9928 |
| 9585 | Slug (C19G7) Rabbit | 0.9928 | 1.2215 | 0.8450 | 0.9657 |
| 12434 | Keratin 17/19 (D4G2) XP® | 1.3207 | 0.0438 | 1.6621 | 0.8736 |
| 13987 | c-Myc (D3N8F) | 1.3034 | 0.4659 | 0.8393 | 1.0409 |
| 36746 | EpCAM (D4K8R) XP® | 1.3104 | 0.0765 | 1.4454 | 1.0412 |
| 59568 | MARCKSL1 (D4I9P) | 0.9850 | 1.3024 | 0.9000 | 0.9678 |
| 2101 | Phospho-Src Family (Y416) | 1.4696 | 0.0030 | 1.0179 | 1.0165 |
| 3169 | PDGF Receptor b (28E1) | 1.3369 | 0.1874 | 1.2342 | 1.1671 |
| 3176 | Progesterone Receptor A/B | 1.2839 | 0.2277 | 1.1544 | 1.1577 |
| 3674 | Phospho-Myosin Light Chain 2 (T18/S19) | 1.1085 | 0.2949 | 1.4576 | 1.3337 |
| 4377 | Phospho-p44/42 MAPK (Erk1/2) (T202/Y204) | 1.3354 | 0.1189 | 1.4134 | 1.1177 |
| 9323 | Phospho-GSK-3β (S9) (5B3) | 1.2176 | 0.1717 | 2.1771 | 1.6669 |
| 13684 | PD-L1 (E1L3N) XP | 1.3004 | 0.1909 | 1.2091 | 1.2250 |
| 2231 | Phospho-EGFR(Y845) | 1.4448 | 0.2076 | 1.0575 | 1.2151 |
| 2947 | p21 Waf1/Cip1 (12D1) | 1.0469 | 0.7991 | 1.1961 | 1.0399 |
| 3011 | PI3 Kinase p110b (C33D4) | 1.3909 | 0.1258 | 1.0990 | 1.3204 |
| 3063 | Akt2 (D6G4) | 0.9961 | 1.0224 | 1.2934 | 1.0009 |
| 3190 | PKM1/2 (C103A3) | 1.0569 | 1.0593 | 1.1346 | 0.8649 |
| 3234 | Phospho-Gab1 (Y307) | 1.0920 | 0.3083 | 0.8315 | 1.1448 |
| 3356 | Phospho-CaMKII (Y231) | 1.1204 | 0.2164 | 1.1203 | 1.0718 |
| 3361 | Phospho-CaMKII (T286) | 1.2289 | 0.0000 | 1.7042 | 1.4056 |
| 9541 | Cleaved PARP (D214) | 1.0656 | 0.7580 | 1.0974 | 1.2858 |
| 13008 | Phospho-YAP (S127) (D9W2I) | 1.2948 | 0.9911 | 1.5236 | 1.3689 |

**Table S4. In vitro AMPK activity.** Purified human AMPKα1β1γ1 complex was used to examine whether BKIDC-1553 and 1553-N-Me could be direct activators of AMPK. A well-established allosteric activator A-769662 caused 2-fold increase in AMPK activity while BKIDCs failed to do so, suggesting BKIDCs do not stimulate AMPK activity directly.

| **enzyme concentration (nM)** | **Percentage ATP consumption** | | | | | |
| --- | --- | --- | --- | --- | --- | --- |
|  | **DMSO** | **1553** | **1553-N-Me** | **1817** | **A769662** | **Compound C** |
| 40 | 52 | 56 | 60 | 55 | 91 | 0 |
| 25 | 35 | 38 | 41 | 44 | 73 | 4 |
| 15 | 19 | 20 | 23 | 24 | 23 | 3 |
| 10 | 10 | 11 | 14 | 14 | 26 | 0 |

**Table S5. BKIDC-1553 intratumoral concentrations after at least 4 weeks of thrice weekly oral dosing at 20 mg/kg.**

|  | **BKIDC-1553 tumor concentrations (µM)** | |
| --- | --- | --- |
|  | **LuCaP35** | **PC3** |
|  | 11.01 | 7.82 |
|  | 8.20 | 10.19 |
|  | 5.37 | 3.71 |
|  | 4.45 | 7.75 |
|  | 12.71 | 6.16 |
|  | 9.18 | 6.81 |
|  | 9.82 | 12.26 |
|  | 8.95 | 7.08 |
|  |  | 10.44 |
|  |  | 7.33 |
| Mean | 8.71 | 7.95 |
| Standard Deviation | 2.73 | 2.44 |
|  | **All samples were collected 48 hrs after final dose, dosing was 20 mg/kg on Mon, Weds, Fri for at least 4 weeks.* | |

**Table S6. Pharmacokinetic parameters after a single dose in mouse, rat, dog, and cynomolgus monkey.**

| **BKIDC-1553 PK Demonstrates Scalable Bioavailability Across Species** | | | | | | | |
| --- | --- | --- | --- | --- | --- | --- | --- |
| **Species & route** | **Mouse PO** | **Rat IV** | **Rat PO** | **Dog IV** | **Dog PO** | **Monkey IV** | **Monkey PO** |
| **Dose (mg/kg)** | 10 | 5 | 20 | 1 | 1 | 1 | 1 |
| **C_max_ (µM)** | 12.8 | 65.7 | 29.4 | 2.52 | 1.58 | 2.45 | 1.08 |
| **T_max_ (min)** | 320 |  | 480 |  | 60 |  | 132 |
| **AUC_0-24 hours_ (min•µmol/L)** | 13,680 | 8734 | 31,902 | 1,229 | 1,101 | 981 | 656 |
| **AUC_0-inf_ (min•µmol/L)** |  | 10,151 | 45,844 | 1,719 | 1,513 | 1,329 | 883 |
| **Clearance (mL/min/kg)** |  | 1.4 |  | 1.5 |  | 2.2 |  |
| **Vz (L/kg)** |  | 1.0 |  | 1.7 |  | 1.8 |  |
| **%F (bioavailability)** |  | ~100 |  | 88 |  | 67 |  |

Note: PO = oral. IV = intravenous.

Table S7. Rat cardiovascular assessment (nonGLP) after IV administration of increasing dosages of BKIDC-1553

| Dose (mg/kg) | Plasma Concentration BKIDC-1553 ± SEM (µM) | MAP (%) | HR (%) | dP/dt@50 (%) |
| --- | --- | --- | --- | --- |
| 3 | 3.54 ± 0.57 | 5 | 0 | 6 |
| 10 | 14.5 ± 0.69 | 5 | 0 | 10 |
| 30 | 54.5 ± 14.98 | 7 | -2 | 25 |

Note: MAP, HR, and contractility (dP/dt@50) data were percent change relative to control. For MAP and HR measurement, >10% is considered significant, whereas for contractility (dP/dt@50) >15% is considered significant.

**Uncropped Western blot images**

**Figure 3B**
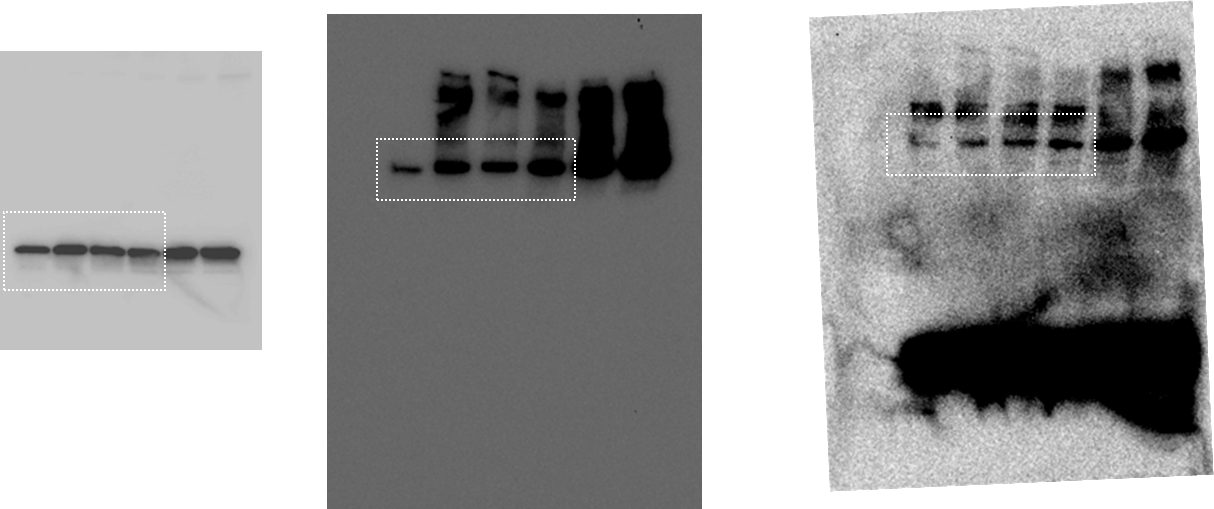

**Figure 5B**
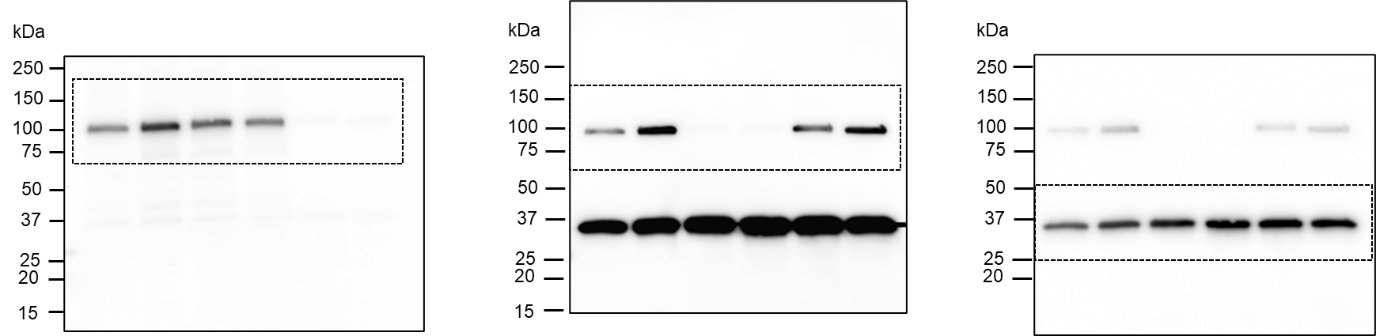

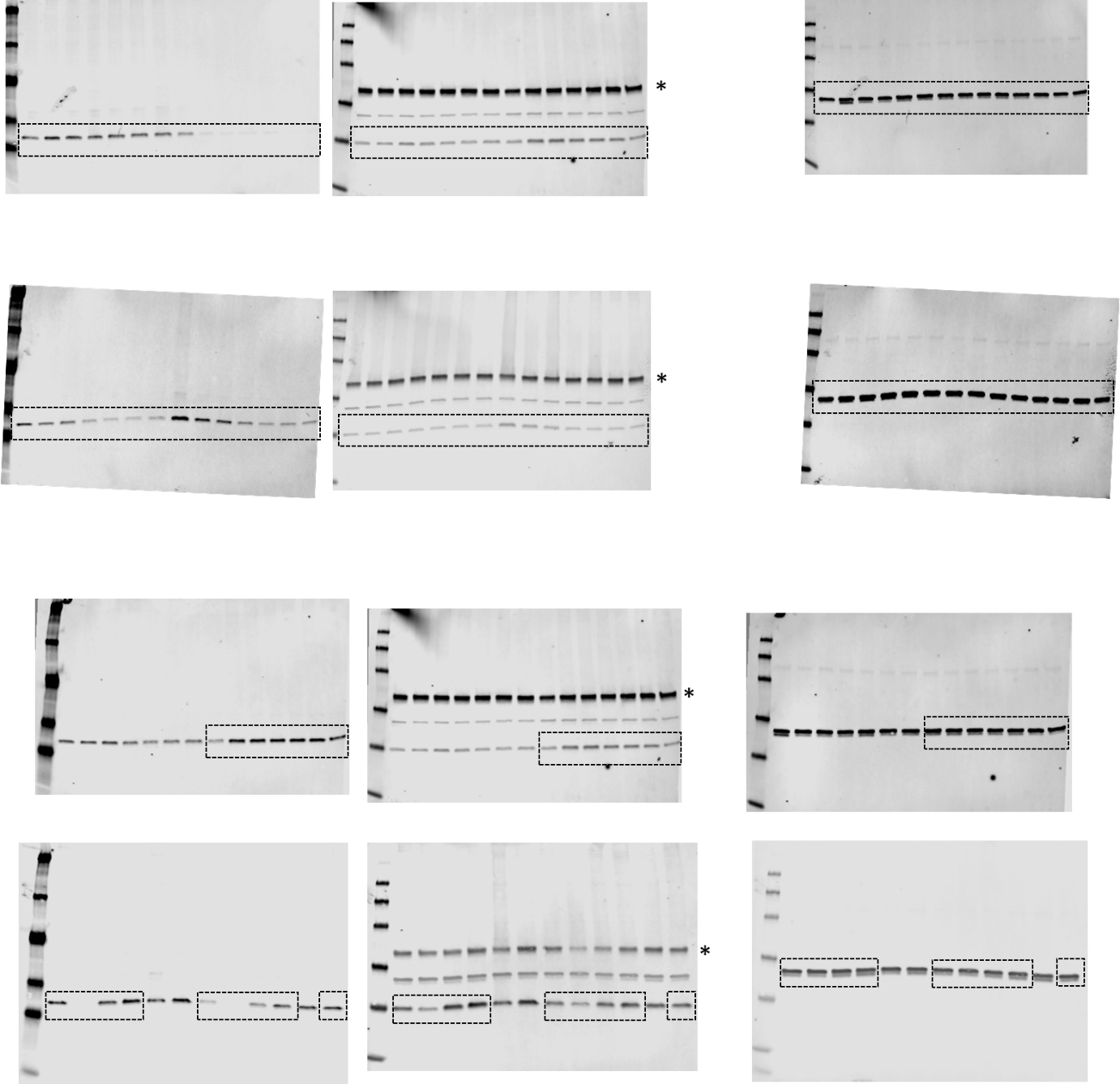
**Figure S4**

**Figure S8**

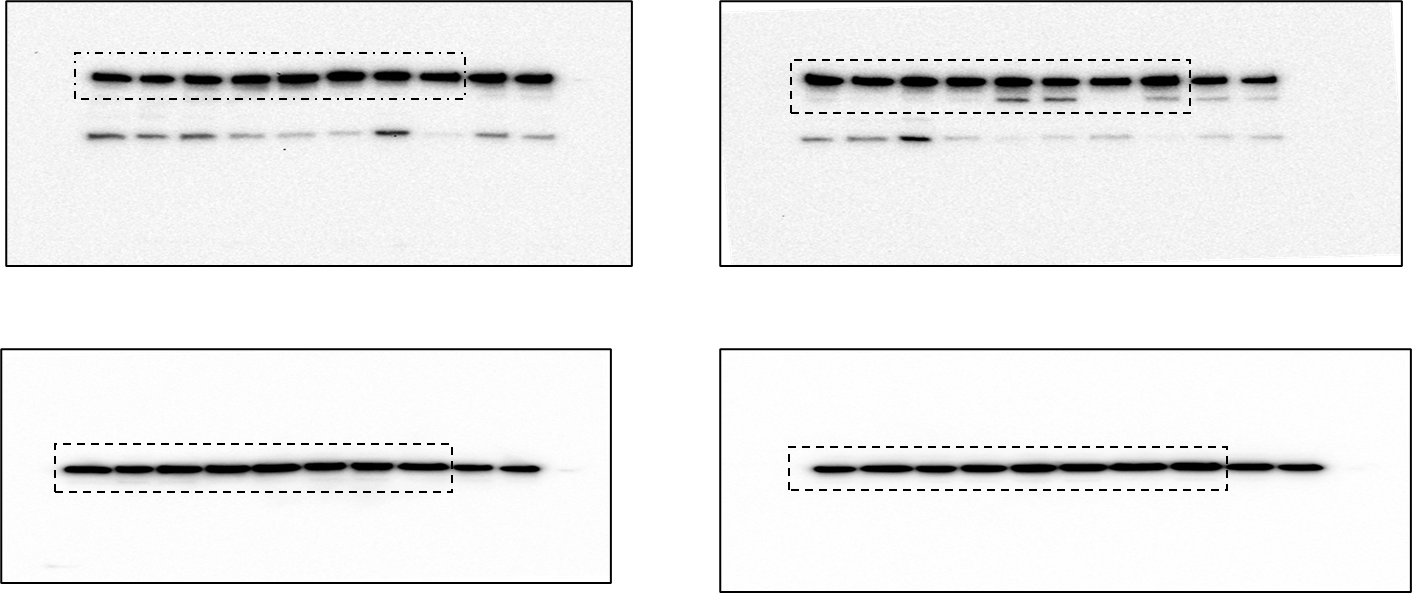

**Figure S12B**

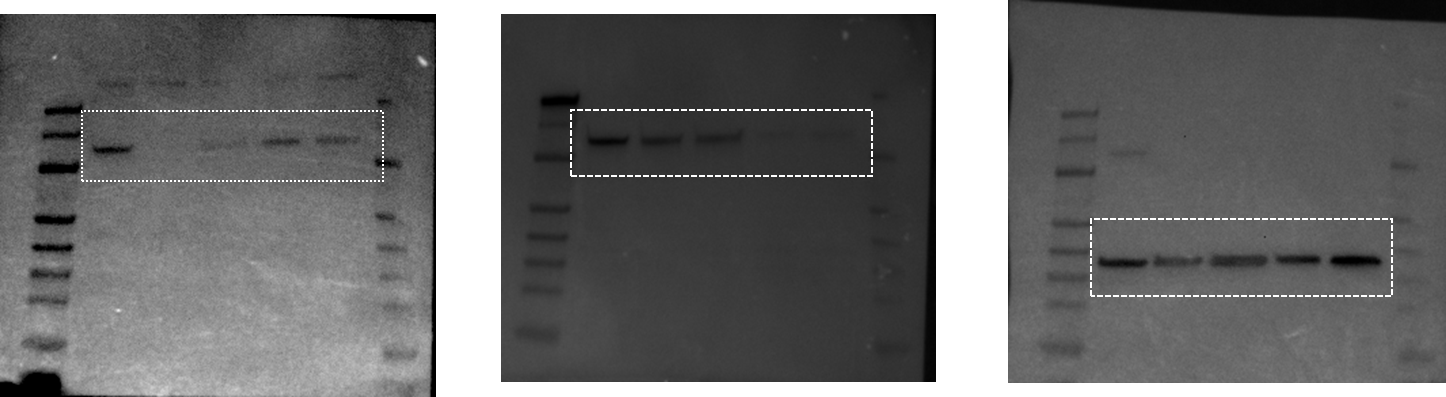

**Figure S14A**

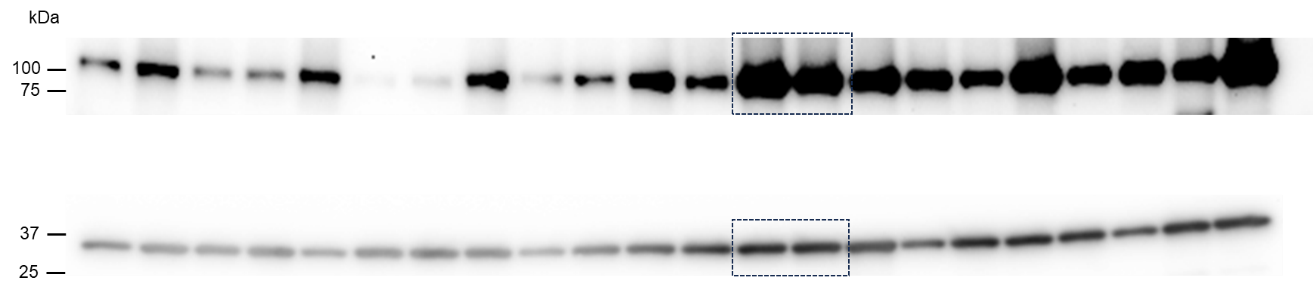
